# Experimental microevolution of *Trypanosoma cruzi* reveals hybridization and clonal mechanisms driving rapid diversification of genome sequence and structure

**DOI:** 10.1101/2021.10.24.465605

**Authors:** Gabriel M. Matos, Michael D. Lewis, Carlos Talavera-López, Matthew Yeo, Edmundo C. Grisard, Louisa A. Messenger, Michael A. Miles, Björn Andersson

## Abstract

Protozoa and fungi are known to have extraordinarily diverse mechanisms of genetic exchange. However, the presence and epidemiological relevance of genetic exchange in *Trypanosoma cruzi*, the agent of Chagas disease, has been controversial and debated for many years. Field studies have identified both predominantly clonal and sexually recombining natural populations. Two of six natural *T. cruzi* lineages (TcV and TcVI) show hybrid mosaicism, using analysis of single-gene locus markers. The formation of hybrid strains *in vitro* has been achieved and this provides a framework to study the mechanisms and adaptive significance of genetic exchange. Using whole genome sequencing of a set of experimental hybrids strains, we have confirmed that hybrid formation initially results in tetraploid parasites. The hybrid progeny showed novel mutations that were not attributable to either (diploid) parent showing an increase in amino acid changes. In long-term culture, up to 800 generations, there was progressive, gradual erosion of progeny genomes towards triploidy, yet retention of elevated copy number was observed at several core housekeeping loci. Our findings indicate hybrid formation by fusion of diploid *T. cruzi*, followed by sporadic genome erosion, but with substantial potential for adaptive evolution, as has been described as a genetic feature of other organisms, such as some fungi.

## Introduction

*Trypanosoma cruzi* is a kinetoplastid protozoan and the etiologic agent of Chagas disease, one of the neglected and highest impact parasitic infections in the Americas^1^. Chagas disease is estimated to cause great loss in both health-care costs and disability-adjusted life years^2^. Human migration and specific modes of transmission than the canonical vector-based have led to a spreading beyond its natural geographical boundaries, becoming a global issue^1, 3^. *T. cruzi* transmission is a zoonosis maintained by numerous species of triatomine insects and different species of mammals^4^. This parasite has a complex life cycle, where transmission to humans occurs most frequently by contamination with infected feces from triatomine vectors. To evade the immune responses the parasite displays a vast, complex repertoire of surface molecule genes involved in cell invasion and pathogenicity^5–7^. The regions of the genome coding for these molecules are composed of highly repetitive sequences, with hundreds to thousands of members of each family, as well as substantial numbers of transposable elements, microsatellite and tandem repeats^7–9^.

The reproductive mode of these parasitic protozoans has been widely debated, and both preponderate clonal evolution (PCE)^10, 11^ and sexual reproduction^12–14^ have been proposed. Many eukaryotic pathogens have essentially clonal population structures while also having non-obligate sexual cycles, which may enable them to adapt to environmental changes^15–17^. In trypanosomatids, the most well studied mating system is that of *Trypanosoma brucei*, for which a meiotic process is well supported based on the identification of a haploid parasite stage in the vector^18^ and on the patterns of allele inheritance and recombination observed in experimental hybrids^19^. In *T. cruzi,* great genetic and phenotypic diversity is observed and six distinct genetic clades have been recognized, named TcI to TcVI (Discrete Typing Units or DTU-I to VI)^20^. Genotyping analysis indicates that TcV and TcVI are recent, natural inter-lineage hybrids of TcII and TcIII, showing that recombination events have shaped the evolution of *T. cruzi* lineages ^13, 16, 21^. In addition to natural evidence of hybridization, the formation of *T. cruzi* hybrids *in vitro* was also described^22^. Genetic marker analyses of hybrid strains showed multi-allelic (non-Mendelian) inheritance of microsatellite alleles and an elevated DNA content^22, 23^. The hybridization events in *T. cruzi* were proposed to occur via a parasexual mechanism, similar to thise observed in certain fungi^16, 17, 22, 24^. While certain aspects of the molecular mechanisms involved in this process have been studied^25^, the contribution of each parental strain in hybrid genomes, as well as the mechanisms and adaptive significance of the hybridization phenomenon, are not well understood.

Experimental evolution approaches in defined environments have been widely used to study the evolution of model organisms (such as bacteria, yeast and *Drosophila* sp.)^26–29^ although this is not the case for parasitic protozoa. To investigate the effects of clonal reproduction and genetic exchange on the *T. cruzi* genome at the microevolutionary scale we selected two closely related TcI strains (P1, P2) and three hybrid clones that were generated from a P1 x P2 cross (1C2, 1D12, 2C1)^22^. In addition, to dissect the underlying mechanisms of hybridization in *T. cruzi* we applied a comparative genomics approach based on genome sequence data generated for parental strains and hybrid clones at the beginning and at the end of the *in vitro* microevolution experiment.

## Results

### Stability of DNA content during experimental *in vitro* microevolution

To investigate the effects of clonal reproduction and genetic exchange on the *T. cruzi* genome at the microevolutionary scale we selected two TcI strains (P1, P2) and three hybrid clones that were generated from a P1 x P2 cross (1C2, 1D12, 2C1)^22^. The two parents and three hybrids were included in an *in vitro* evolution experiment in which they were continuously cultured as epimastigotes forms for five years, equivalent to approximately 800 generations of replication by binary fission (Fig. 1a). At the end of the experiment, we re-cloned the parasite lines and selected three clones per line for analysis. During this microevolution experiment, we monitored the DNA content of each parasite line at approximately 200-generation intervals to check for evidence of large-scale structural genomic changes. We observed that the DNA content of the two parental lines remained approximately constant (Fig. 1b-1d). At the start of the experiment, the total DNA content of the hybrid clones was approximately 70% greater than the parentals (Fig. 1c-1d,^23^), but the mechanism underlying this difference has remained unknown. Over the course of the microevolution experiment, in contrast to P1 and P2, we found that the DNA content of the hybrids became progressively lower over time, up to a maximum decrease of 22.5% by generation 800. By extrapolation from the estimated genome sizes of P1 (94.5Mb) and P2 (92.5 Mb)^23^, these data indicate genome erosion in the hybrids occurred at an average rate of 23 kb per generation. Next, we looked for evidence of instability at specific loci in the hybrid genomes using PCR-based multilocus microsatellite genotyping, which had previously provided some evidence for non-Mendelian inheritance in the hybrids from the original P1 x P2 cross^22^. Specifically, at many loci the hybrids inherited more than one allele per parent. We found that three of the nine evolved hybrid clones had lost at least one microsatellite allele (Supplementary Table 1). These data indicated that the enlarged genomes of *T. cruzi* hybrids were subject to gradual erosion over an extended period of clonal replication. To dissect the underlying mechanisms, we applied a comparative genomics approach based on sequence data for parental and hybrid samples from the beginning and end of the microevolution experiment.

**Figure 1.**
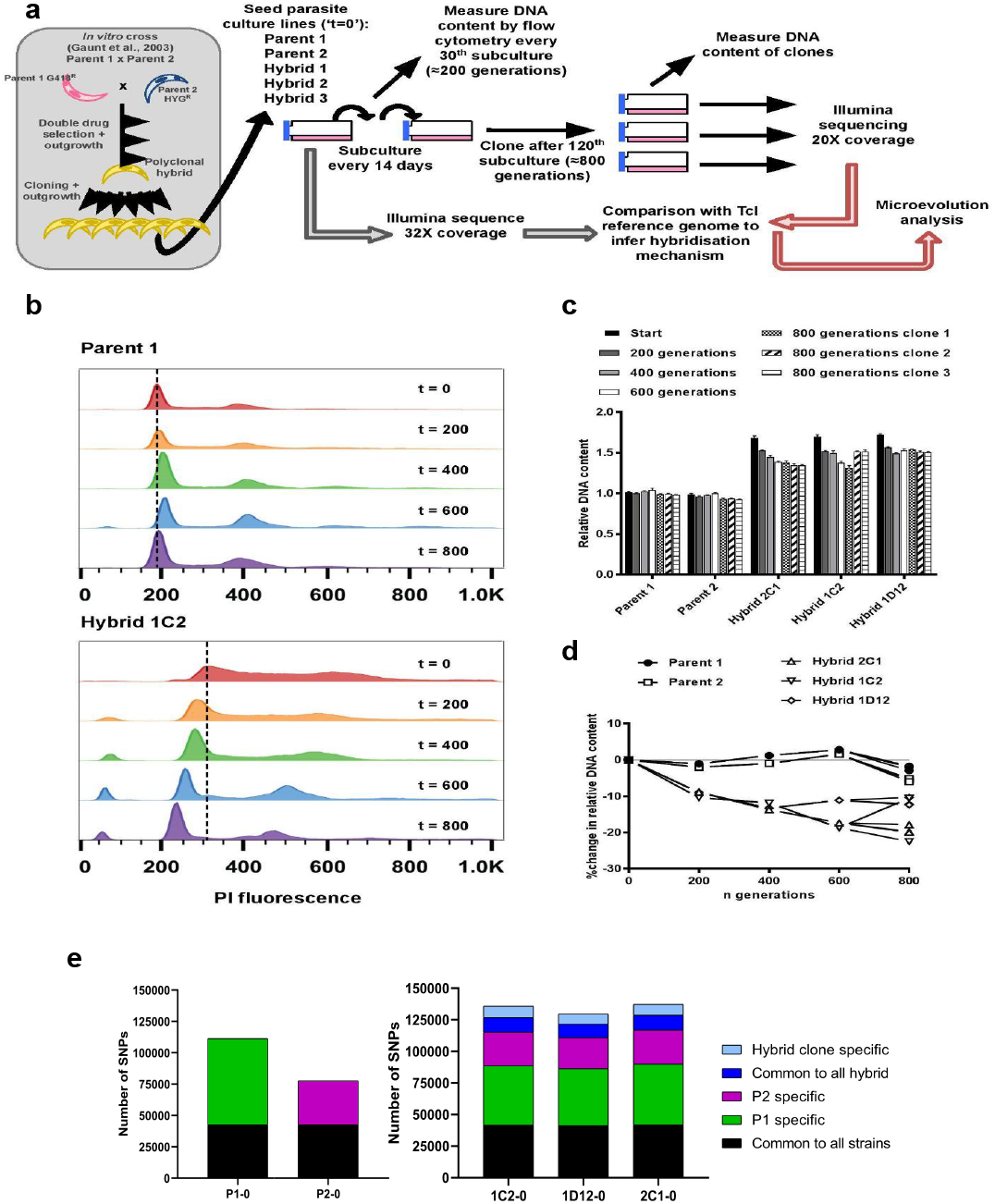
*In vitro* microevolution of *T. cruzi* hybrids **a.** Overview of experimental design. **b.** DNA content profiles shown as propidium iodide (PI) staining intensity histograms measured by flow cytometry. PI fluoresence = relative fluorescence units; *y*- axis shows number of events from the population of parasites at the indicated generation of the *in vitro* culture experiment from start (t = 0) to finish (t = 800). Plots of one parent and one hybrid are shown, representative of three biological replicates for each time point. **c.** Quantitative analysis of DNA content for both parents and three hybrid strains at each time point and for three clones per strain generated after the final time point of 800 generations. Data are the mean + SEM of three biological replicates after normalisation against an internal standard and conversion to a ratio of the mean of the two parents at t=0. **d.** The same data as shown in C, represented as the % change in DNA content over time compared to t=0. **e.** Distinct genomic signatures exclusive to each parental strain were inherited by the hybrid strains. In black are represented SNPs present in both parental strains, in green SNPs exclusive to the P1 strain, in purple SNPs exclusive to the P2 strain, in dark blue SNPs common to all hybrids and in light blue SNPs exclusive to each hybrid clone.

### Parental strains display distinct genomic signatures

The genomes of the two parental strains were characterized using *de novo* genome assemblies from short Illumina reads. Despite high sequence coverage and the linking information provided by additional long insert size libraries, the final assemblies only reconstructed 77.1 % and 74.8 % of the P1 and P2 genomes, respectively (Supplementary Table 2), due to the high repeat content of the genomes as also observed to other previously sequenced *T. cruzi* genomes. The haploid estimated genome size of P1 differed from P2 by approximately 2 Mbp, and both parental strains genomes were found to be slightly larger than the 44 Mbp reference TcI-Sylvio X10/1 strain^7^, in line with flow cytometry-based DNA content measurements^23^.

While the parental strains both belong to the TcI clade, with highly conserved synteny in the core regions, they still show significant genetic diversity. Reads from the parental strains were mapped to the reference TcI-Sylvio X10/1 genome, followed by SNV calling and a comparative analysis. The resultant genome-wide average SNP density was 9.28 SNP/kb and 3.71 SNP/kb for P1 and P2, respectively (Table 1). The majority of SNPs were located within repetitive regions, particularly in areas containing surface molecule gene family members, which can be related to gene expansion in these regions. Indeed, CNV analysis showed that P1 displays these tandem-repeated and surface molecule coding regions expanded in comparison to P2. To avoid the influence of low mapping quality in repeated regions, we applied a strict mapping quality filter, removing any SNP in regions with mapping quality below 50. SNP density in the retained high-quality regions was 2.84 and 2.11 SNPs/kb for P1 and P2, respectively (Table 1), and a total of 68,616 and 34,843 SNPs exclusive for each parental strain, were catalogued for use in the analysis of the hybrid strains (Fig. 1e). This level of differentiation between P1 and P2 provided ample scope to explore the genetic composition and inheritance patterns in the hybrid clones.

**Table 1.** Average SNP density per kilobase in starting generation parental strains and hybrid clones.

|  | SNP/kb<br>genome-wide | SNP/kb<br>high-quality mapping regions |
| --- | --- | --- |
| **P1-0** | 9.28 | 2.84 |
| **P2-0** | 3.71 | 2.11 |
| **1C2-0** | 7.63 | 3.48 |
| **1D12-0** | 7.46 | 3.29 |
| **2C1-0** | 8.22 | 3.49 |

### An increase in genetic diversity is observed after hybridization

To evaluate the impact of hybridization on genetic diversity we compared variant densities in the high-quality mapping regions of the genomes of all hybrid and parental strains. The average SNP density in these regions was higher in all hybrids than in parental strains: 3.48 SNP/kb for 1C2-0, 3.29 SNP/kb for 1D12-0 and 3.49 SNP/kb for 2C1-0 (Table 1), partially due to the presence of alleles from both parental strains. In addition to the SNPs inherited from each parental strain, additional SNPs common to all hybrids, and SNPs exclusive to each hybrid clone were identified in both coding and non-coding regions in patterns specific to each sample (Fig. 1e). The SNPs that were common to all hybrid strains are to a smaller extent SNPs that were not detected in the parental genomes, but mainly mutations that occurred in culture during the approximately 50 generations of growth before hybrid formation. Interestingly, *de novo* mutations common to all hybrids and exclusive to each hybrid clone were predicted to generate a higher percentage of non-synonymous changes in comparison to the mutations inherited from the parental strains (Supplementary Table 3). Indeed, the ratio of non-synonymous changes per synonymous changes was higher in the *de novo* mutations common to all hybrids (2.36 for 1C2-0, 2.25 for 1D12-0 and 2.19 for 2C1-0), and exclusive to each hybrid clone (2.40 for 1C2-0, 2.23 for 1D12-0 and 2.37 for 2C1-0) than in those inherited from the parental strains (1.41 for 1C2-0, 1.40 for 1D12-0 and 1.42 for 2C1-0) (Supplementary Table 3).

To investigate the impact of these mutations on the variable regions of the genome we compared the ratio of SNPs in surface molecule genes per non-surface molecule genes. This ratio is approximately 4-fold higher in SNPs common to all hybrids and in SNPs exclusive to each hybrid clone than in those inherited from the parental strains (Supplementary Table 4). In addition, we compared the ratio of non-synonymous mutations in surface molecule genes per non-synonymous mutations in other genes. Interestingly, this ratio was approximately 3.3-fold higher in the SNPs that appeared after the hybridization event in comparison to those inherited from the parental strains (Supplementary Table 5).

### Chromosome copy number variation analysis reveals tetraploid hybrids

Chromosome copy number variation (CCNV) was determined using the combination of Read Depth Coverage (RDC) and Allele Balance (AB) methods^30^. An increase or decrease in the mean RDC of a chromosome when compared to the overall genome coverage is an indicator of a gain or loss of chromosomal sequences, respectively^30^. If the ratio between the median chromosome coverage and the median genome coverage is approximately one, the chromosome has the same copy number as the genome, while fluctuations in this ratio indicates aneuploidies. Here, the RDC analysis was based on the ratio between the mean coverage of predicted single-copy genes in each chromosome and the mean coverage of all single-copy genes in the genome as described by^31^ (Fig. 2a). This methodology was shown to eliminate bias caused by chromosomal repetitive content in *T. cruzi*^31^, which would otherwise obscure the correct copy numbers.

**Figure 2.**
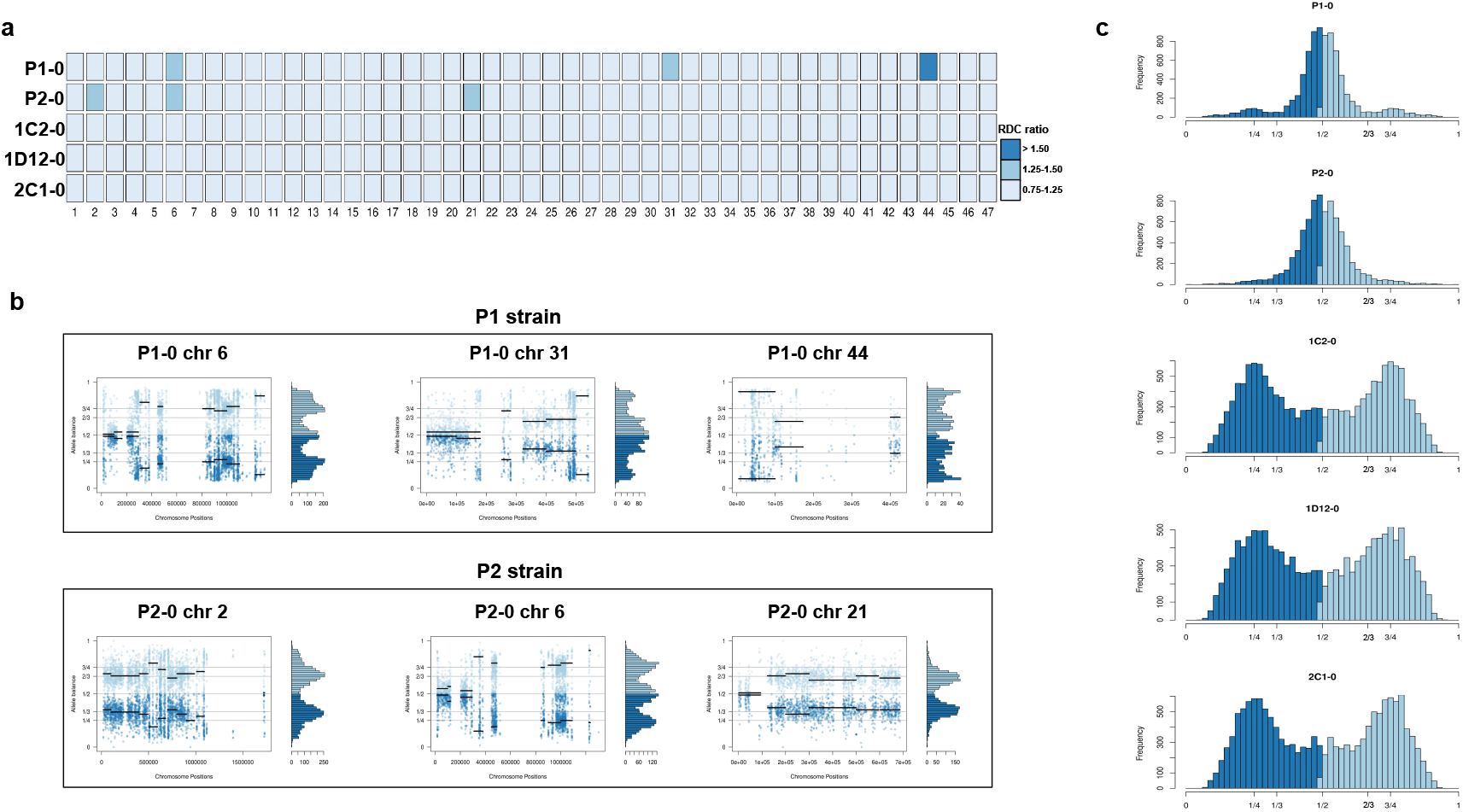
- Somy estimation based on Read Depth Coverage (RDC) and Allelic Balance (AB) reveals aneuploidies in parental strains and tetrasomic hybrid clones. **a.** Aneuploidy analysis based on RDC among all 47 chromosomes in the starting generation of parental strains (P1-0 and P2-0) and hybrids (1C2-0, 1D12-0 and 2C1-0). An increase or decrease in the mean RDC of a chromosome when compared to the genome coverage is an indicator of a gain or loss of chromosomal sequences. When the ratio between the median chromosome coverage and the median genome coverage was approximately one, the chromosome was considered having the same somy as the genome, while fluctuations in this ratio (lower than 0.75 or higher than 1.25) were considered as putative aneuploidies. **b.** Proportion of the alleles in heterozygous SNP positions in chromosomes with RDC ratio higher than 1.25 to confirm somy estimations. Blue points represent the proportion of the alleles in heterozygous SNP positions along the chromosome. Darker blue represents the frequency of the first allele while lighter blue represents the frequency of the second allele, black lines represent the median value in windows. **c.** Ploidy estimation based on the proportion of the alleles in heterozygous SNP positions of single-copy genes. It is expected that diploid genomes display a proportion of each allele around 50%, triploid genomes around 33.3% and 66.6%, while tetraploid genomes around 25% and 75% or 50%^29^. Darker blue represents the frequency of the first allele while lighter blue represents the frequency of the second allele.

We were unable to estimate CCNV in chromosomes 17, 22, 30 and 47 by this methodology due to the lack of single-copy genes in these chromosomes, and therefore the chromosomal somy prediction presented for them was estimated based on the ratio between the mean RDC of each chromosome position and the mean coverage of all genome positions (Fig. 2a). Aneuploidy was observed in the P2 strain, involving trisomy of chromosomes 2 and 21 (Fig. 2a-2b). The RDC analysis indicated aneuploidy in chromosome 44 in the P1 strain, but due to the lack of regions with high-quality SNPs, it was not possible to confirm this in the AB analysis (Fig. 2a-2b). The RDC analysis also indicated aneuploidies in chromosome 6 in both parental strains and in chromosome 31 of P1 strain (Fig. 2a). However, the AB analysis was not consistent across the entire chromosome (Fig. 2b), which may be related to copy number variation in specific regions of these chromosomes rather than aneuploidy, as observed in other *T. cruzi* strains^13, 31^.

RDC analysis does not allow differentiation between different levels of euploidy (or near-euploidy), therefore whole genome ploidy was estimated by analysing the proportion of the alleles in heterozygous SNP positions of all single-copy genes (Fig. 2c). In both parental strains a peak at 50% was observed, while in the hybrids, peaks at 25 and 75% or 50% were observed (Fig. 2c), which is expected for diploid and tetraploid genomes, respectively^30^. This ploidy estimation is in accordance with the previous DNA content estimation based on flow cytometry data, in which the hybrids displayed ∼70% greater DNA content than the parental strains (Fig. 1c,^23^), given that this method could not determine the relative amounts of mitochondrial kDNA and nuclear DNA. Inheritance of SNPs exclusive to both parental strains was observed in the hybrid genomes (Fig. 1e), which shows that the tetrasomic profile observed resulted from a contribution from both parental chromosomes.

In order to address chromosome specificities and confirm the somy estimates, the AB analysis was performed independently for each chromosome. The proportion of each allele with heterozygous SNPs per chromosome position was plotted for the 47 TcI chromosomes (Supplementary Figs. 1-5). Clear patterns of somy were observed in 37 chromosomes (chrs 1-16, 18-19, 21, 23, 25-29, 31-33, 35-39, 41, 43, 45-46; Supplementary Figs. 1-5). Due to a lack of regions containing high-quality heterozygous SNPs generating inconsistent AB, somy estimation was not reliable for chromosomes 17, 20, 22, 24, 30, 34, 40, 42, 44, 47 (Supplementary Figs. 1-5). AB analysis shown to be reliable for estimating aneuploidy in chromosomes with longer core regions that contain housekeeping genes and other genes that are conserved between kinetoplastids, while the presence of large tandemly repeated regions and surface molecule gene families made read mapping, and thus copy number estimation difficult for some chromosomes. High resolution representations of the CCNV analysis are shown for six chromosomes (Fig. 3). Chromosomes 2, 7 and 13 have longer tandem repeated regions and coding sequences for surface molecules (percentage of surface molecule genes: chr2 7.26%, chr7 9.55%, chr13 8.38%), while chromosomes 18, 21 and 37 have longer core regions (percentage of surface molecule genes: chr18 1.25%, chr21 1.98%, chr37 3.74%) (Fig. 3a). The mapping quality in surface molecules’ coding sequences was often observed to drop significantly, which affected the inferred proportions of alleles in those regions, and this was found to lead to erroneous somy estimates (Fig. 3b). To avoid this, a strict mapping quality filter was applied, and all the allele proportions were plotted by chromosome position. Using this strategy, it was possible to identify specific regions in chromosomes that could be used to infer CCNV with high confidence. Chromosomes 2, 7 and 13 showed longer regions with diverse allele proportions, nonetheless it was still possible to identify the trisomy of chromosome 2 of P2 and tetrasomies in hybrids based on using the smaller core regions (Fig. 3c). Chromosomes 18, 21 and 37 showed somy patterns that were clearly consistent with the whole genome CCNV, confirming the trisomy in chromosome 21 of P2 and tetrasomy in the hybrid clones (Fig. 3d).

**Figure 3.**
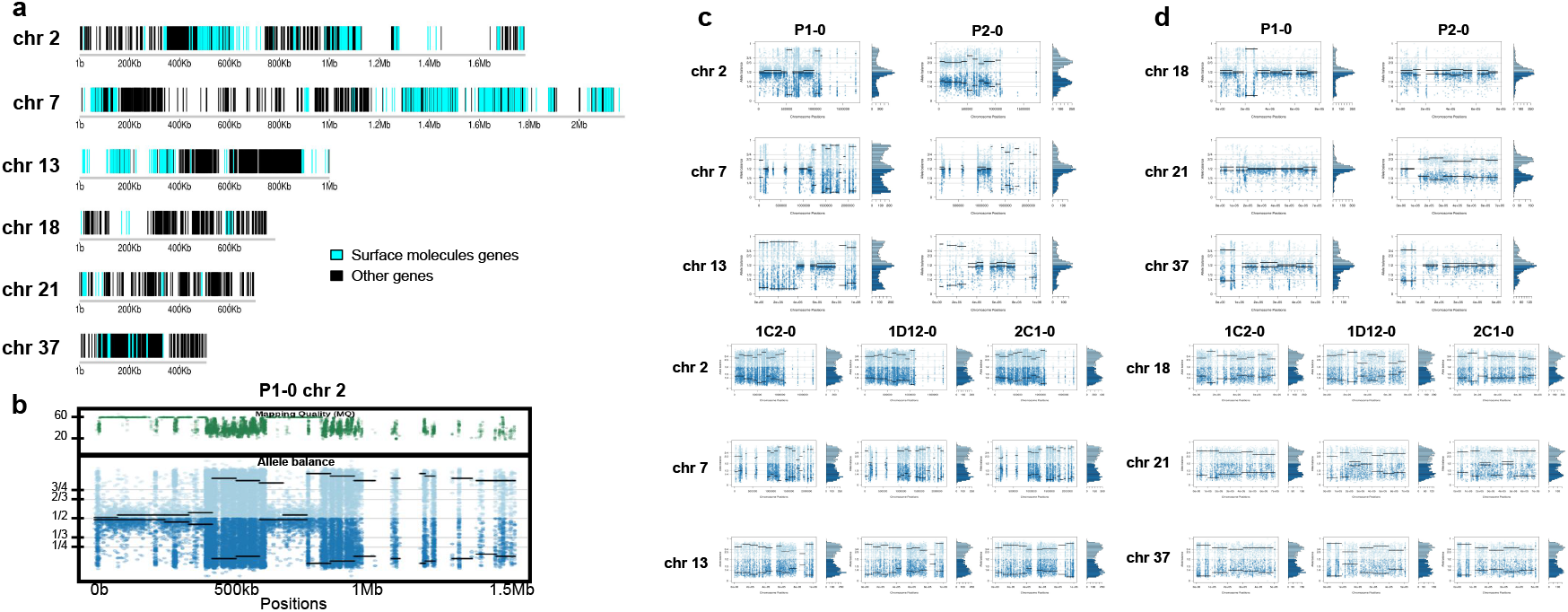
Somy estimation per chromosome based on AB in starting generation confirming aneuploidies in parental strains and tetrasomic hybrid clones. **a.** Gene distribution and its implication in mapping quality and somy estimation of six representative TcI chromosomes with longer surface molecule-encoding regions (chromosomes 2, 7 and 13) and shorter encoding surface molecule-encoding regions (chromosomes 18, 21 and 37). Surface molecule genes are represented as cyan blue boxes while other genes are represented as black boxes. **b.** Implication of chromosome gene distribution in mapping quality and somy estimation based on AB in chromosomes 2 (longer surface molecule-encoding regions) of parental strain 1 (P1-0). Dark green represents the fluctuation in mapping quality along chromosome positions. Blue points represent the proportion of the alleles in heterozygous SNP positions along the chromosome. Darker blue represents the frequency of the first allele while lighter blue represents the frequency of the second allele, black lines represent the median value in windows. **c.** Proportion of the alleles in heterozygous SNP positions in chromosomes with longer surface molecule-encoding regions. **d.** Proportion of the alleles in heterozygous SNP positions in chromosomes with shorter surface molecule-encoding regions. Blue points represent the proportion of the alleles in heterozygous SNP positions along the chromosome. Darker blue represents the frequency of the first allele while lighter blue represents the frequency of the second allele, black lines represent the median value in windows.

### Chromosomal copy number variation reveals sequential loss of chromosomal copy in hybrid clones after culture growth

CCNV and whole genome ploidy were evaluated in three replicate clones for each parental and hybrid strains after 800 generations in continuous *in vitro* culture. The trisomies in chromosomes 2 and 21 of P2 were no longer observed in any replicate clone after growth in culture (Fig. 4a). New aneuploidies were identified in the parental strains, involving trisomy of chromosomes 37 and 45 in all P1 clones and trisomy of chromosome 32 and 37 in all P2 clones (Fig. 4a-4b). As observed in the first generation, the RDC and AB analyses in chromosomes 6 and 31 were not consistent throughout the chromosomes (Supplementary Figs. 6-10). Whole genome ploidy estimation revealed a shrinking pattern in the hybrids, which was consistent with the prior flow cytometry analysis (Fig. 1c). While parental strains remained essentially diploid (apart from the few aforementioned trisomies), a mixture between trisomic and tetrasomic profiles was observed in the hybrids after growth in culture (Fig. 4c). Indeed, hybrid 2C1 displayed a clear transition from a tetraploid to a triploid pattern after growth in culture (Fig. 4c), suggesting that genome erosion eventually leads to a return towards diploidy after the hybridization event, as observed in the naturally occurring TcV and TcVI hybrids^23^. In contrast with the first generation, the RDC analysis indicates putative aneuploidies in hybrid clones after extended culture growth (Fig. 4a; Supplementary Figs. 6-10). Despite the patterns of genome erosion, tetrasomic profiles in chromosomes 19 and 37 were present in all hybrid clones after culture growth (Fig. 4A-4B). Taken together with previous DNA content analysis of intermediate time points (Fig. 1), our results from the different *in vitro* evolved clones indicate that the genome erosion was gradual, with sequential losses of chromosome copies and regions rather than coordinated jumps between ploidy levels. The proportion of each allele in heterozygous SNPs per chromosome position was also plotted for the 47 chromosomes in each replicate after culture growth (Supplementary Figs. 6-10). As observed in the representative chromosomes, the evolved parental clones still have a clear disomic pattern, while a mixture of trisomic and tetrasomic patterns is observed in evolved hybrid clones (Fig. 5; Supplementary Figs. 6-10).

**Figure 4.**
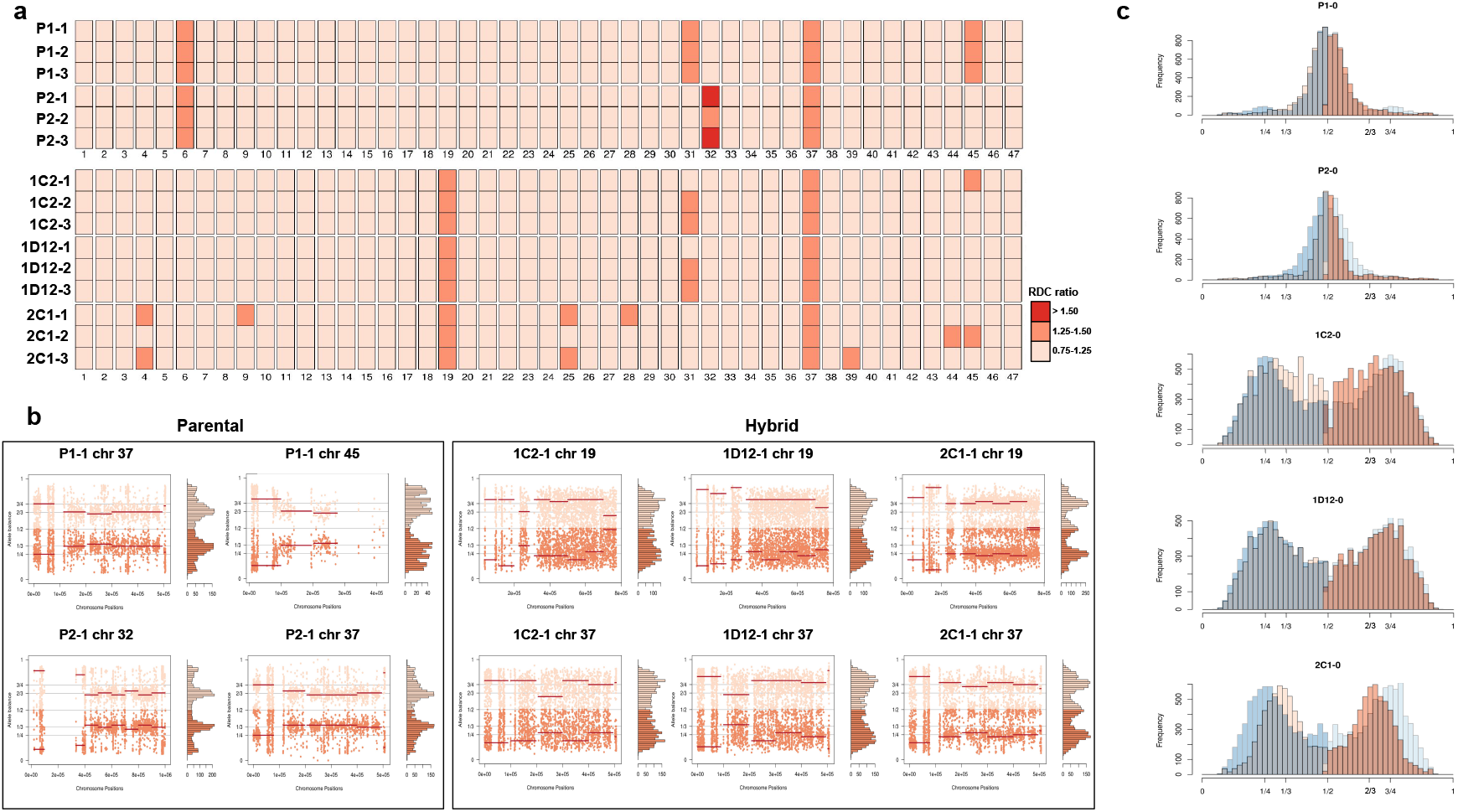
- Somy estimation based on RDC and AB reveals novel aneuploidies and a shift in hybrid strains ploidy after 800 generations in *in vitro* culture. **a.** Aneuploidy analysis based on RDC among the 47 chromosomes in all clones after culture growth. **b.** Proportion of the alleles in heterozygous SNP positions in chromosomes showing aneuploidies in all parental or hybrid clones. **c.** Comparison of somy estimation based on the proportion of the alleles in heterozygous SNP positions of single-copy genes. First generation is presented in shaded blue, while the 800 generation is presented in orange.

**Figure 5.**
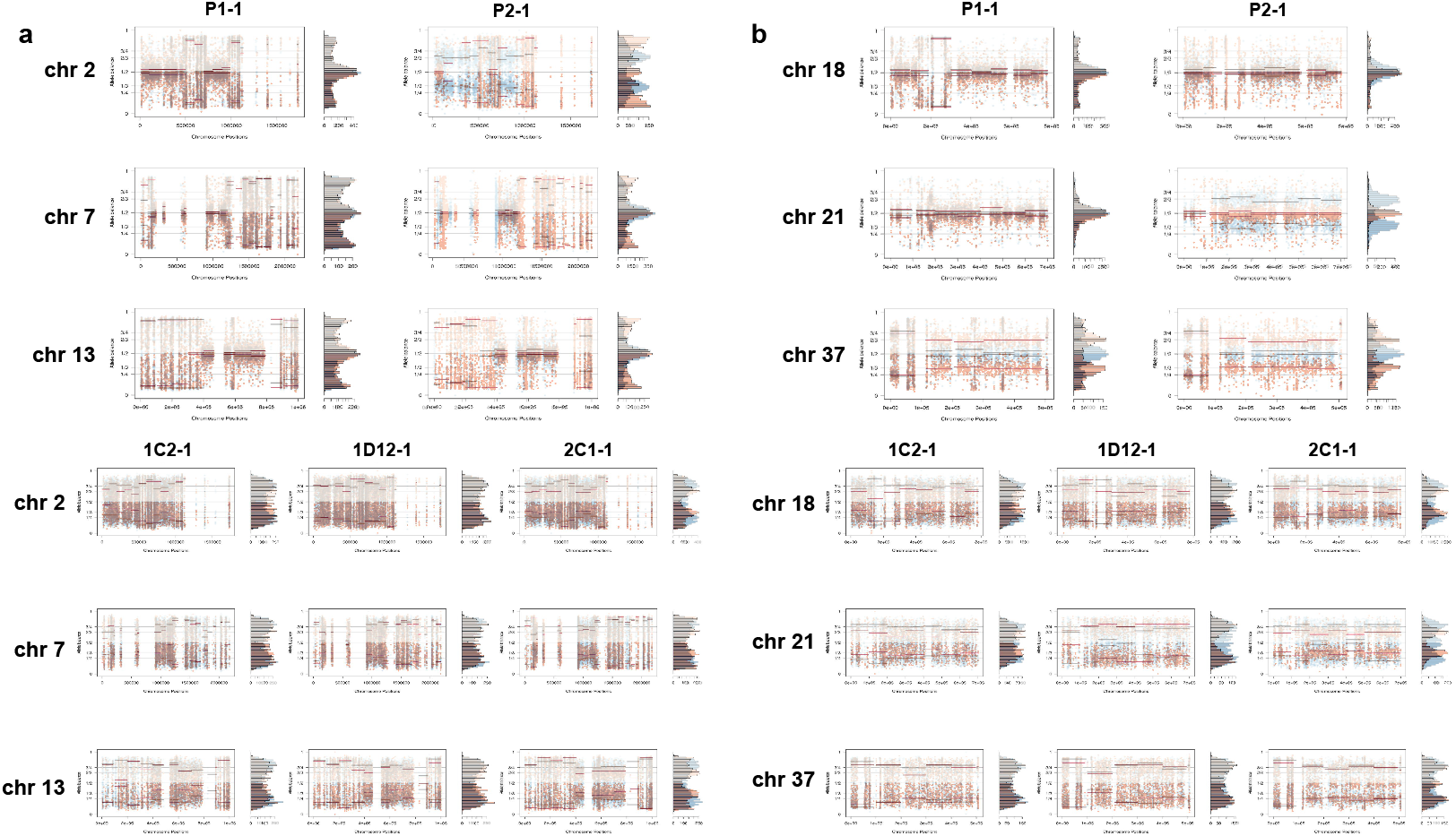
Somy comparison between first generation and 800 generation confirming novel aneuploidies after *in vitro* culture. **a.** Proportion of the alleles in heterozygous SNP positions in chromosomes with longer surface molecule-encoding regions. **b.** Proportion of the alleles in heterozygous SNP positions in chromosomes with shorter surface molecule-encoding regions. Points represent the proportion of the alleles in heterozygous SNP positions along the chromosome. 800 generation data is plotted in orange, while first generation data is plotted in shaded blue.

### Patterns of genome erosion after *in vitro* microevolution

To investigate the patterns of genome erosion and CCNV in the hybrids, we performed a copy number variation (CNV) analysis throughout the whole genome of parental and hybrid clones after extended culture growth. Despite the stability of DNA content in parental clones during our experiment (Fig. 1c-1d), similar gene categories showed copy number variation in both parental and hybrid clones after culture growth (Supplemental_Tables_S6-S9). Gene losses occurred mainly in host-parasite interaction genes such as surface molecule genes and genes related to exocytosis and cell secretion, while genes related to macromolecule metabolism, ion transport and cell division showed an increase in copy numbers (Fig. 6; Supplementary Fig. 11). Despite the events of genome erosion, chromosomes 19 and 37 remained tetrasomic in all hybrid clones and an extra copy of chromosome 37 is observed in all parental clones after culture growth (Fig. 4b and Fig. 5b). Chromosomes 19 and 37 contain long core regions with abundant housekeeping genes involved in cellular metabolism, DNA replication and transcription (Supplementary Table 10-11) which may contribute to fitness in culture. Altogether, these results suggest that the presence or absence of selective pressure from the environment may shape CCNV and genome erosion in *T. cruzi*, resulting in distinct genome patterns after hybridization events occur.

**Figure 6.**
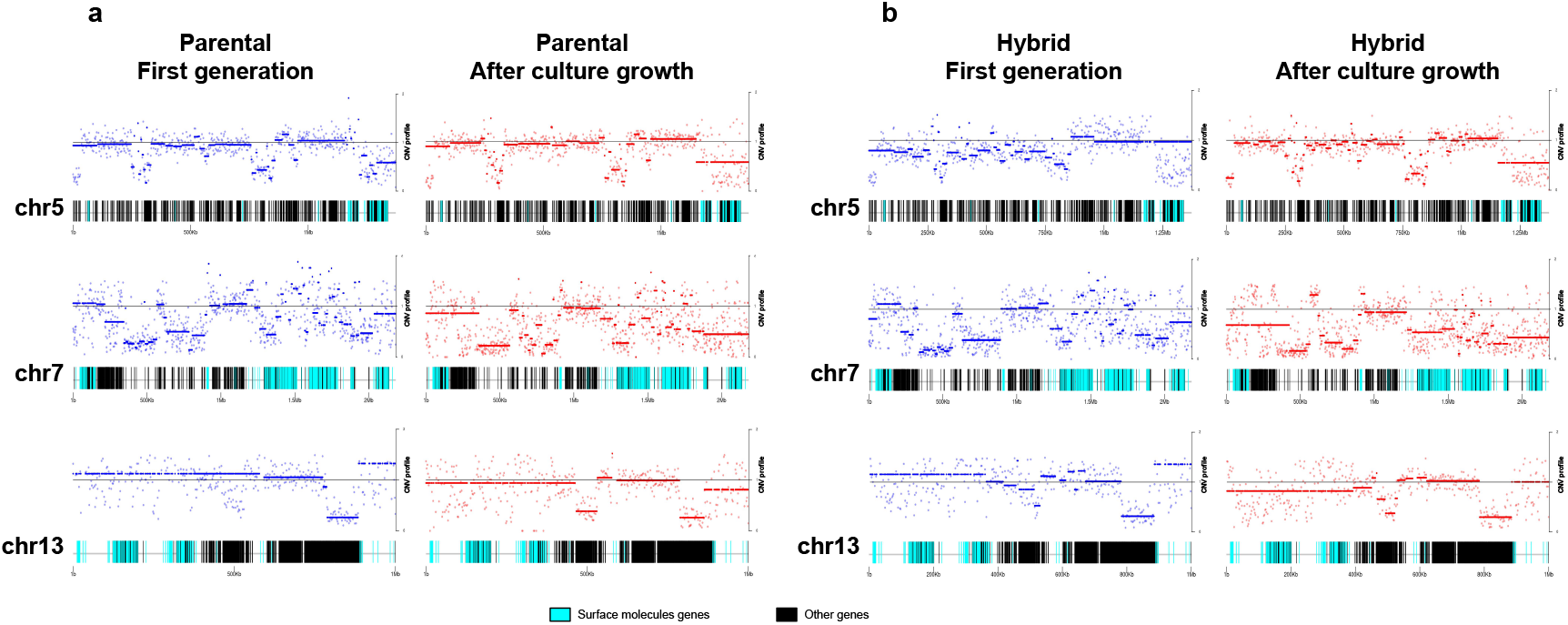
Chromosome gene composition and copy number variation (CNV) comparison indicates genome erosion patterns in surface molecule coding regions after *in vitro* culture growth. **a.** CNV comparison in parental strain. **b.** CNV comparison in hybrid strain. Each dot in the CNV profile chart represents the normalized depth per kb, while lines represent the median ratio^56^. Blue points and lines represent CNV profiles in the first generation, while red points and lines represent CNV profiles after 800 generations of *in vitro* culture growth. Surface molecule genes are represented as cyan blue boxes, while other genes are represented as black boxes.

### Surface molecule genes evolve faster than other regions of the genome

The effect of extended culture growth on genetic diversity was evaluated in all parental and hybrid evolved clones. Regarding the parental strains, all P1 replicates showed a similar SNP density in the retained high-quality regions after culture growth (P1-0: 2.84; P1-1: 2.80; P1-2: 2.77; P1-3: 2.82), while all P2 replicates showed an increase in SNP density after culture growth (P2-0: 2.11; P2-1 2.78; P2-2 2.81; P2-3 2.66) (Fig. 7a). The P2 replicates have a reduction in copies of HUS1-like encoding genes, which are involved in DNA repair, after culture growth (Supplementary Table 12). A gene enrichment analysis showed that a number of these genes were located on chromosome 2, suggesting that the loss of trisomy in this chromosome after culture growth could possibly have contributed to the higher number of mutations that was found in the P2 clones. New genomic variants were found in both evolved parental clones (Fig. 7a). All evolved hybrid replicate clones, with exception of 1D12-3, showed a slight decrease in SNP density after culture growth (1C2-0: 3.48; 1C2-1: 2.96; 1C2-2: 3.26; 1C2-3: 3.10; 1D12-0: 3.29; 1D12-1: 3.24; 1D12-2: 3.22; 1D12-3: 3.36; 2C1-0: 3.49; 2C1-1: 3.34; 2C1-2: 2.64; 2C1-3: 3.25, Fig. 7a), likely caused by the loss of genetic material. Multiple new genomic variants were found in the *in vitro* evolved hybrid progeny, displaying patterns specific to each clonal isolate (Fig. 7a). However, these variants were distinct from the novel mutations that emerged in the period between hybrid formation and the start of the microevolution experiment (*t*=0) as described above.

**Figure 7.**
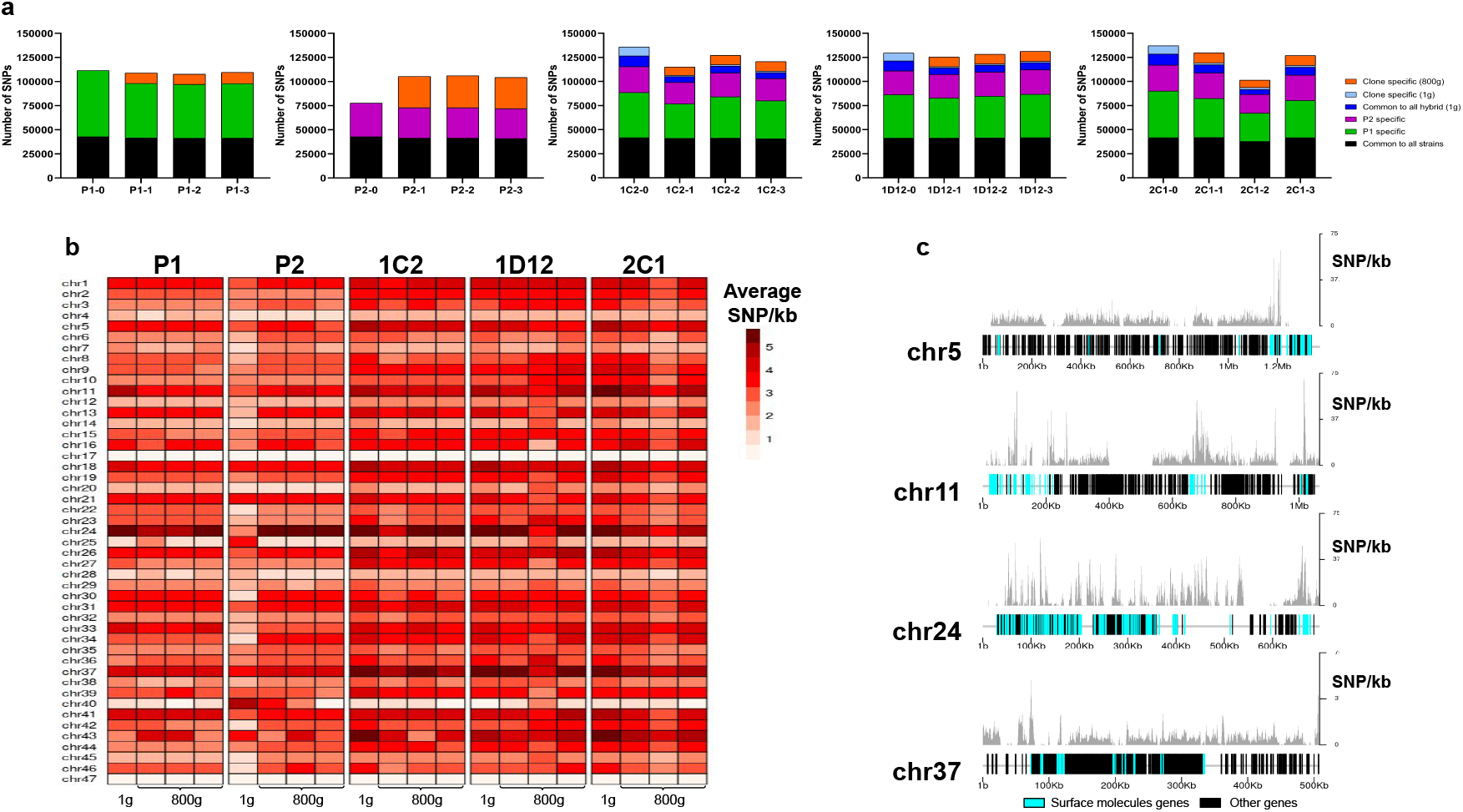
SNP density analysis throughout the genome indicates that surface molecule coding regions displayed a higher amount of variants than other regions of the genome for all strains. **a.** Number of SNPs in all clones after long-term *in vitro* culture. In black are represented SNPs common to all hybrid strains, in green SNPs specific to P1 strain, in purple SNPs specific to the P2 strain, in dark blue SNPs common to all hybrids in the first generation, in light blue SNPs exclusive to each hybrid clone in the first generation and in orange SNPs exclusive to each clone after culture growth. **b.** Chromosome average SNP density per kilobase in all samples before (1g) and after culture growth (800g). **c.** Chromosomes with higher SNP density plotted displaying gene composition and SNP density. Surface molecule genes are represented as cyan blue boxes, while other genes are represented as black boxes.

To better understand how these genomes evolved during culture growth, the SNP density was also evaluated in each chromosome separately. The SNP density pattern per chromosome remained similar for all samples in first generation (*t*=0) and after culture growth (*t*=800). Some chromosomes (e.g. chrs 5, 11, 24 and 37) display higher SNP density than the others across all samples (Fig. 7b). After culture growth, the P2 evolved clone replicates had an increase in SNP density in nearly all chromosomes (Fig. 7b). The emergence of novel SNPs after culture growth resulted in a majority of non-synonymous mutations. The ratio of non-synonymous per synonymous changes in novel SNPs was similar between both parental strains (P1-1: 2.26; P1-2: 2.22; P1-3: 2.38; P2-1: 2.25; P2- 2: 2.22; P2-3: 2.26) and hybrid clones (1C2-1: 2.20; 1C2-2: 2.48; 1C2-3: 2.26; 1D12-1: 2.41; 1D12-2: 2.5; 1D12-3: 2.40; 2C1-1: 2.21; 2C1-2: 2.18; 2C1-3: 2.30). Interestingly, the regions of the genome with higher genetic variation in all clone isolates contain surface molecule coding sequences (Fig. 7c). To investigate the impact of new mutations in these regions, we compared the ratio of non-synonymous mutations in surface molecule genes to non-synonymous mutations in other genes. This ratio was higher for the mutations that appeared in each parental clone after culture growth in comparison to those present in the first generation (approx. 2.5-fold higher for P1 clones and 3.5-fold higher for P2 clones; Supplementary Table 13). Regarding the hybrid strains, a similar ratio was observed in the SNPs that appeared in each hybrid clone after culture growth and those exclusive to each hybrid clone present in the first generation (Supplementary Table 13). These results indicate that surface molecule coding regions may mutate faster than the rest of the genome either by clonal or parasexual mechanisms, representing an important region of genomic diversification.

## Discussion

The results from whole genome sequencing analysis of this *T. cruzi* genetic cross have clearly demonstrated that, while the parental strains were diploid, all initial hybrid clones were essentially tetraploid. We showed that genomic variants exclusive to both parental strains were present in all hybrid clones, indicating that the tetraploid profile observed is obtained by the fusion of the two disomic parental genomes, although more complex pathways cannot be ruled out in the absence of direct observational evidence. In addition, we showed that genome erosion was gradual, with sequential losses of chromosome copies rather than coordinated jumps between ploidy levels. As previously proposed^16, 22^, our results confirmed that hybridization in *T. cruzi* happened via a parasexual mechanism rather than a canonical meiotic process. This phenomenon resembles the parasexual cycle of the pathogenic fungus *Candida albicans*, which also involves diploid fusion followed by non-meiotic genome reduction back to aneuploids and diploids (reviewed by^24^). In contrast, hybridization in *T. brucei* and possibly in *Leishmania* spp. has been associated with parental strains having undergone a meiosis-like process followed by a fusion of haploid cells^14, 18, 32–35^. In this process, mainly diploid hybrids are formed, but polyploid hybrids may also be produced by the fusion between parental cells that failed to undergo meiosis^14, 18, 32–35^. Despite the clear evidence of parasexual hybridization in *in vitro T. cruzi* hybrids, it is still not clear if a meiosis-like mechanism could also contribute to the generation of natural *T. cruzi* hybrids, as observed in other trypanosomatids. Further work will be required to establish whether there is a single common mechanism or several alternative modes of hybridization.

Our microevolution experiment has shown that *T. cruzi* genomes are highly responsive to the environmental conditions. The CNV analysis showed that while surface molecule gene numbers were being eroded after culture growth, genes involved in cell metabolism, cell division and ion transport were being expanded. This genome plasticity was not only observed at the level of gene copy number variation, but also at the chromosome level. The parental strains displayed distinct aneuploidies before and after culture, and despite the overall pattern of genome erosion, some chromosomes remained tetrasomic in the hybrid strains. Indeed, aneuploidies have been widely reported in natural *T. cruzi* populations, and although CCNV varies among and within DTUs, it seems constant within a given population^13, 31, 36^. While in many multicellular organisms, aneuploidy is known to have severe consequences, in trypanosomatids it represents an important mechanism for rapidly overcoming environmental diversity^36–39^. Aneuploidy is reversed quickly once the benefits of an extra chromosome expire, as observed in P2 aneuploidies in chromosomes 2 and 21. In addition, trypanosomatids may control overexpression caused by additional gene copies, at the protein level^40^. We found aneuploidy in chromosome 37 in all parental and hybrid clones after culture growth, suggesting a benefit to parasite fitness in culture. As observed in *Leishmania* spp., the fixation of genetic alterations is the result of positive selection processes that adapt parasite fitness to a given ecology or transmission cycle^38^. We can speculate that in natural *cruzi* hybrids, distinct CCNV and genome erosion patterns would be observed due to specific environmental selective pressure with the expansion of genes involved in host-parasite interactions.

Genome variant analyses showed that an increase in genetic sequence diversity was present after hybridization. Hybridization events are associated with a burst of novel mutations and a myriad of genomic rearrangements (e.g. transpositions, genome size changes, chromosomal rearrangements) in distinct eukaryotes^17, 41^. Our results indicate that the process of *T. cruzi* hybrid formation included both new mutations and most likely recombination between parental chromosomes. Here, we were able to show an increase in the number of SNPs in the hybrid strains with the accumulation of amino acid substitutions. Interestingly, the numbers of new SNPs in culture showed a clear difference between the two parental strains after 800 generations, in which P2 showed a higher mutation rate. A previous study has reported that TcI strains have a mismatch repair machinery, which repairs base misincorporation and erroneous insertions and deletions during DNA recombination and replication, that is more efficient than in other DTUs^31, 42, 43^. As described in *Leishmania major*, HUS1 was shown to be required for genome stability under non-stressed conditions and its suppression had a genome-wide mutagenic effect^44^. Since many of the genes coding for the HUS1-like proteins are encoded on chromosome 2, the loss of trisomy in this chromosome after culture growth could possibly be related to the increase in mutations observed in the P2 clones. In general, culture growth led to a reduction in genetic variants in nearly all evolved clones in comparison to the first generation. This reduction may be related to the loss of heterozygosity, but also to the loss of gene copies and genome erosion of surface molecules regions after culture growth. In addition, we could not identify any bias in the loss of a specific parental chromosome even in triploid hybrid clones, suggesting recombination between the parental chromosomes. As material from both parental strains has been lost during over 800 generations, it is likely that the hybrid clones would stabilize at diploidy and contain chromosomes from both parental strains, with a higher polymorphism than the parental strains. This is similar to strains from the known natural hybrid clades, TcV and TcVI. Strains from these clades have diploid genomes with high repeat content, and high levels of polymorphism in many regions, resulting from alleles from two divergent parental strains (from clades TcII and TcIII)^16, 21, 23^. From our analyses, it thus appears that the *in vitro* hybrid formation presented here is highly similar to the process that generated the natural hybrids.

Our microevolution experiment has shown that surface molecule genes represent the regions of the genome with higher genetic diversity and more rapid evolution. Accumulations of distinct genomic signatures and amino acid substitutions were observed in both parental and hybrid clones after relatively few generations. We have previously observed that there is a higher frequency of recombination in repeat-rich regions of the *T. cruzi* genome^7^, and it is likely that this is also the case for the hybrids. We can speculate that this is a likely explanation for variability in repeat and gene copy numbers between hybrid and non-hybrid clades in response to the distinct environmental conditions. Since these parasites were cultivated solely in culture, it is still not clear if this mechanism is related to the lack of pressure caused by the progression within triatomine bugs or by the interaction with the host immune system, or if these genomes display an intrinsic mechanism of high diversification of these regions. Recent comparative genomics of natural *T. cruzi* strains have shown a similar pattern, suggesting that these regions indeed evolve more rapidly than other regions of the genome^9^.

In summary, we confirmed that *in vitro* hybrid formation in *T. cruzi* happened in a parasexual mechanism, representing an important mechanism of genetic diversification. Tetraploid hybrids showed patterns of progressive genome erosion shaped by the environment’s selective pressure, which may explain distinct phenotypes in isolated natural hybrid populations. In addition, an increase in coding sequence diversity is observed in hybrids in comparison to the parental strains, related to both new mutations and most likely recombination events. Finally, our microevolution experiment has shown that repetitive gene families related to immune evasion evolved more rapidly than other regions of the genome, indicating an intrinsic mechanism of genetic variation of these regions.

## Methods

### Parasites and microevolution experiment

For routine culture, *T. cruzi* epimastigotes were grown axenically in supplemented RPMI as previously described^45^. Parasites were cloned by limiting dilution. The microevolution experiment comprised continuous *in vitro* cultures of each parasite line for five years. Three experimentally-generated hybrid clones (1C2, 1D12 and 2C1) and their two parents (P1 and P2) were seeded in the primary culture at 2 x 10^5^/ml and then allowed to grow for seven generations to stationary phase. Parasites were passaged into fresh media every two weeks at a seeding density of 2 x 10^5^/ml, equating typically to 1% of the stationary phase cultures.

### Flow cytometry

DNA content was determined as previously described^23^. In brief, mid-log phase epimastigotes were washed in PBS, then fixed overnight in ice-cold 70% methanol / 30% PBS. Fixed cells were washed in PBS, adjusted to 1 x 10^6^ cells/ml and incubated for 45 minutes with 10 μg/ml propidium iodide (PI) and 10 μg/ml RNAse A at 37°C. Fluorescence was detected using a FACSCalibur flow cytometer for a minimum of 10,000 events. FlowJo software (Tree Star Inc., Oregon, USA) was used to plot histograms and identify G1-0 and G2-M peaks. Mean G1-0 values were taken to infer relative DNA content. An internal control *T. cruzi* strain, Esm cl3, was included in each run. Relative DNA content values were calculated as ratios compared to the internal standard. For experimental hybrids, the ratios relative to each parent (P1 or P2) were also determined using the mean standard:parent ratios derived from 12 independent experiments.

### Microsatellite analysis

Genotyping was done for four microsatellite loci: MCLF10, 10101(TA), 7093(TC) and 10101(TC) as previously described^23^. Briefly, microsatellites were PCR amplified from genome DNA samples using fluorescently-labelled primers targeting conserved flanking regions. Amplicon lengths were determined using a 48-capillary 3730 DNA analyzer (Applied Biosystems, UK) and analysed using Genemapper v3.5 software (Applied Biosystems, UK). The size of different PCR products (alleles), visualised as fluorescence peaks, were determined automatically by the software using a size standard to calibrate the calculations. All allele size calls made by the software were checked manually against a library of known TcI alleles^46^.

### Whole genome sequencing

Total genomic DNA was isolated directly from long-term parasite cultures using the Gentra Puregene Tissue Kit (Qiagen) according to the manufacturers’ instructions. A total of 40 ng of genomic DNA was used as a template to prepare the sequencing libraries with the Rubicon Kit and the Illumina TruPlex® kit. For the two parent strains, a 180bp and 350 bp insert sizes paired end libraries were produced, plus an additional 8 Kb insert size mate paired library. For the hybrid offspring, a single paired end library was produced with an insert size of 350 bp. All libraries were sequenced using the Illumina HiSeq 2500 platform. The library complexity of each parent was analysed using the 17mer distribution of the Illumina libraries using Jellyfish^46^.

### *de novo* genome assembly

The genomes of the parent strains were assembled using the 180-bp Illumina paired end library and the 8 Kb Illumina mate pair library using the ALLPATHS-LG v52488 (https://software.broadinstitute.org/allpaths-lg/blog/) assembler with a K=96, using the TcI Sylvio X10/1 reference for evaluations. The resulting contigs were scaffolded using both Illumina libraries and the BESST scaffolder^48^. Later, gaps in the scaffolded assemblies were filled using the 180-bp with GapFiller^49^. Each genome was submitted for annotation, prior removal of repetitive elements using RepeatMasker (https://www.repeatmasker.org/), to the Companion annotation pipeline^50^.

### Data processing and read mapping

Paired end libraries were quality filtered using Nesoni Clip program (https://github.com/Victorian-Bioinformatics-Consortium/nesoni) removing bases with a Phred quality score below 30 and reads shorter than 64 nucleotides; later, sequencing adaptors were removed. Mate paired libraries were processed as above, plus an additional step to reverse complement the filtered reads. Filtered reads, from all the sequencing libraries produced, were mapped against the TcI Sylvio X10/1 reference genome (https://tritrypdb.org/tritrypdb/app/record/dataset/TMPTX_tcruSylvioX10-1) using Burrows-Wheeler Aligner^51^. The mapping files were sorted, PCR and optical duplicates were removed and read groups were added using Picard Tools v1.134 (https://broadinstitute.github.io/picard/). These mapping files were used for downstream analyses.

### Identification of genomic variation

Short variants, such as InDels and SNPs, were called using a mapping-based approach. Mapping files were used as input for the GATK variant caller (https://gatk.broadinstitute.org/hc/en-us) using a minimum quality value (QUAL) of 10 and a minimum depth of coverage (DP) of 10. To avoid the influence of low mapping quality in repeated regions, a strict mapping quality filter was applied and any SNP in regions with mapping quality below 50 was removed using BCFtools^52^.

VCFtools package^52^ was used to infer SNPdensity per 1kB (option *--SNPdensity*) and SNPs exclusive to each strain were identified using both VCFtools and BEDtools^53, 54^. The functional effect of these variants and the ratio of synonymous per non-synonymous mutations was predicted using SNPEff v4.4^55^.

### Chromosome copy number variation and ploidy estimation

Chromosome copy number variation (CCNV) and ploidy estimation were performed using a combination of Read Depth Coverage (RDC) and Allele Balance (AB) methods as previously described^30, 31^. Briefly, the estimation of CCNV was based on the ratio between the mean coverage of predicted single-copy genes in a given chromosome and the mean coverage of all single-copy genes in the genome. This approach was based on the RDC of 2602 1:1 orthologs between the haplotypes of the reference SylvioX10/1 genome and the parental strains assemblies, identified using OrthoVenn2 (https://orthovenn2.bioinfotoolkits.net/home). For chromosomes lacking single-copy genes, CCNV was estimated based on the ratio between the mean RDC of each chromosome position and the mean coverage of all genome positions. If the ratio between the median chromosome coverage and the median genome coverage was approximate one, the chromosome had the same copy as the genome overall, while fluctuations in this ratio (lower than 0.75 or higher than 1.25) were putative aneuploidies.

Heterozygous SNPs with a mapping quality higher than 50 were selected for AB analyses to confirm aneuploidies and to estimate chromosome somy and whole genome ploidy. Whole genome ploidy was estimated by analysing the proportion of the alleles in heterozygous SNP positions of all single-copy genes. To confirm the RDC somy estimations, the proportion of each allele with heterozygous SNPs per chromosome position was plotted for the 47 TcI chromosomes. All RDC and AB graphs were generated in R with ggplot2 (https://ggplot2.tidyverse.org), pheatmap (https://CRAN.R-project.org/package=pheatmap) and vcfR^56^ packages.

Genome erosion patterns were identified by copy number variation (CNV) estimations using Control-FREEC package^57^. Gene Ontology Enrichment analyses in CNV regions were performed using TriTryp tools (https://tritrypdb.org/tritrypdb/app/) and surface molecules genes were assigned from the available TcI Sylvio X10/1 annotation^7^.

## Data Access

The raw data generated in this study have been submitted to the NCBI BioProject database (https://www.ncbi.nlm.nih.gov/bioproject/) under accession number PRJNA748998.

## Acknowledgments

This study was financed in part by the Swedish Research Council, Dnr 20016-02951 and Coordenação de Aperfeiçoamento de Pessoal de Nível Superior – CAPES (Brazilian Government Agency) - Finance Code 001, GMM was supported by a scholarship provided by CAPES-PrInt and ECG was funded by grants from CNPq (Brazilian Government Agency). We are thankful to João Luís Reis Cunha for his guidance on ploidy estimation.

## Author contributions

BA, MAM, CTL and GMM designed the study and the analyses, and wrote the manuscript. ML carried out the in vitro work, while CTL designed the genome sequencing strategy and carried out initial assemblies and analyses. GMM carried out the analysis of the hybrid data. MY and LAM contributed to the parasite work. ECG contributed to the mutation analysis and the writing.

