## Supplementary figures for "Experimental microevolution of *Trypanosoma cruzi* reveals hybridization and clonal mechanisms driving rapid diversification of genome sequence and structure"

[2] - Departamento de Biologia Celular, Embriologia e Genética, Universidade Federal de Santa Catarina, Florianópolis, Brazil

[3] - Faculty of Infectious and Tropical Diseases, London School of Hygiene and Tropical Medicine, London, United Kingdom.

[4] - Departamento de Microbiologia, Imunologia e Parasitologia, Universidade Federal de Santa Catarina, Florianópolis, Brazil

[5] - These authors contributed equally to this work

**Running title:** Following genetic exchange in *Trypansoma cruzi* in real time using whole genome sequencing reveals tetraploid and triploid stages

**Keywords:** *Trypanosoma cruzi*, genetic exchange, hybrid formation, parasitism, Chagas disease

**Supplementary Figures**

**Supplementary Fig. 1:** Somy estimation per chromosome based on AB in starting generation confirming aneuploidies in essentially diploid parental strain 1. **(Page: 4)**

**Supplementary Fig. 2:** Somy estimation per chromosome based on AB in starting generation confirming aneuploidies in essentially diploid parental strain 2. **(Page: 6)**

**Supplementary Fig. 3:** Somy estimation per chromosome based on AB in starting generation confirming tetrasomic chromosomes in hybrid 1C2. **(Page: 8)**

**Supplementary Fig. 4:** Somy estimation per chromosome based on AB in starting generation confirming tetrasomic chromosomes in hybrid 1D12. **(Page: 10)**

**Supplementary Fig. 5:** Somy estimation per chromosome based on AB in starting generation confirming tetrasomic chromosomes in hybrid 2C1. **(Page: 12)**

**Supplementary Fig. 6:** Somy estimation per chromosome based on AB confirming novel aneuploidies after *in vitro* culture in the three essentially diploid clone replicates of parental strain 1. **(Page: 14)**

**Supplementary Fig. 7:** Somy estimation per chromosome based on AB confirming novel aneuploidies after *in vitro* culture in the three essentially diploid clone replicates of parental strain 2. **(Page: 20)**

**Supplementary Fig. 8:** Somy estimation per chromosome based on AB confirming trisomic and tetrasomic chromosomes after *in vitro* culture in the three clone replicates of hybrid 1C2. **(Page: 26)**

**Supplementary Fig. 9:** Somy estimation per chromosome based on AB confirming trisomic and tetrasomic chromosomes after *in vitro* culture in the three clone replicates of hybrid 1D12. **(Page: 32)**

**Supplementary Fig. 10:** Somy estimation per chromosome based on AB confirming trisomic and tetrasomic chromosomes after *in vitro* culture in the three clone replicates of hybrid 2C1. **(Page: 38)**

**Supplementary Fig. 11:** Counts of genes displaying copy number variations after culture growth. **(Page: 44)**

**Supplementary Tables**

**Supplementary Table 1:** PCR-based multilocus microsatellite genotyping in parental and hybrid strains before (t=0) and after (t=800) the microevolution experiment. **(Page: 45)**

**Supplementary Table 2**: Genome assembly statistics of parental strains. **(Page: 46)**

**Supplementary Table 3**: Effects of SNPs in the first generation hybrids. **(Page: 47)**

**Supplementary Table 4:** Number of SNPs in surface molecules genes and in other genes. **(Page: 48)**

**Supplementary Table 5:** Number of non-synonymous SNPs in surface molecules genes and in other genes. **(Page: 49)**

**Supplementary Table 6:** Gene Ontology analysis of expanded genes in parental strains after culture growth. **(Page: 50)**

**Supplementary Table 7**: Gene Ontology analysis of contracted genes in parental strains after culture growth. **(Page: 53)**

**Supplementary Table 8:** Gene Ontology analysis of expanded genes in hybrid strains after culture growth. **(Page: 55)**

**Supplementary Table 9**: Gene Ontology analysis of contracted genes in hybrid strains after culture growth. **(Page: 59)**

**Supplementary Table 10**: Gene Ontology analysis of genes encoded in chromosome 19. **(Page: 60)**

**Supplementary Table 11**: Gene Ontology analysis of genes encoded in chromosome 37. **(Page: 63)**

**Supplementary Table 12**: Gene Ontology analysis of contracted genes in Parental 2 clones after culture growth. **(Page: 64)**

**Supplementary Table 13**: Number of non-synonymous SNPs in surface molecules genes and in other genes after culture growth. **(Page: 66)**

**Supplementary Fig. 1:** Somy estimation per chromosome based on AB in starting generation confirming aneuploidies in essentially diploid parental strain 1. Blue points represent the proportion of the alleles in heterozygous SNP positions along the chromosome. Darker blue represents the frequency of the first allele while lighter blue represents the frequency of the second allele, black lines represent the median value in windows.

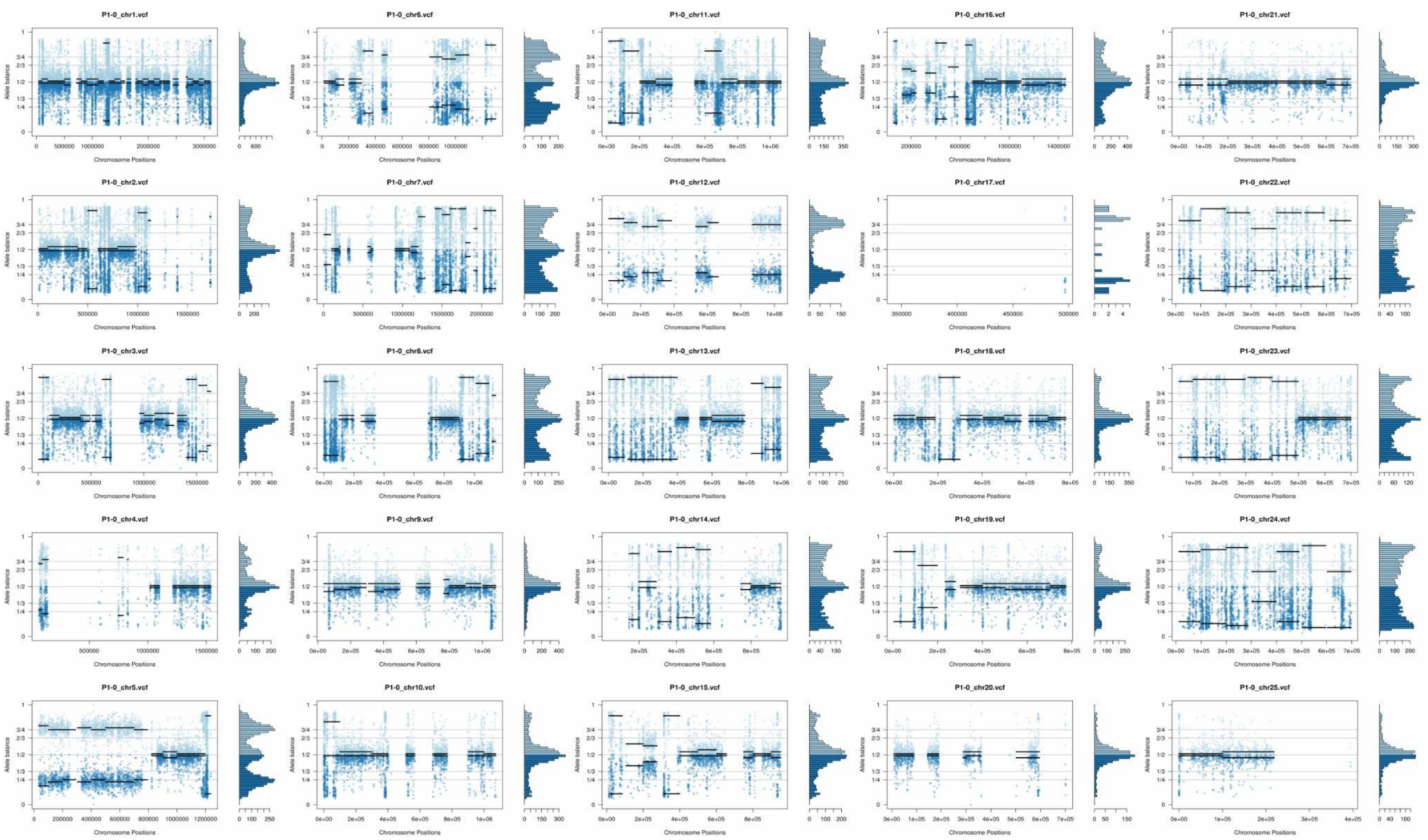

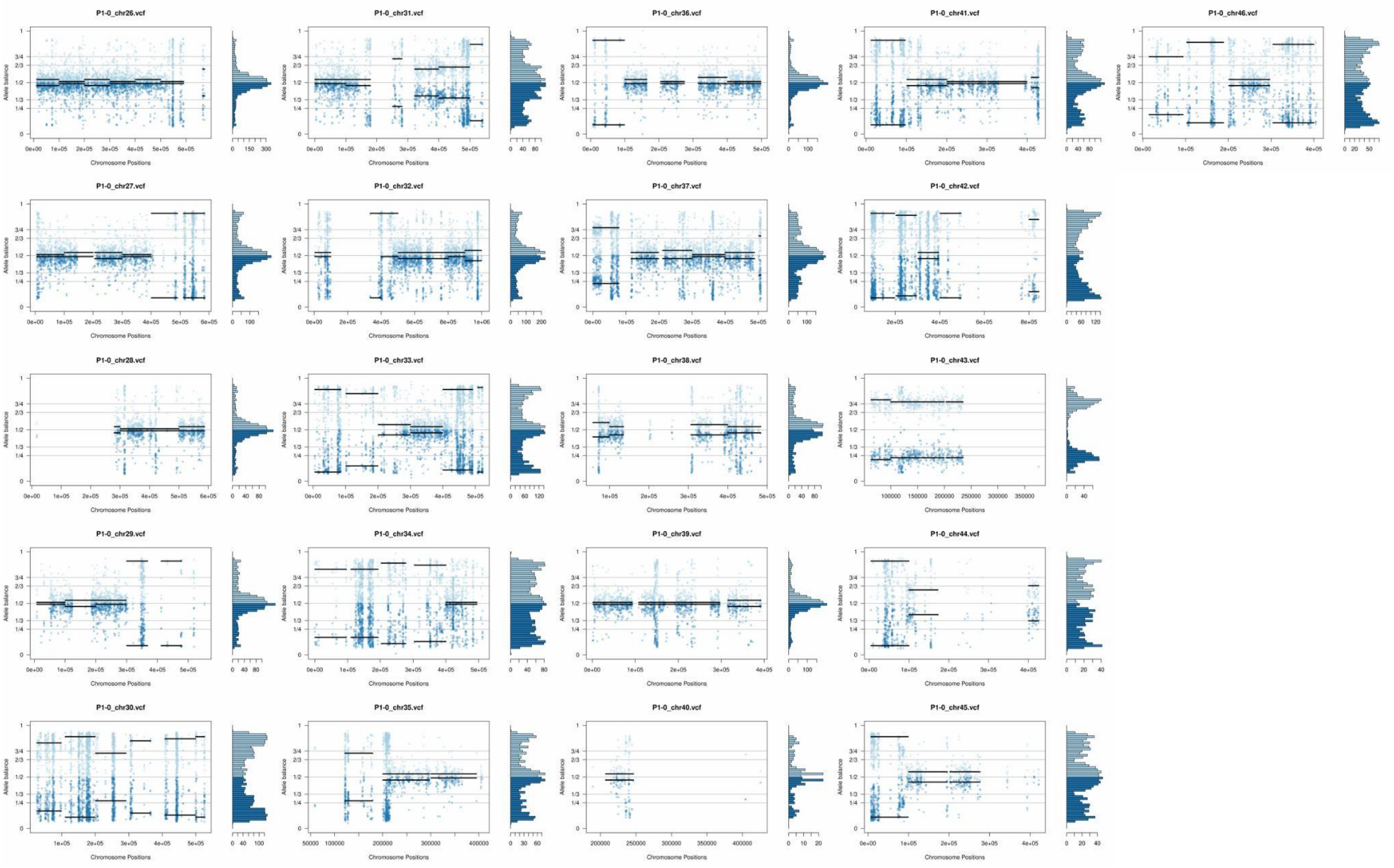
P1-0_chr26.vcf
P1-0_chr31.vcf
P1-0_chr36.vcf
P1-0_chr41.vcf
P1-0_chr46.vcf
P1-0_chr27.vcf
P1-0_chr32.vcf
P1-0_chr37.vcf
P1-0_chr42.vcf
P1-0_chr28.vcf
P1-0_chr33.vcf
P1-0_chr38.vcf
P1-0_chr43.vcf
P1-0_chr29.vcf
P1-0_chr34.vcf
P1-0_chr39.vcf
P1-0_chr44.vcf
P1-0_chr30.vcf
P1-0_chr35.vcf
P1-0_chr40.vcf
P1-0_chr45.vcf
Allele balance
Chromosome Positions

**Supplementary Fig. 2:** Somy estimation per chromosome based on AB in starting generation confirming aneuploidies in essentially diploid parental strain 2. Blue points represent the proportion of the alleles in heterozygous SNP positions along the chromosome. Darker blue represents the frequency of the first allele while lighter blue represents the frequency of the second allele, black lines represent the median value in windows.

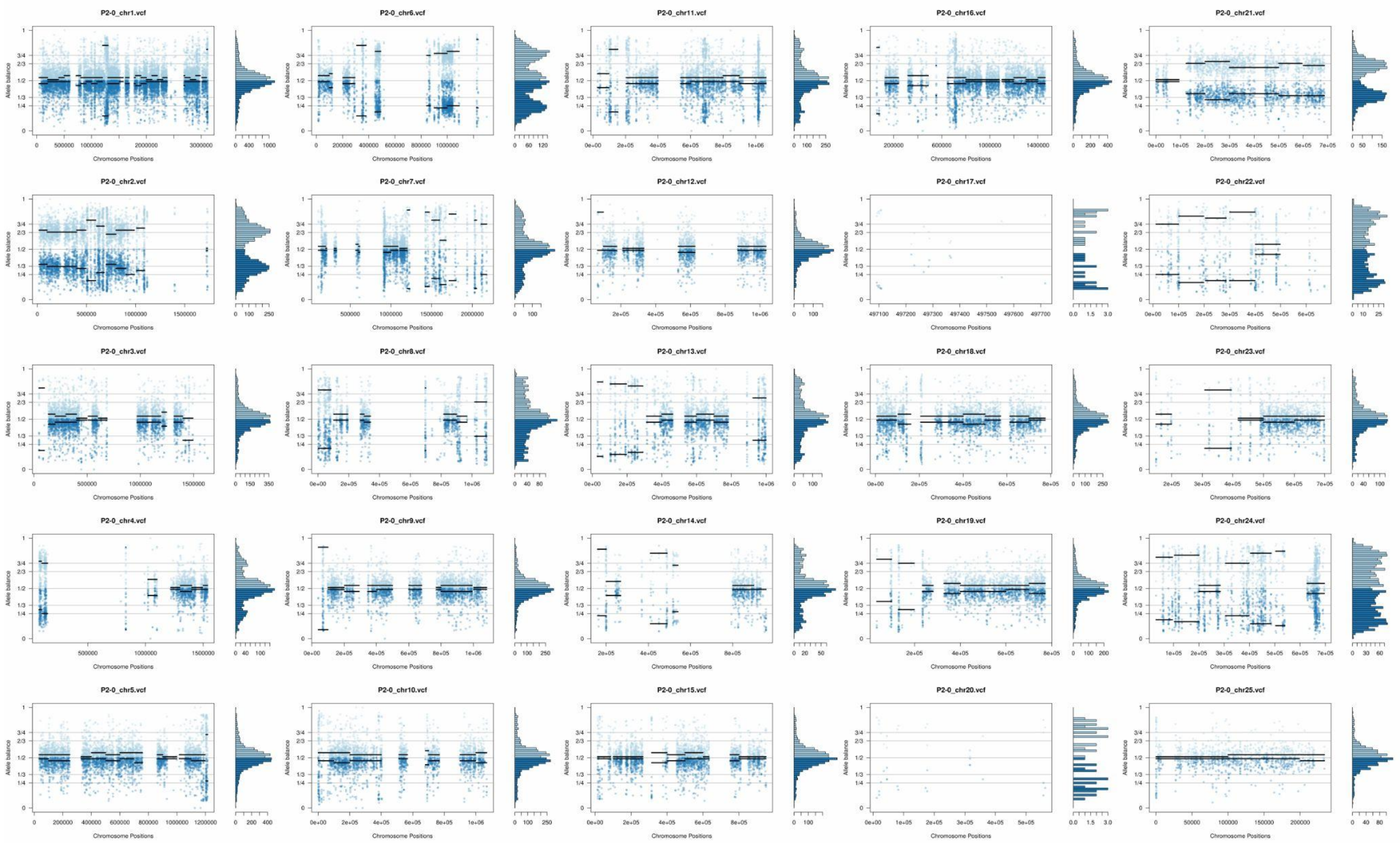

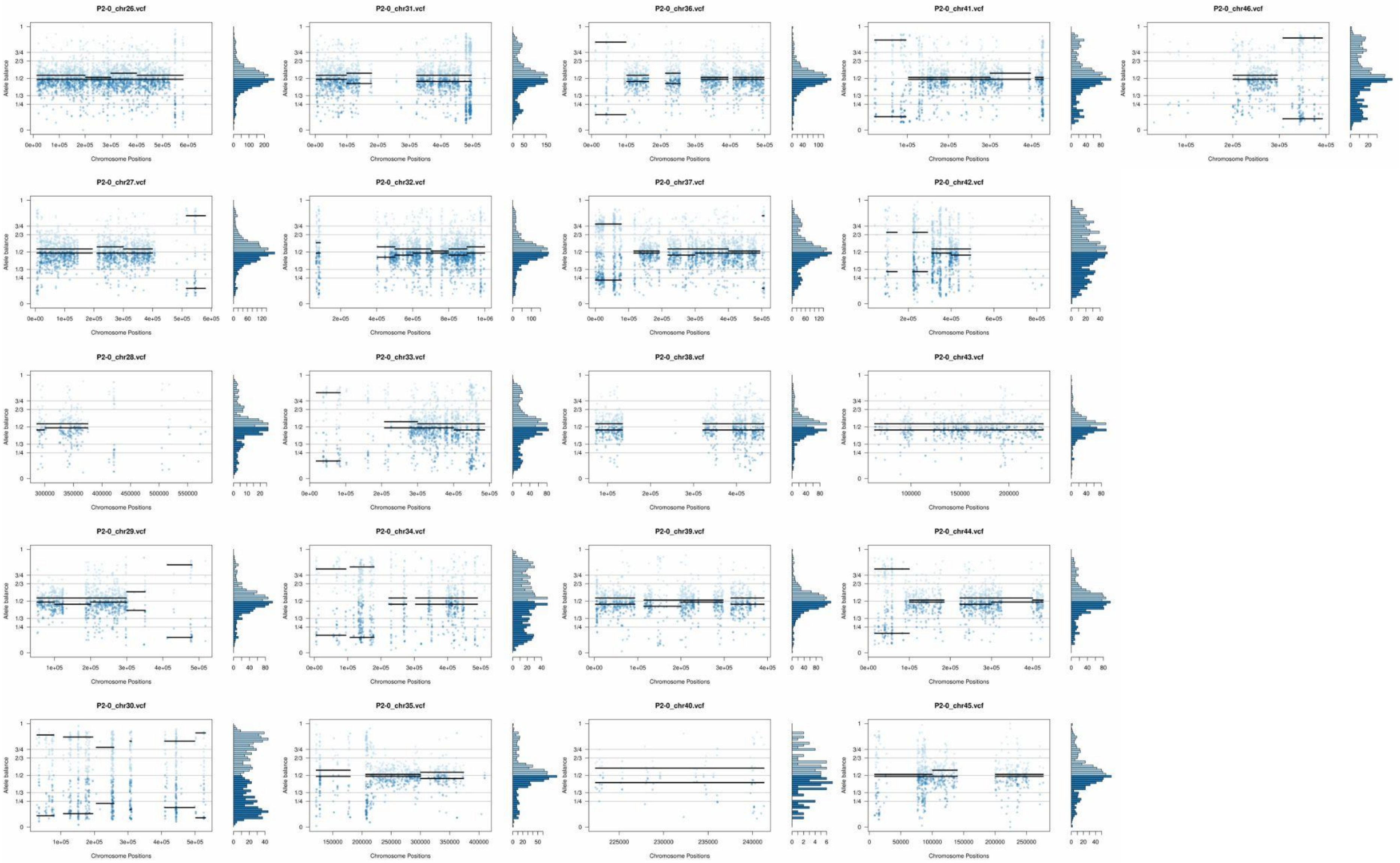
P2-0_chr26.vcf
P2-0_chr31.vcf
P2-0_chr36.vcf
P2-0_chr41.vcf
P2-0_chr46.vcf
P2-0_chr27.vcf
P2-0_chr32.vcf
P2-0_chr37.vcf
P2-0_chr42.vcf
P2-0_chr28.vcf
P2-0_chr33.vcf
P2-0_chr38.vcf
P2-0_chr43.vcf
P2-0_chr29.vcf
P2-0_chr34.vcf
P2-0_chr39.vcf
P2-0_chr44.vcf
P2-0_chr30.vcf
P2-0_chr35.vcf
P2-0_chr40.vcf
P2-0_chr45.vcf
Allele balance
Chromosome Positions

**Supplementary Fig. 3:** Somy estimation per chromosome based on AB in starting generation confirming tetrasomic chromosomes in hybrid 1C2. Blue points represent the proportion of the alleles in heterozygous SNP positions along the chromosome. Darker blue represents the frequency of the first allele while lighter blue represents the frequency of the second allele, black lines represent the median value in windows.

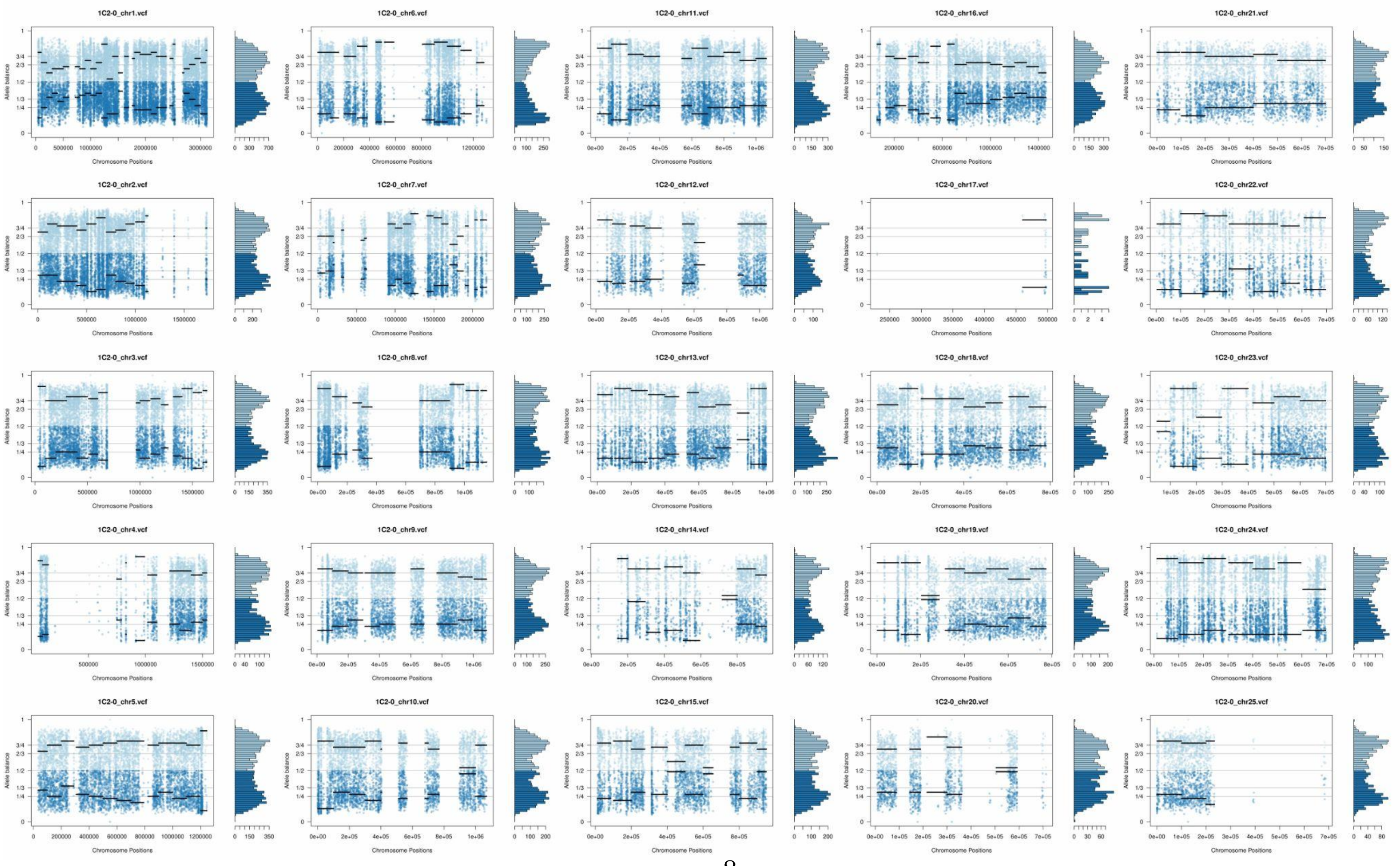

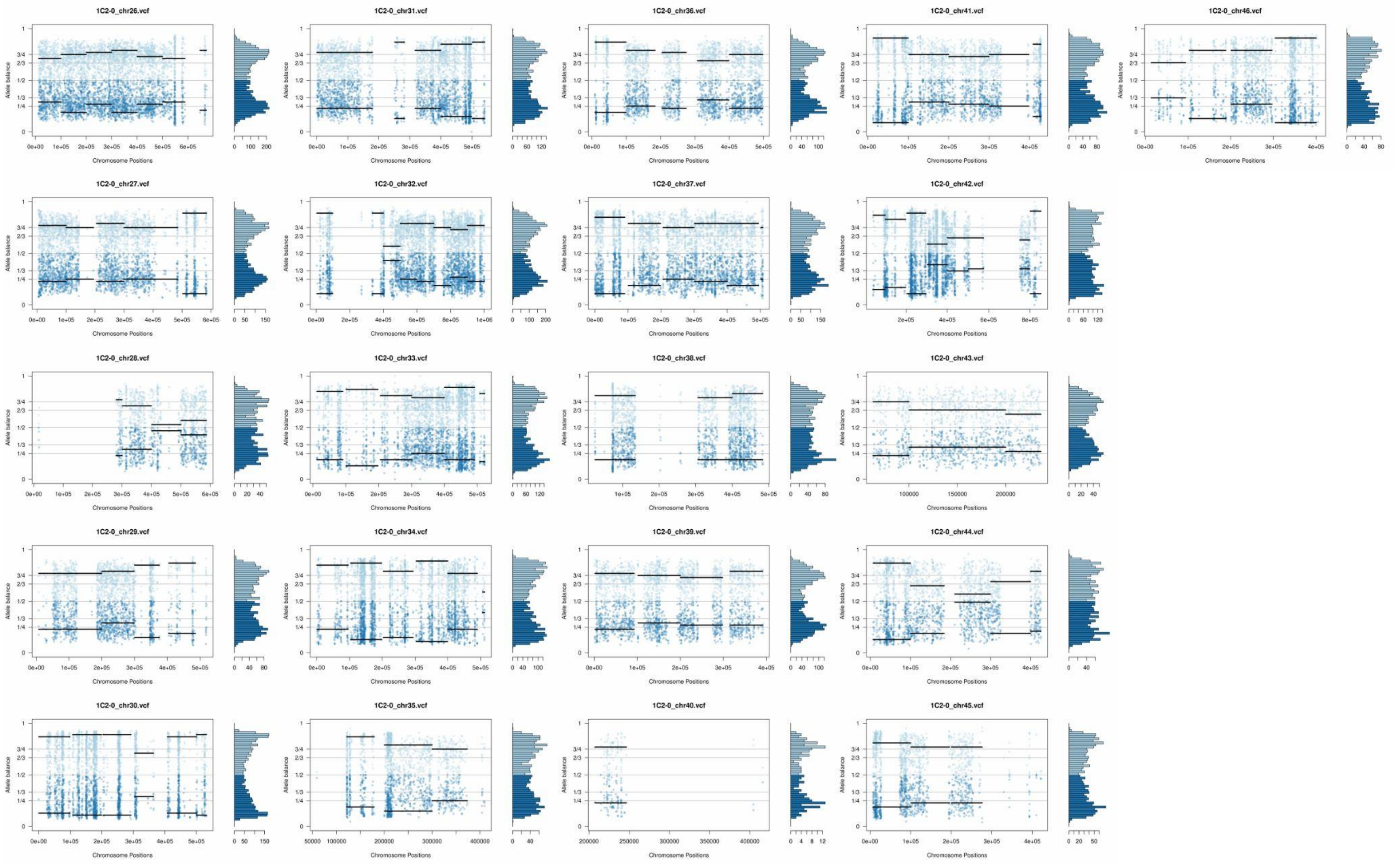
1C2-0_chr26.vcf
1C2-0_chr31.vcf
1C2-0_chr36.vcf
1C2-0_chr41.vcf
1C2-0_chr46.vcf
1C2-0_chr27.vcf
1C2-0_chr32.vcf
1C2-0_chr37.vcf
1C2-0_chr42.vcf
1C2-0_chr28.vcf
1C2-0_chr33.vcf
1C2-0_chr38.vcf
1C2-0_chr43.vcf
1C2-0_chr29.vcf
1C2-0_chr34.vcf
1C2-0_chr39.vcf
1C2-0_chr44.vcf
1C2-0_chr30.vcf
1C2-0_chr35.vcf
1C2-0_chr40.vcf
1C2-0_chr45.vcf
Allele balance
Chromosome Positions

**Supplementary Fig. 4:** Somy estimation per chromosome based on AB in starting generation confirming tetrasomic chromosomes in hybrid 1D12. Blue points represent the proportion of the alleles in heterozygous SNP positions along the chromosome. Darker blue represents the frequency of the first allele while lighter blue represents the frequency of the second allele, black lines represent the median value in windows.

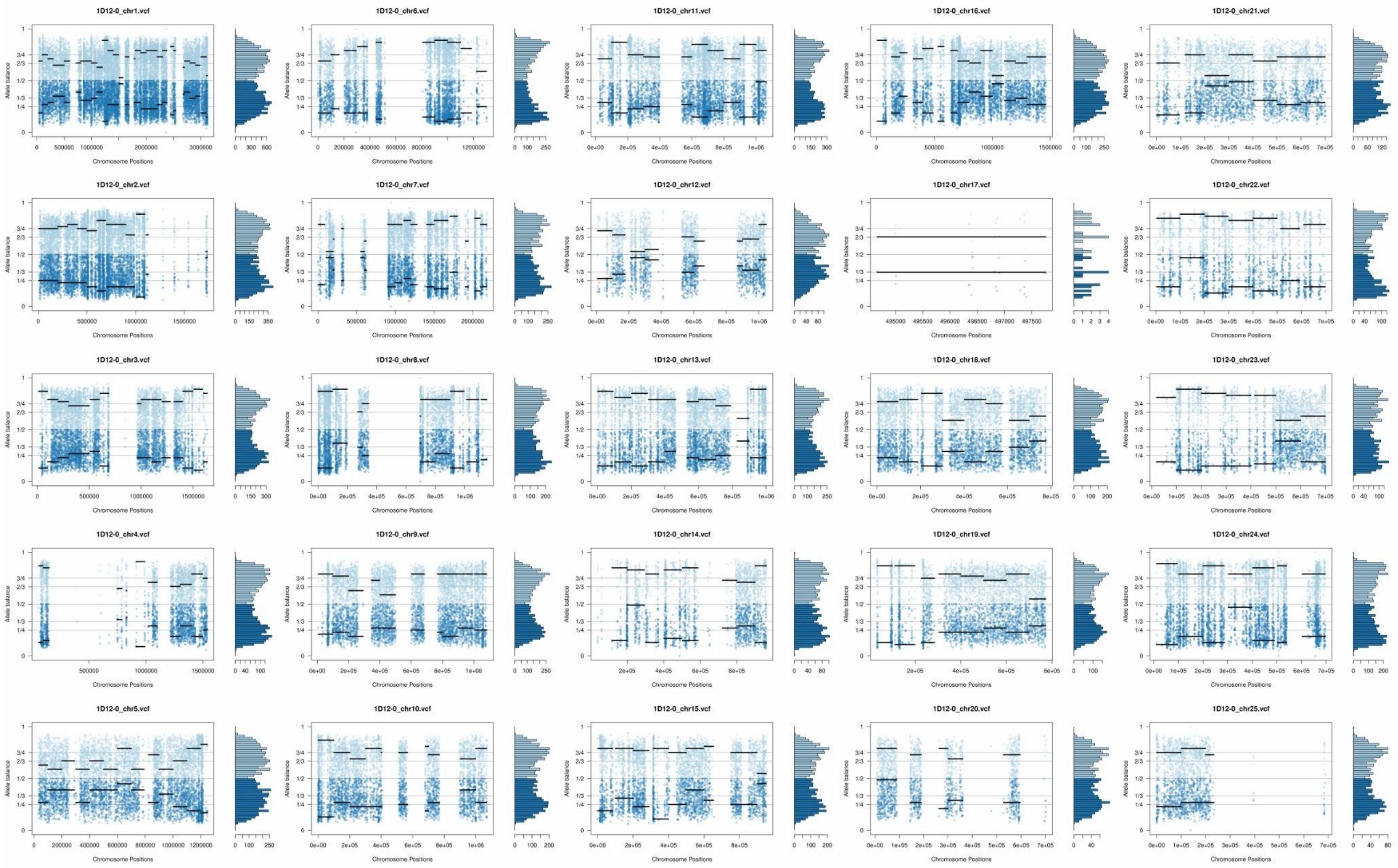

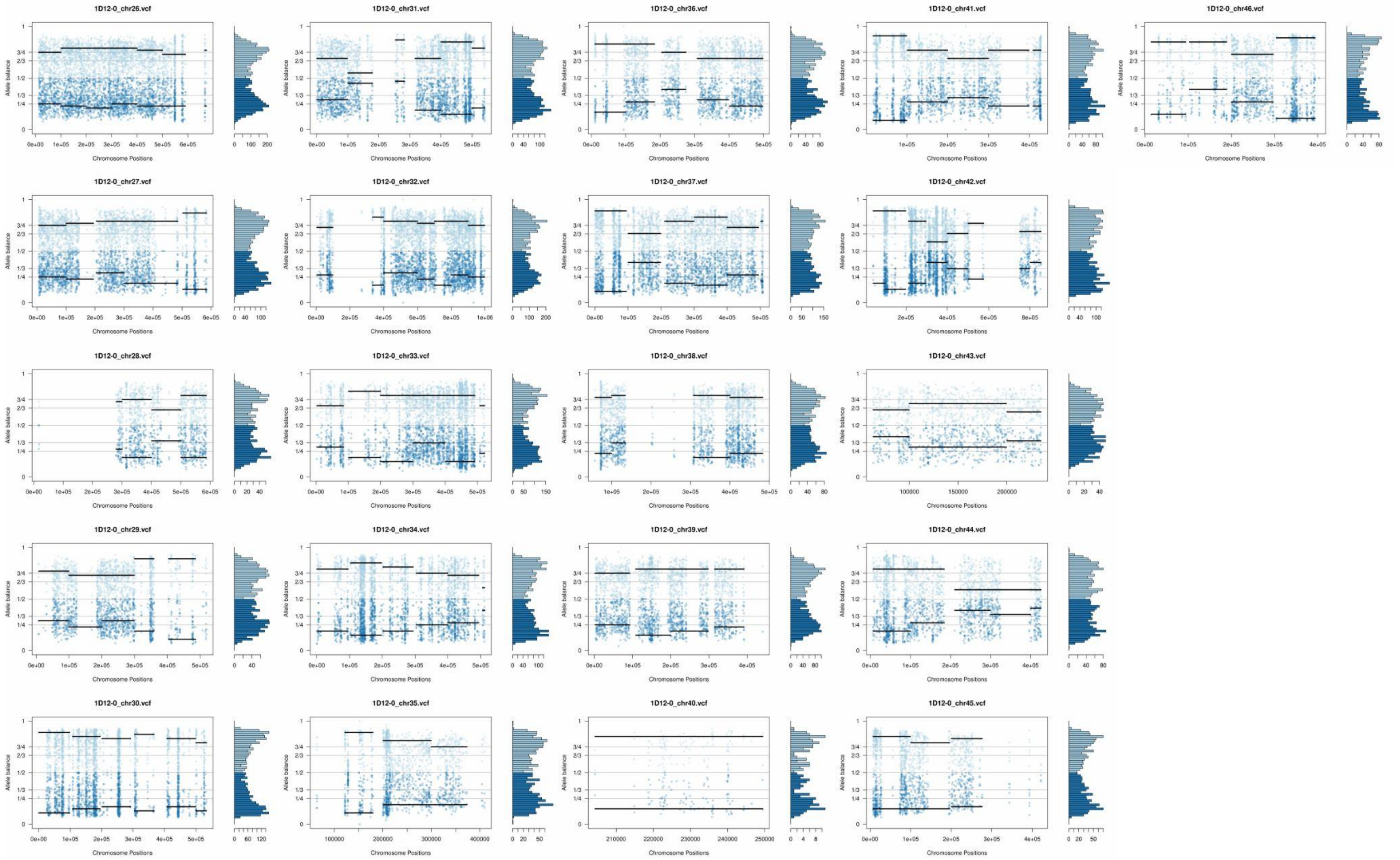
1D12-0_chr26.vcf
1D12-0_chr31.vcf
1D12-0_chr36.vcf
1D12-0_chr41.vcf
1D12-0_chr46.vcf
1D12-0_chr27.vcf
1D12-0_chr32.vcf
1D12-0_chr37.vcf
1D12-0_chr42.vcf
1D12-0_chr28.vcf
1D12-0_chr33.vcf
1D12-0_chr38.vcf
1D12-0_chr43.vcf
1D12-0_chr29.vcf
1D12-0_chr34.vcf
1D12-0_chr39.vcf
1D12-0_chr44.vcf
1D12-0_chr30.vcf
1D12-0_chr35.vcf
1D12-0_chr40.vcf
1D12-0_chr45.vcf
Allele balance
Chromosome Positions

**Supplementary Fig. 5:** Somy estimation per chromosome based on AB in starting generation confirming tetrasomic chromosomes in hybrid 2C1. Blue points represent the proportion of the alleles in heterozygous SNP positions along the chromosome. Darker blue represents the frequency of the first allele while lighter blue represents the frequency of the second allele, black lines represent the median value in windows.

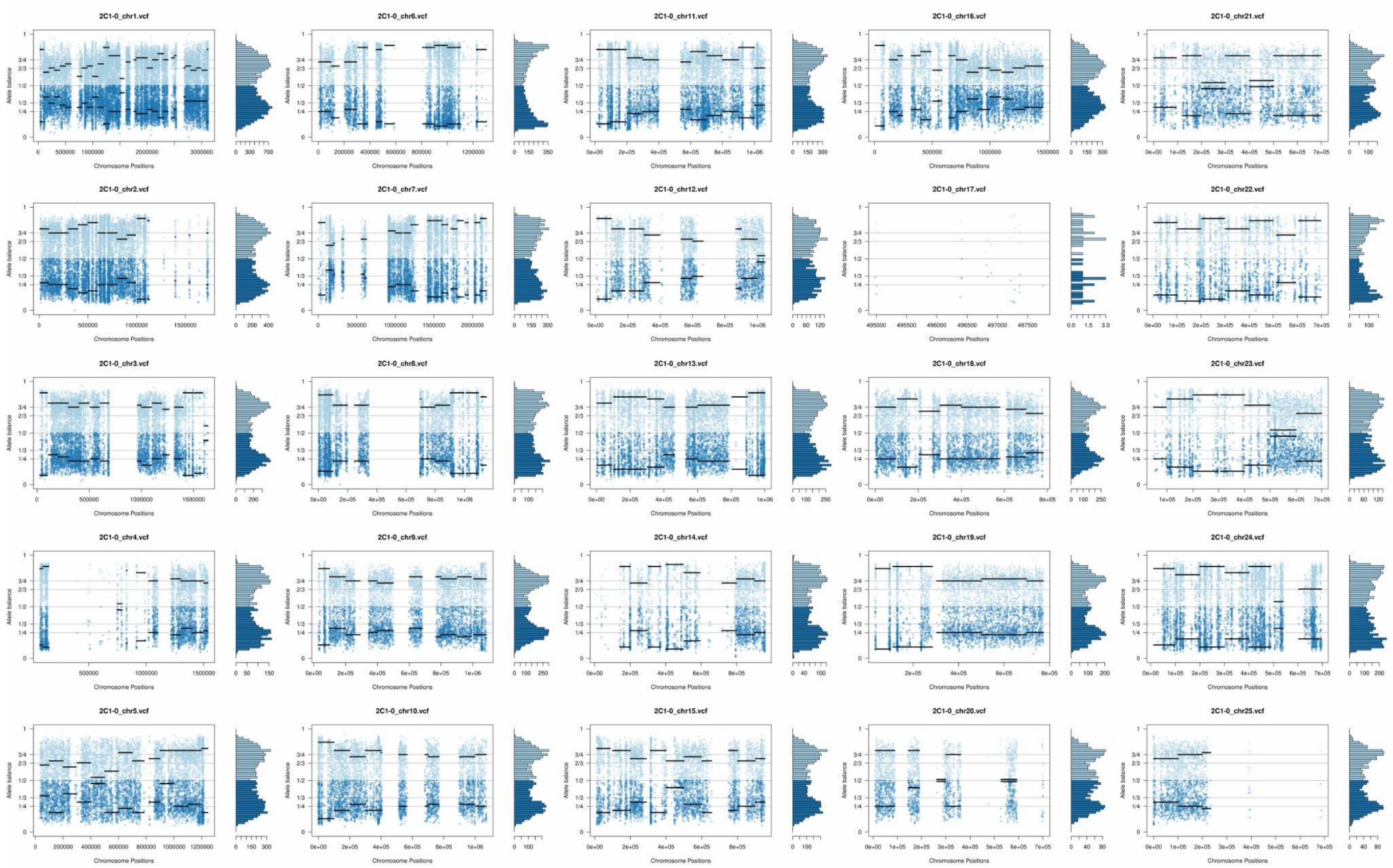

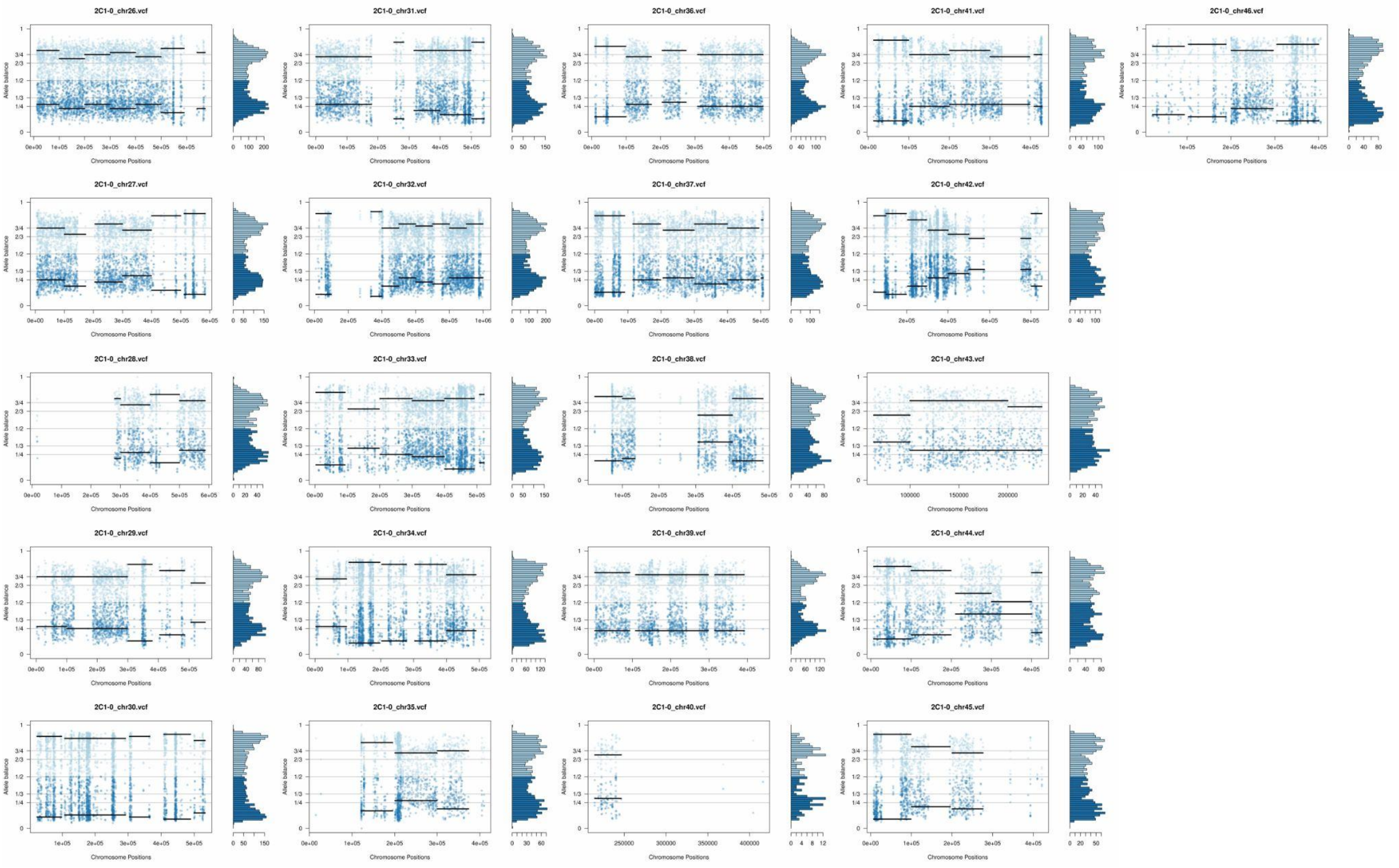
2C1-0_chr26.vcf
2C1-0_chr31.vcf
2C1-0_chr36.vcf
2C1-0_chr41.vcf
2C1-0_chr46.vcf
2C1-0_chr27.vcf
2C1-0_chr32.vcf
2C1-0_chr37.vcf
2C1-0_chr42.vcf
2C1-0_chr28.vcf
2C1-0_chr33.vcf
2C1-0_chr38.vcf
2C1-0_chr43.vcf
2C1-0_chr29.vcf
2C1-0_chr34.vcf
2C1-0_chr39.vcf
2C1-0_chr44.vcf
2C1-0_chr30.vcf
2C1-0_chr35.vcf
2C1-0_chr40.vcf
2C1-0_chr45.vcf
Allele balance
Chromosome Positions

**Supplementary Fig. 6:** Somy estimation per chromosome based on AB confirming novel aneuploidies after *in vitro* culture in the three essentially diploid clone replicates of parental strain 1. Orange points represent the proportion of the alleles in heterozygous SNP positions along the chromosome. Darker orange represents the frequency of the first allele while lighter orange represents the frequency of the second allele, red lines represent the median value in windows.

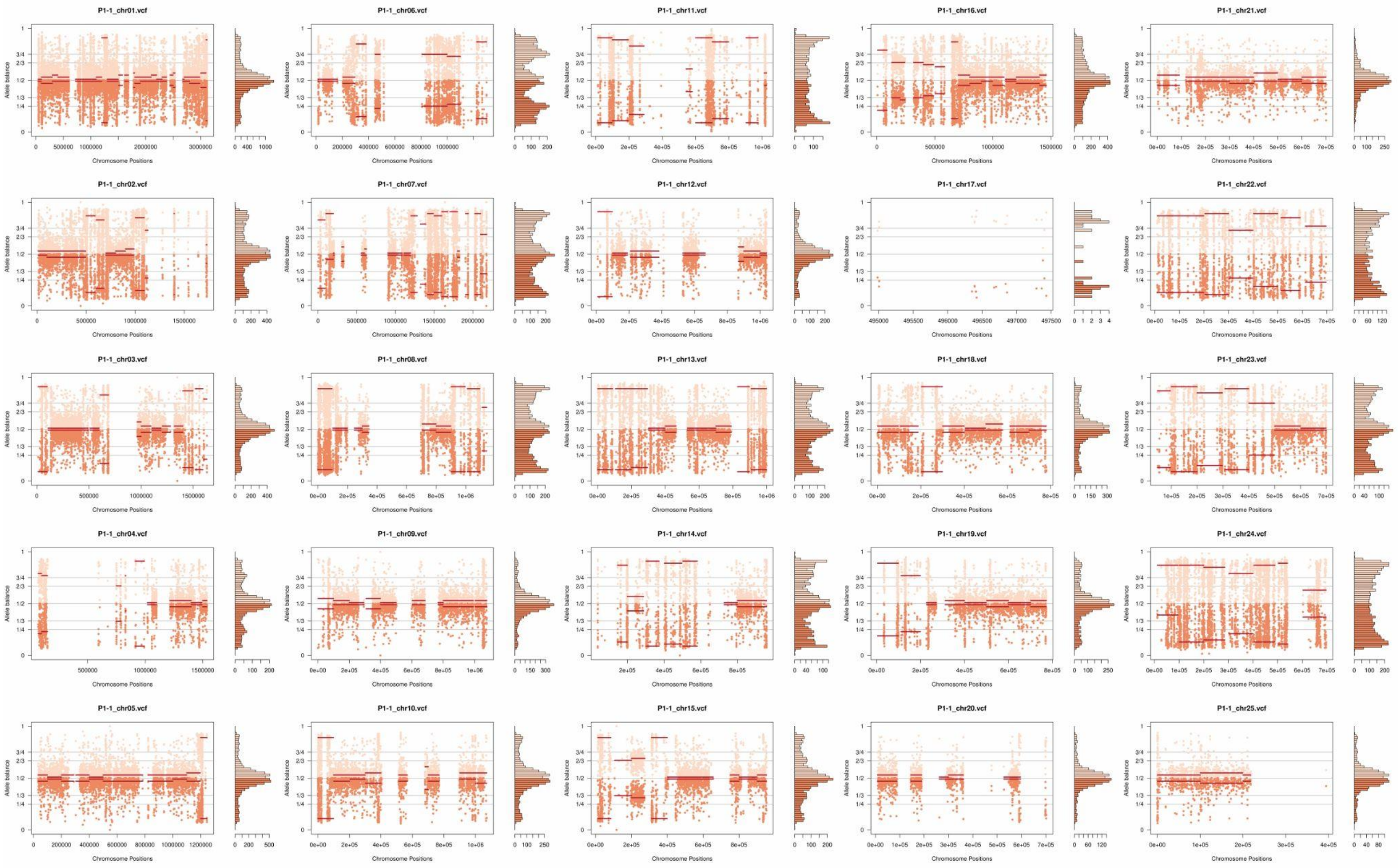

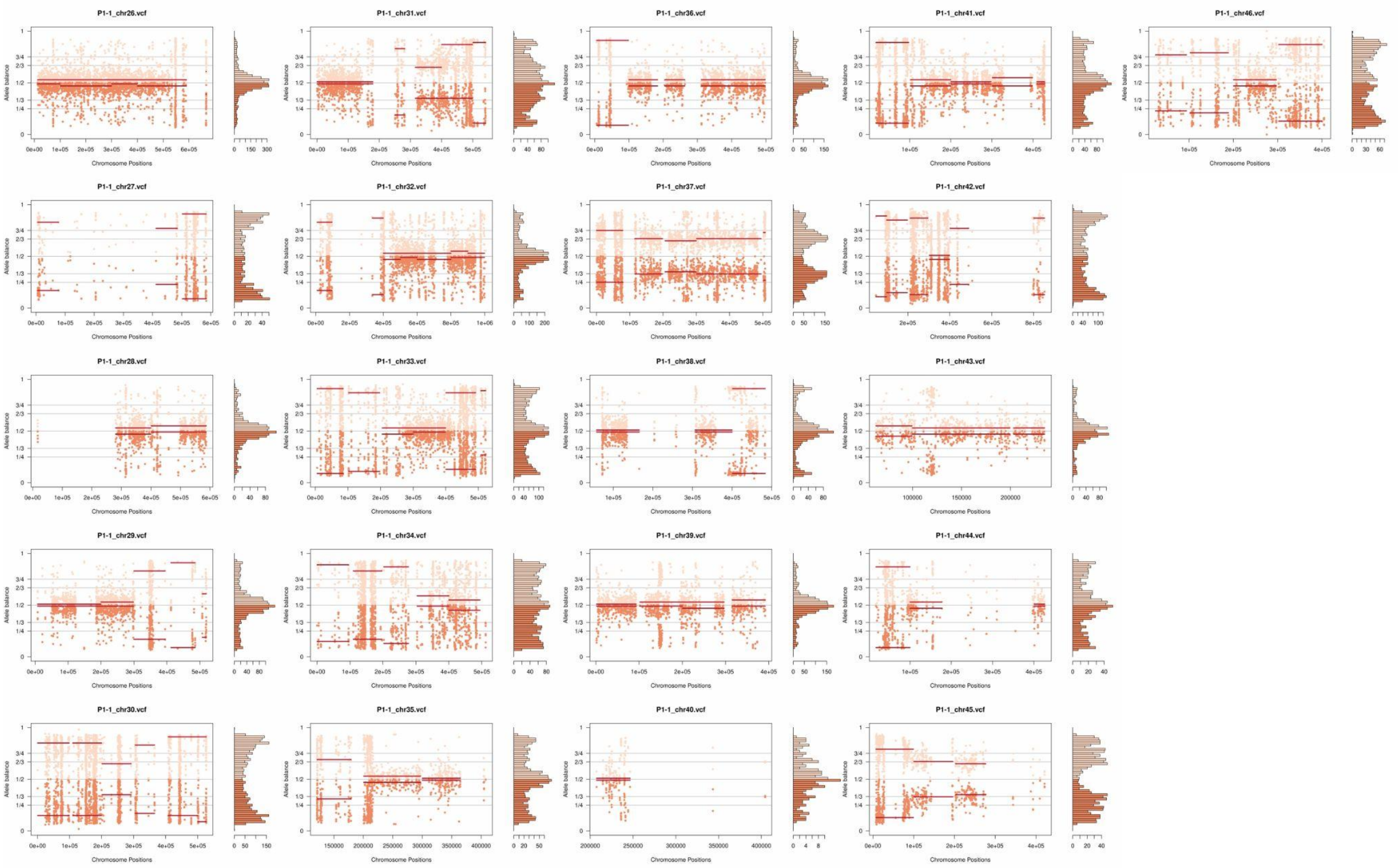
P1-1_chr26.vcf
P1-1_chr31.vcf
P1-1_chr36.vcf
P1-1_chr41.vcf
P1-1_chr46.vcf
P1-1_chr27.vcf
P1-1_chr32.vcf
P1-1_chr37.vcf
P1-1_chr42.vcf
P1-1_chr28.vcf
P1-1_chr33.vcf
P1-1_chr38.vcf
P1-1_chr43.vcf
P1-1_chr29.vcf
P1-1_chr34.vcf
P1-1_chr39.vcf
P1-1_chr44.vcf
P1-1_chr30.vcf
P1-1_chr35.vcf
P1-1_chr40.vcf
P1-1_chr45.vcf
Allele balance
Chromosome Positions

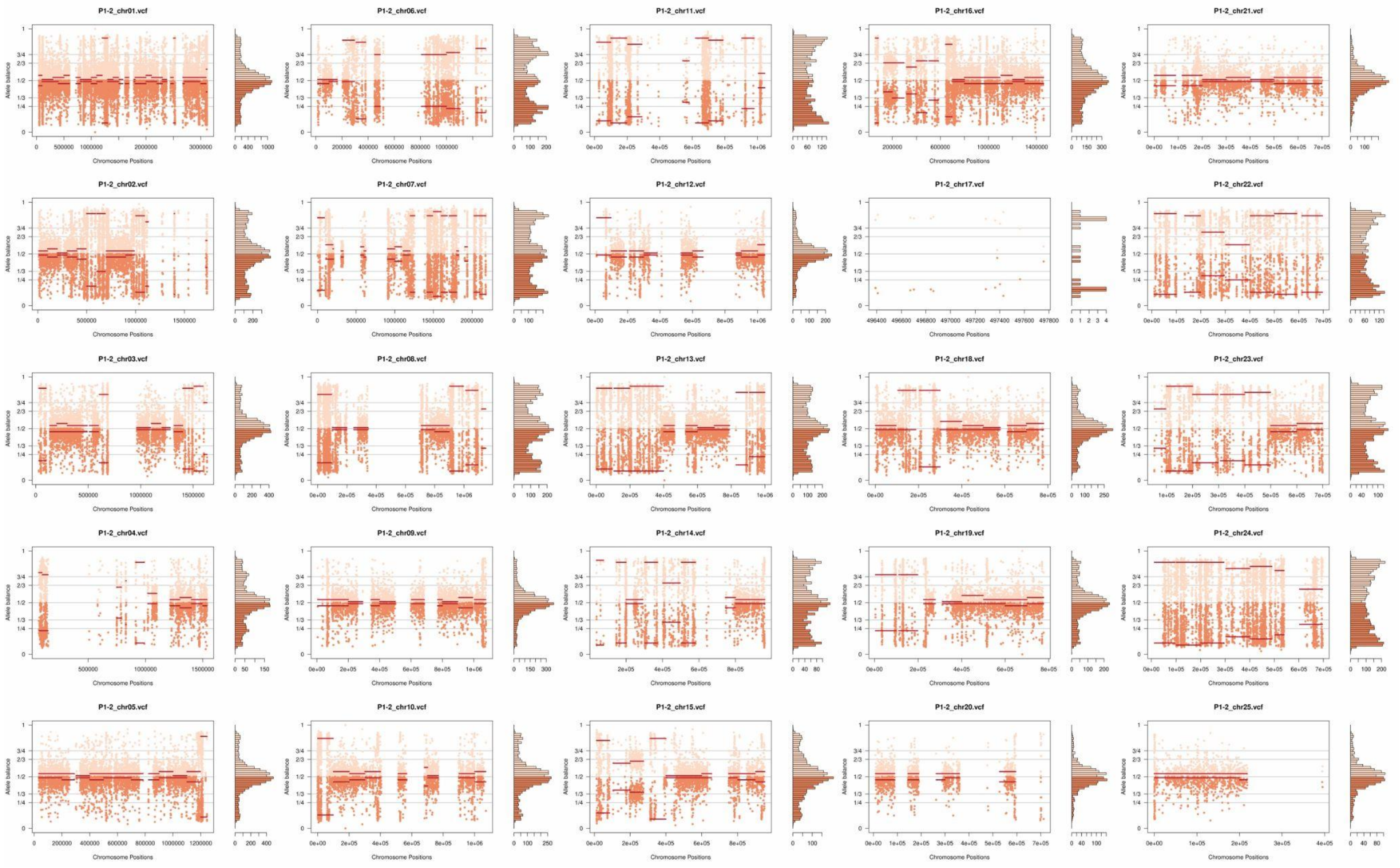
P1-2_chr01.vcf
P1-2_chr06.vcf
P1-2_chr11.vcf
P1-2_chr16.vcf
P1-2_chr21.vcf
P1-2_chr02.vcf
P1-2_chr07.vcf
P1-2_chr12.vcf
P1-2_chr17.vcf
P1-2_chr22.vcf
P1-2_chr03.vcf
P1-2_chr08.vcf
P1-2_chr13.vcf
P1-2_chr18.vcf
P1-2_chr23.vcf
P1-2_chr04.vcf
P1-2_chr09.vcf
P1-2_chr14.vcf
P1-2_chr19.vcf
P1-2_chr24.vcf
P1-2_chr05.vcf
P1-2_chr10.vcf
P1-2_chr15.vcf
P1-2_chr20.vcf
P1-2_chr25.vcf
Allele balance
Chromosome Positions

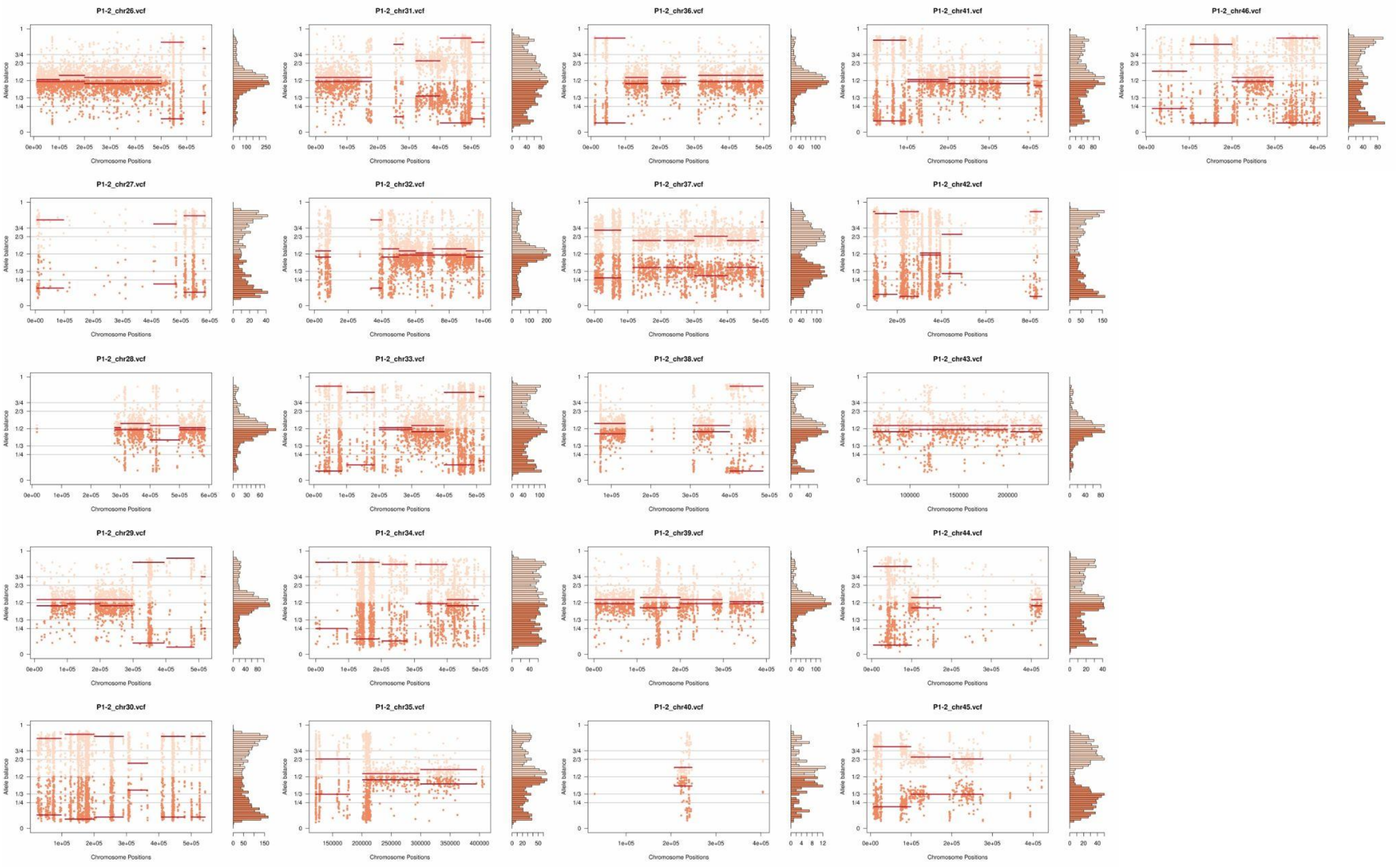
P1-2_chr26.vcf
P1-2_chr31.vcf
P1-2_chr36.vcf
P1-2_chr41.vcf
P1-2_chr46.vcf
P1-2_chr27.vcf
P1-2_chr32.vcf
P1-2_chr37.vcf
P1-2_chr42.vcf
P1-2_chr28.vcf
P1-2_chr33.vcf
P1-2_chr38.vcf
P1-2_chr43.vcf
P1-2_chr29.vcf
P1-2_chr34.vcf
P1-2_chr39.vcf
P1-2_chr44.vcf
P1-2_chr30.vcf
P1-2_chr35.vcf
P1-2_chr40.vcf
P1-2_chr45.vcf
Allele balance
Chromosome Positions

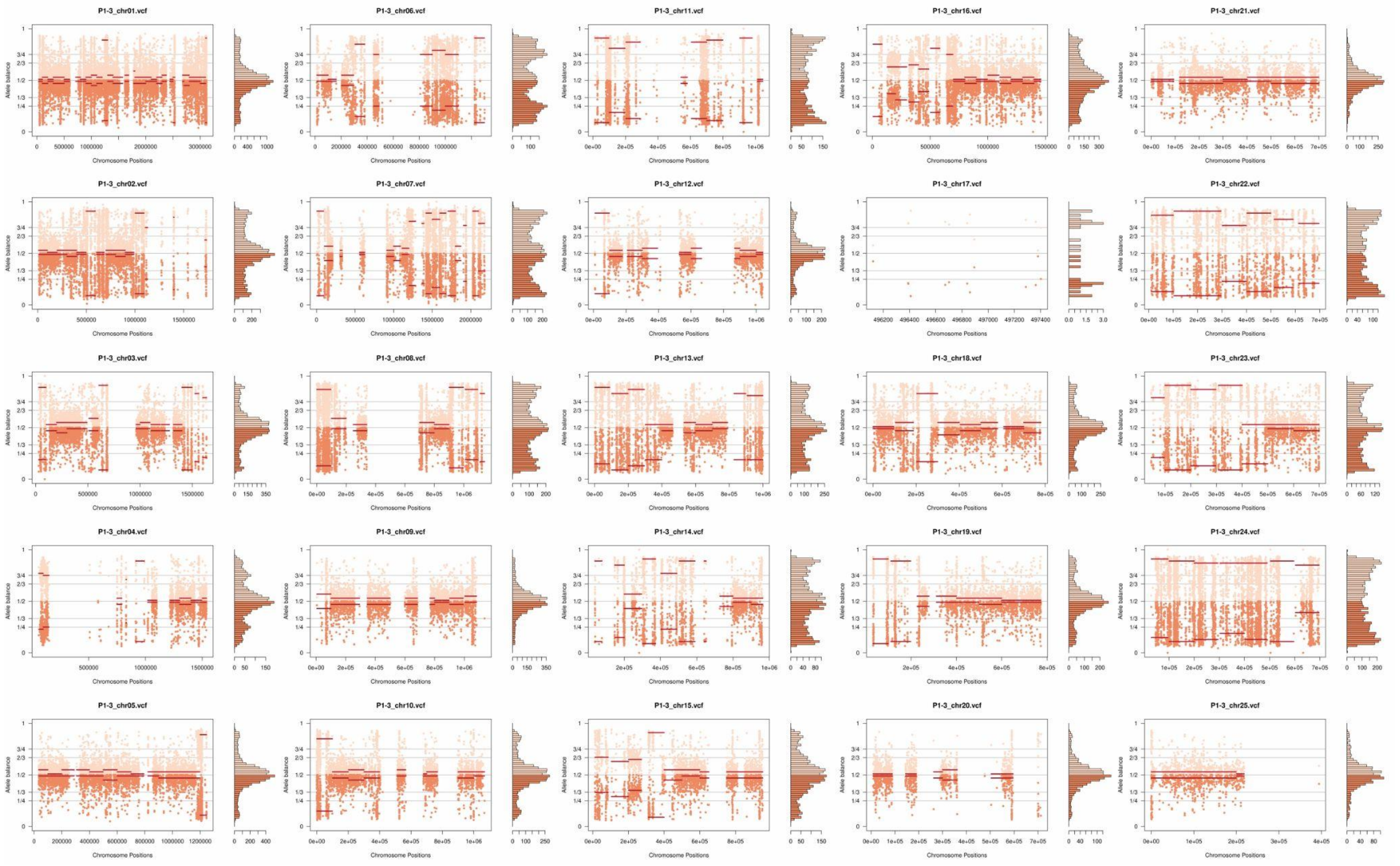
P1-3_chr01.vcf
P1-3_chr06.vcf
P1-3_chr11.vcf
P1-3_chr16.vcf
P1-3_chr21.vcf
P1-3_chr02.vcf
P1-3_chr07.vcf
P1-3_chr12.vcf
P1-3_chr17.vcf
P1-3_chr22.vcf
P1-3_chr03.vcf
P1-3_chr08.vcf
P1-3_chr13.vcf
P1-3_chr18.vcf
P1-3_chr23.vcf
P1-3_chr04.vcf
P1-3_chr09.vcf
P1-3_chr14.vcf
P1-3_chr19.vcf
P1-3_chr24.vcf
P1-3_chr05.vcf
P1-3_chr10.vcf
P1-3_chr15.vcf
P1-3_chr20.vcf
P1-3_chr25.vcf
Allele balance
Chromosome Positions

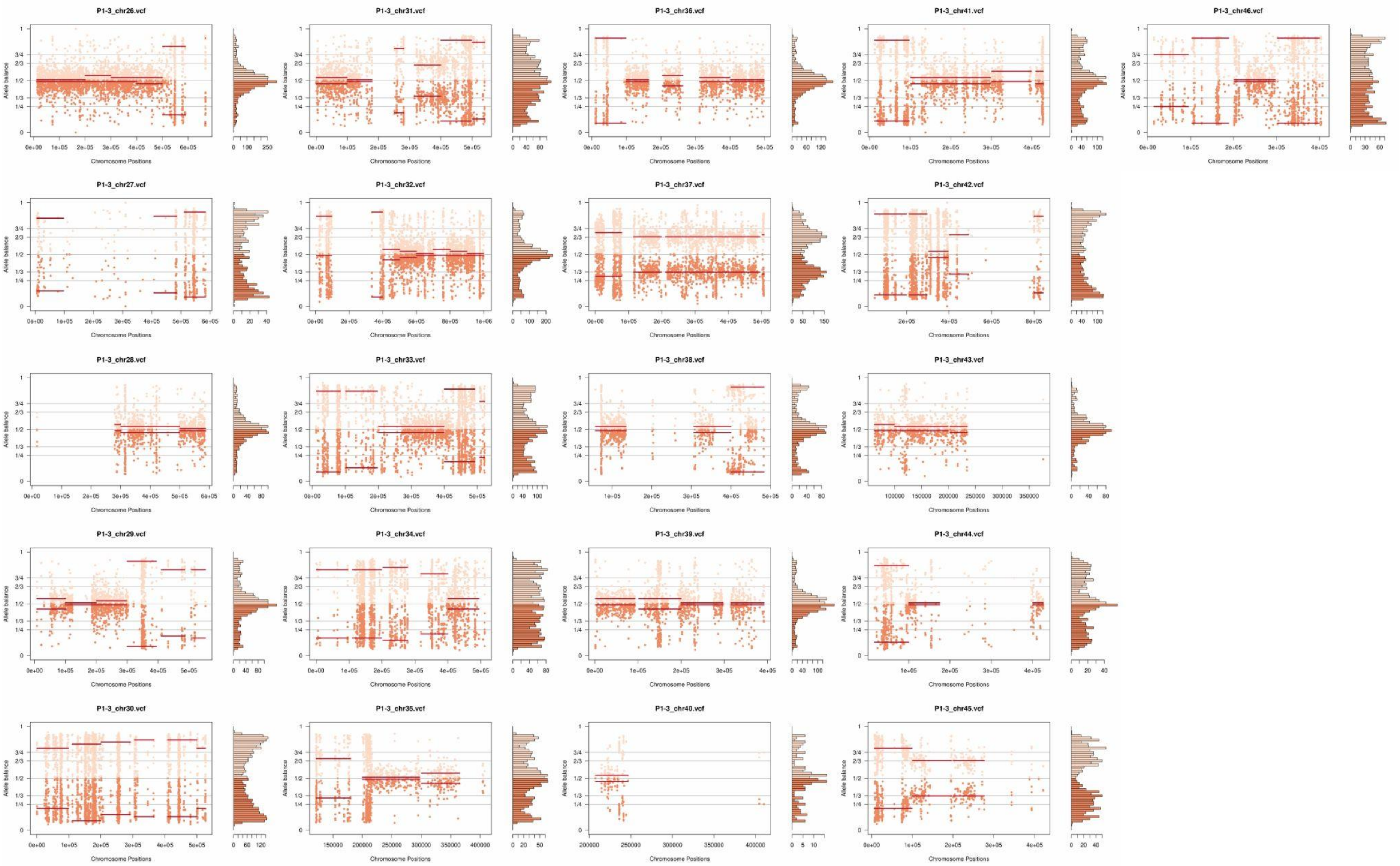
P1-3_chr26.vcf
P1-3_chr31.vcf
P1-3_chr36.vcf
P1-3_chr41.vcf
P1-3_chr46.vcf
P1-3_chr27.vcf
P1-3_chr32.vcf
P1-3_chr37.vcf
P1-3_chr42.vcf
P1-3_chr28.vcf
P1-3_chr33.vcf
P1-3_chr38.vcf
P1-3_chr43.vcf
P1-3_chr29.vcf
P1-3_chr34.vcf
P1-3_chr39.vcf
P1-3_chr44.vcf
P1-3_chr30.vcf
P1-3_chr35.vcf
P1-3_chr40.vcf
P1-3_chr45.vcf
Allele balance
Chromosome Positions

**Supplementary Fig. 7:** Somy estimation per chromosome based on AB confirming novel aneuploidies after *in vitro* culture in the three essentially diploid clone replicates of parental strain 2. Orange points represent the proportion of the alleles in heterozygous SNP positions along the chromosome. Darker orange represents the frequency of the first allele while lighter orange represents the frequency of the second allele, red lines represent the median value in windows.

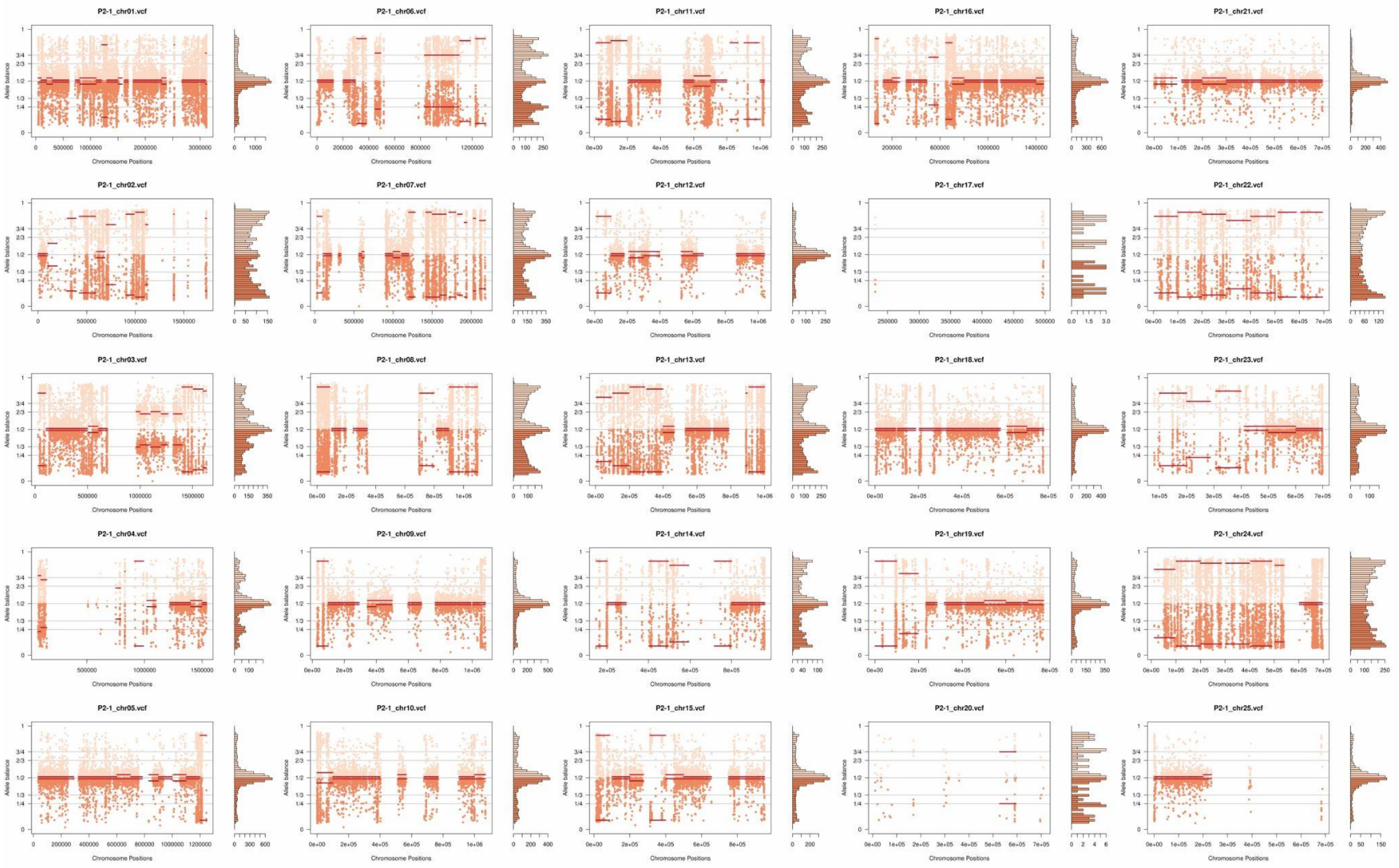

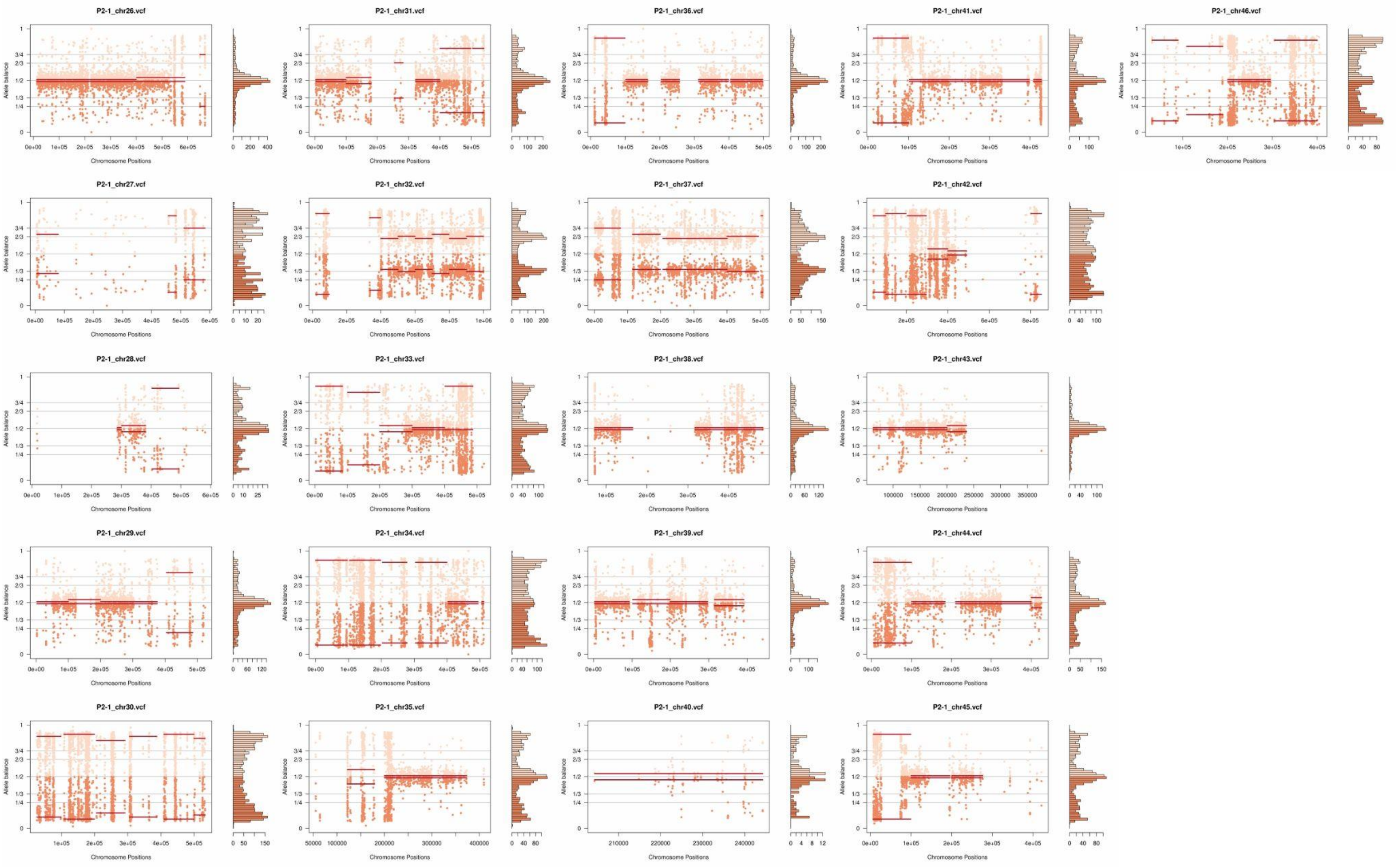
P2-1_chr26.vcf
P2-1_chr31.vcf
P2-1_chr36.vcf
P2-1_chr41.vcf
P2-1_chr46.vcf
P2-1_chr27.vcf
P2-1_chr32.vcf
P2-1_chr37.vcf
P2-1_chr42.vcf
P2-1_chr28.vcf
P2-1_chr33.vcf
P2-1_chr38.vcf
P2-1_chr43.vcf
P2-1_chr29.vcf
P2-1_chr34.vcf
P2-1_chr39.vcf
P2-1_chr44.vcf
P2-1_chr30.vcf
P2-1_chr35.vcf
P2-1_chr40.vcf
P2-1_chr45.vcf
Allele balance
Chromosome Positions

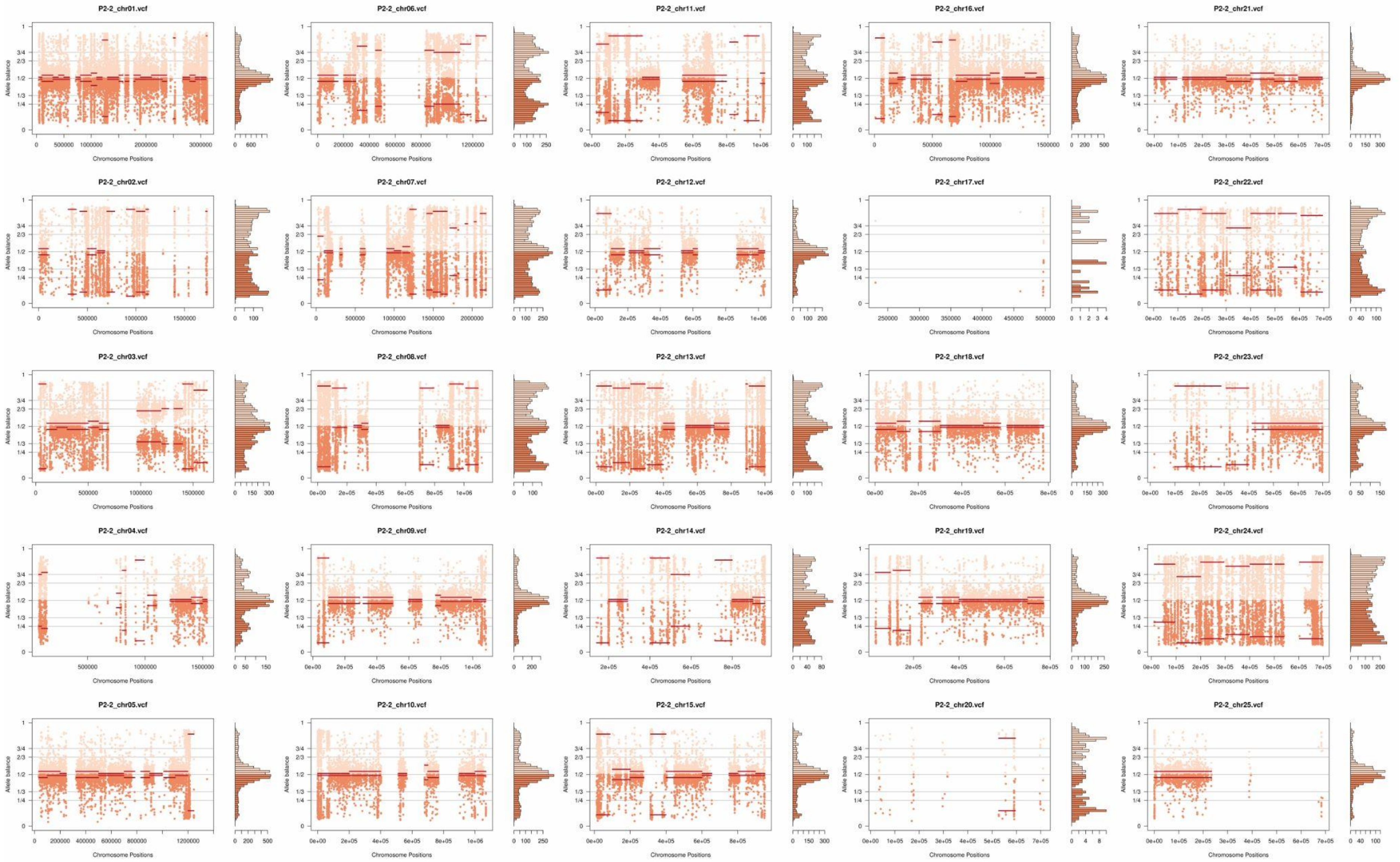
P2-2_chr01.vcf
P2-2_chr06.vcf
P2-2_chr11.vcf
P2-2_chr16.vcf
P2-2_chr21.vcf
P2-2_chr02.vcf
P2-2_chr07.vcf
P2-2_chr12.vcf
P2-2_chr17.vcf
P2-2_chr22.vcf
P2-2_chr03.vcf
P2-2_chr08.vcf
P2-2_chr13.vcf
P2-2_chr18.vcf
P2-2_chr23.vcf
P2-2_chr04.vcf
P2-2_chr09.vcf
P2-2_chr14.vcf
P2-2_chr19.vcf
P2-2_chr24.vcf
P2-2_chr05.vcf
P2-2_chr10.vcf
P2-2_chr15.vcf
P2-2_chr20.vcf
P2-2_chr25.vcf
Allele balance
Chromosome Positions

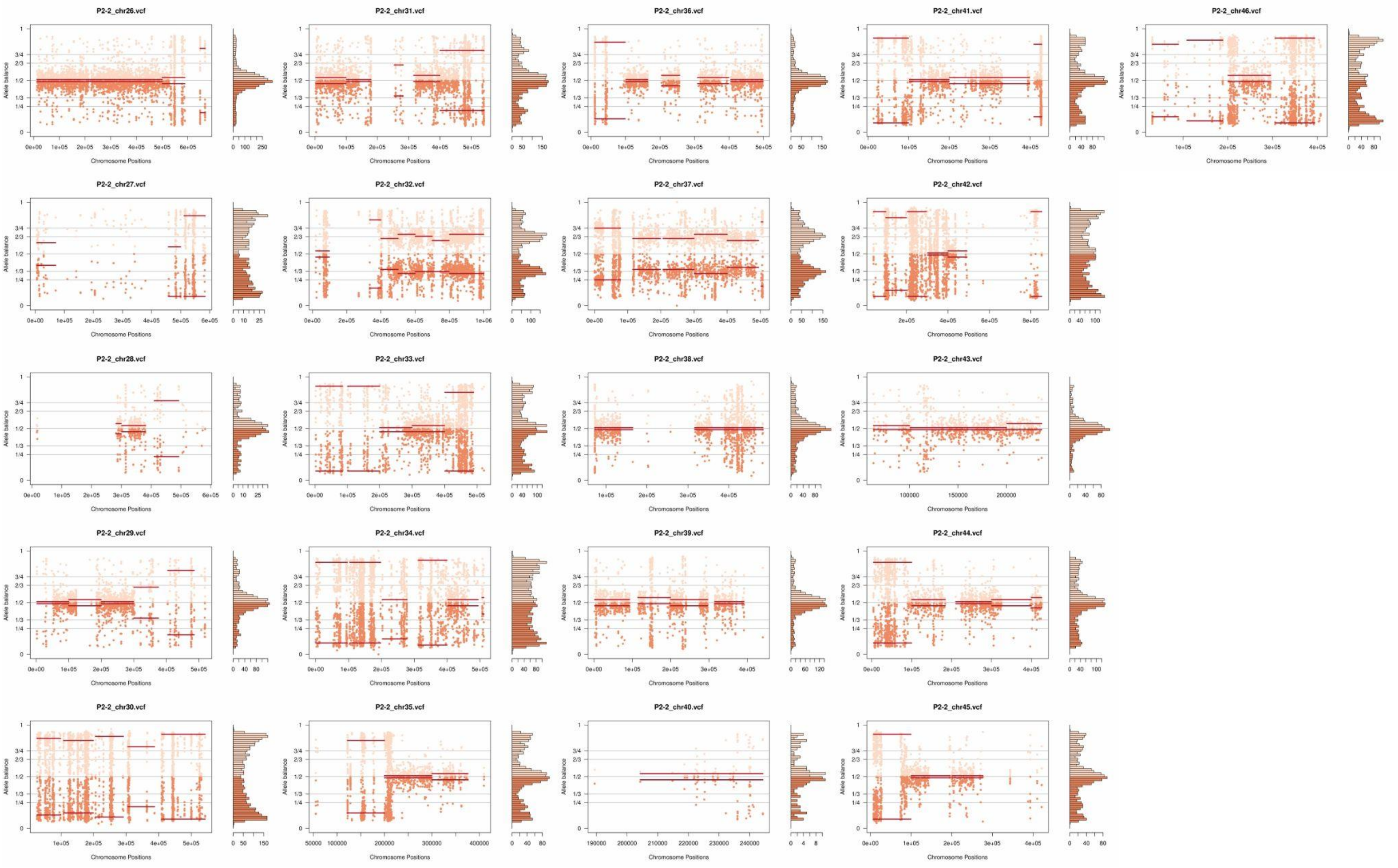
P2-2_chr26.vcf
P2-2_chr31.vcf
P2-2_chr36.vcf
P2-2_chr41.vcf
P2-2_chr46.vcf
P2-2_chr27.vcf
P2-2_chr32.vcf
P2-2_chr37.vcf
P2-2_chr42.vcf
P2-2_chr28.vcf
P2-2_chr33.vcf
P2-2_chr38.vcf
P2-2_chr43.vcf
P2-2_chr29.vcf
P2-2_chr34.vcf
P2-2_chr39.vcf
P2-2_chr44.vcf
P2-2_chr30.vcf
P2-2_chr35.vcf
P2-2_chr40.vcf
P2-2_chr45.vcf
Allele balance
Chromosome Positions

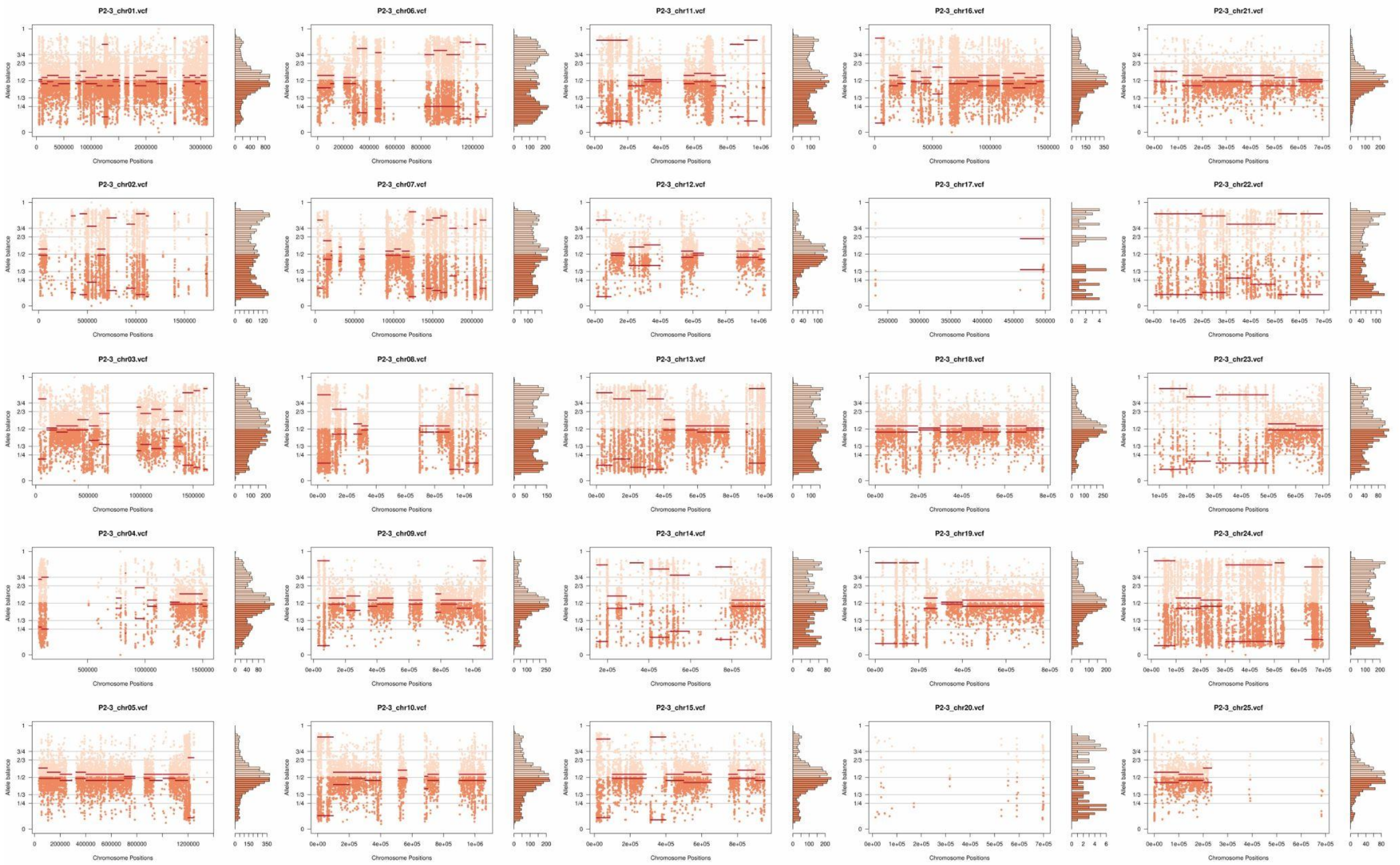
P2-3_chr01.vcf
P2-3_chr06.vcf
P2-3_chr11.vcf
P2-3_chr16.vcf
P2-3_chr21.vcf
P2-3_chr02.vcf
P2-3_chr07.vcf
P2-3_chr12.vcf
P2-3_chr17.vcf
P2-3_chr22.vcf
P2-3_chr03.vcf
P2-3_chr08.vcf
P2-3_chr13.vcf
P2-3_chr18.vcf
P2-3_chr23.vcf
P2-3_chr04.vcf
P2-3_chr09.vcf
P2-3_chr14.vcf
P2-3_chr19.vcf
P2-3_chr24.vcf
P2-3_chr05.vcf
P2-3_chr10.vcf
P2-3_chr15.vcf
P2-3_chr20.vcf
P2-3_chr25.vcf
Allele balance
Chromosome Positions

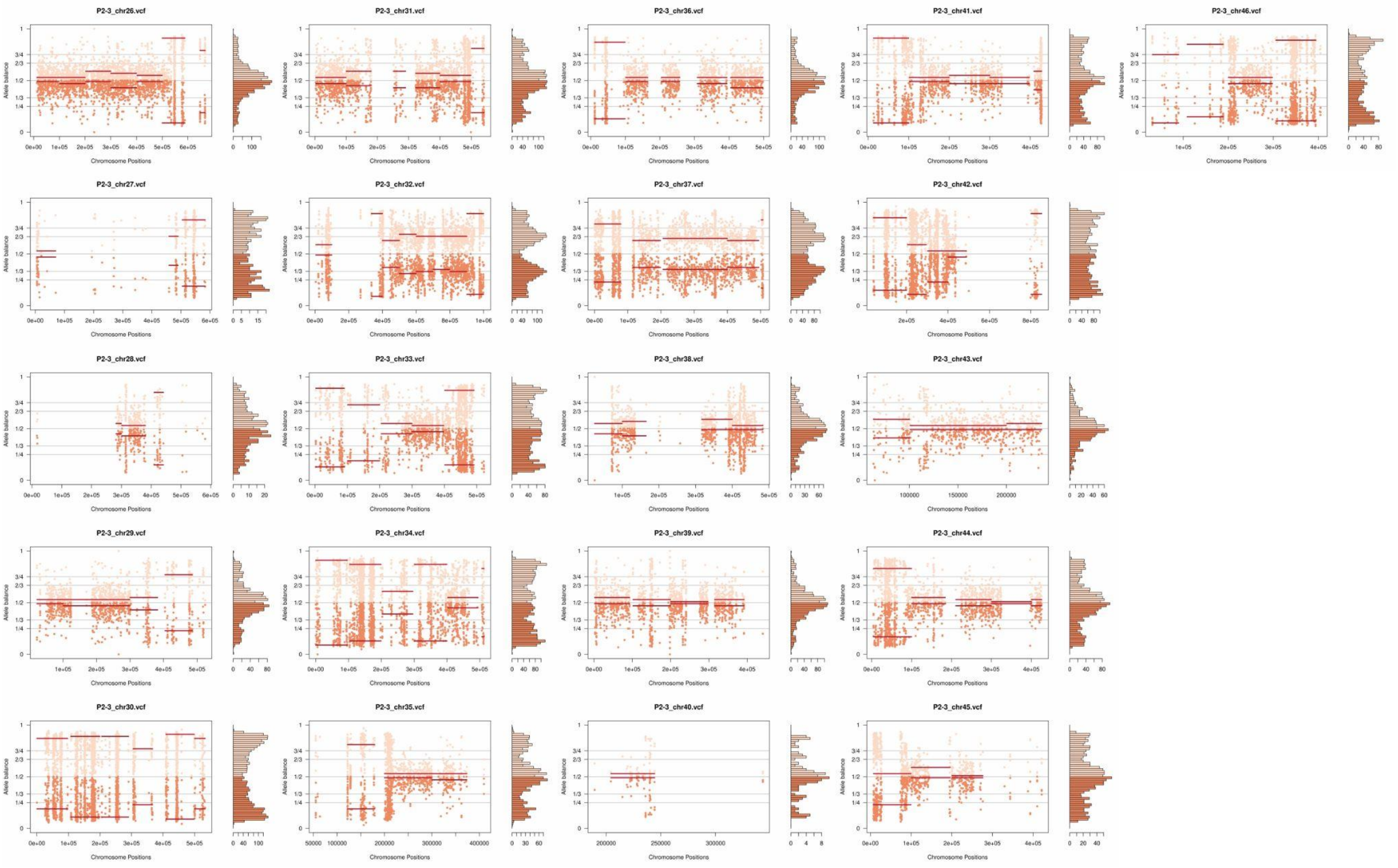
P2-3_chr26.vcf
P2-3_chr31.vcf
P2-3_chr36.vcf
P2-3_chr41.vcf
P2-3_chr46.vcf
P2-3_chr27.vcf
P2-3_chr32.vcf
P2-3_chr37.vcf
P2-3_chr42.vcf
P2-3_chr28.vcf
P2-3_chr33.vcf
P2-3_chr38.vcf
P2-3_chr43.vcf
P2-3_chr29.vcf
P2-3_chr34.vcf
P2-3_chr39.vcf
P2-3_chr44.vcf
P2-3_chr30.vcf
P2-3_chr35.vcf
P2-3_chr40.vcf
P2-3_chr45.vcf
Allele balance
Chromosome Positions

**Supplementary Fig. 8:** Somy estimation per chromosome based on AB confirming trisomic and tetrasomic chromosomes after *in vitro* culture in the three clone replicates of hybrid 1C2. Orange points represent the proportion of the alleles in heterozygous SNP positions along the chromosome. Darker orange represents the frequency of the first allele while lighter orange represents the frequency of the second allele, red lines represent the median value in windows.

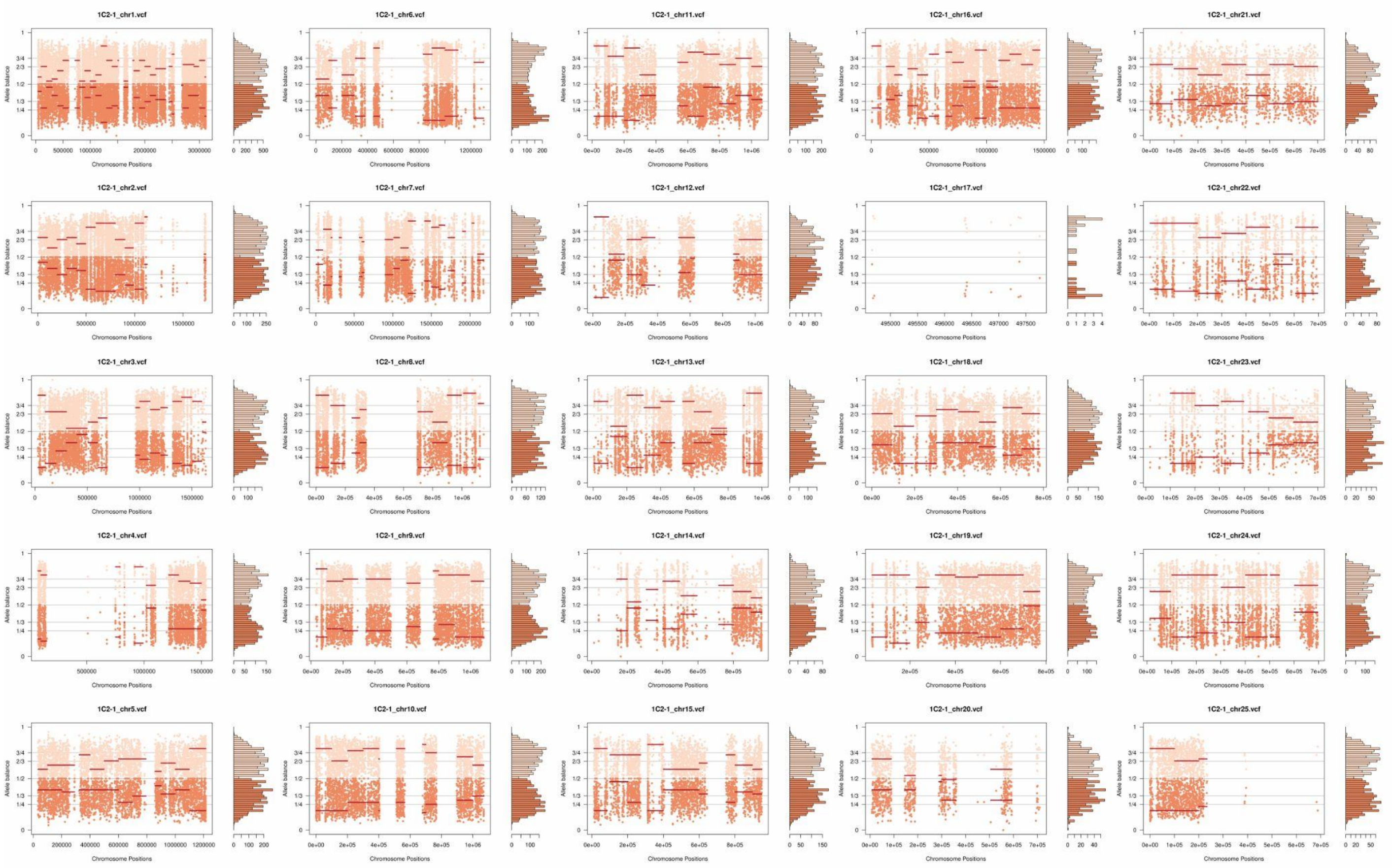

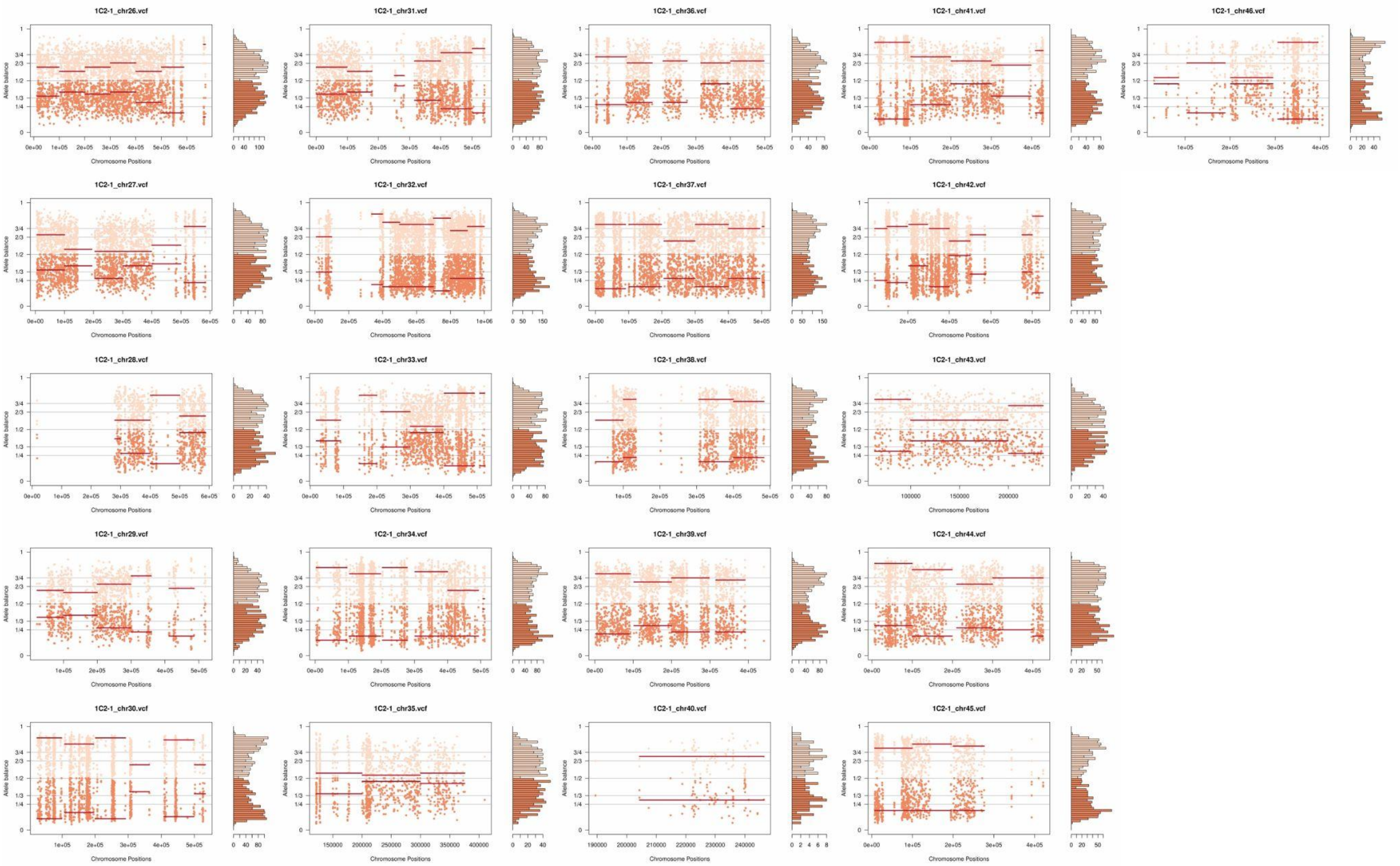
1C2-1_chr26.vcf
1C2-1_chr31.vcf
1C2-1_chr36.vcf
1C2-1_chr41.vcf
1C2-1_chr46.vcf
1C2-1_chr27.vcf
1C2-1_chr32.vcf
1C2-1_chr37.vcf
1C2-1_chr42.vcf
1C2-1_chr28.vcf
1C2-1_chr33.vcf
1C2-1_chr38.vcf
1C2-1_chr43.vcf
1C2-1_chr29.vcf
1C2-1_chr34.vcf
1C2-1_chr39.vcf
1C2-1_chr44.vcf
1C2-1_chr30.vcf
1C2-1_chr35.vcf
1C2-1_chr40.vcf
1C2-1_chr45.vcf
Allele balance
Chromosome Positions

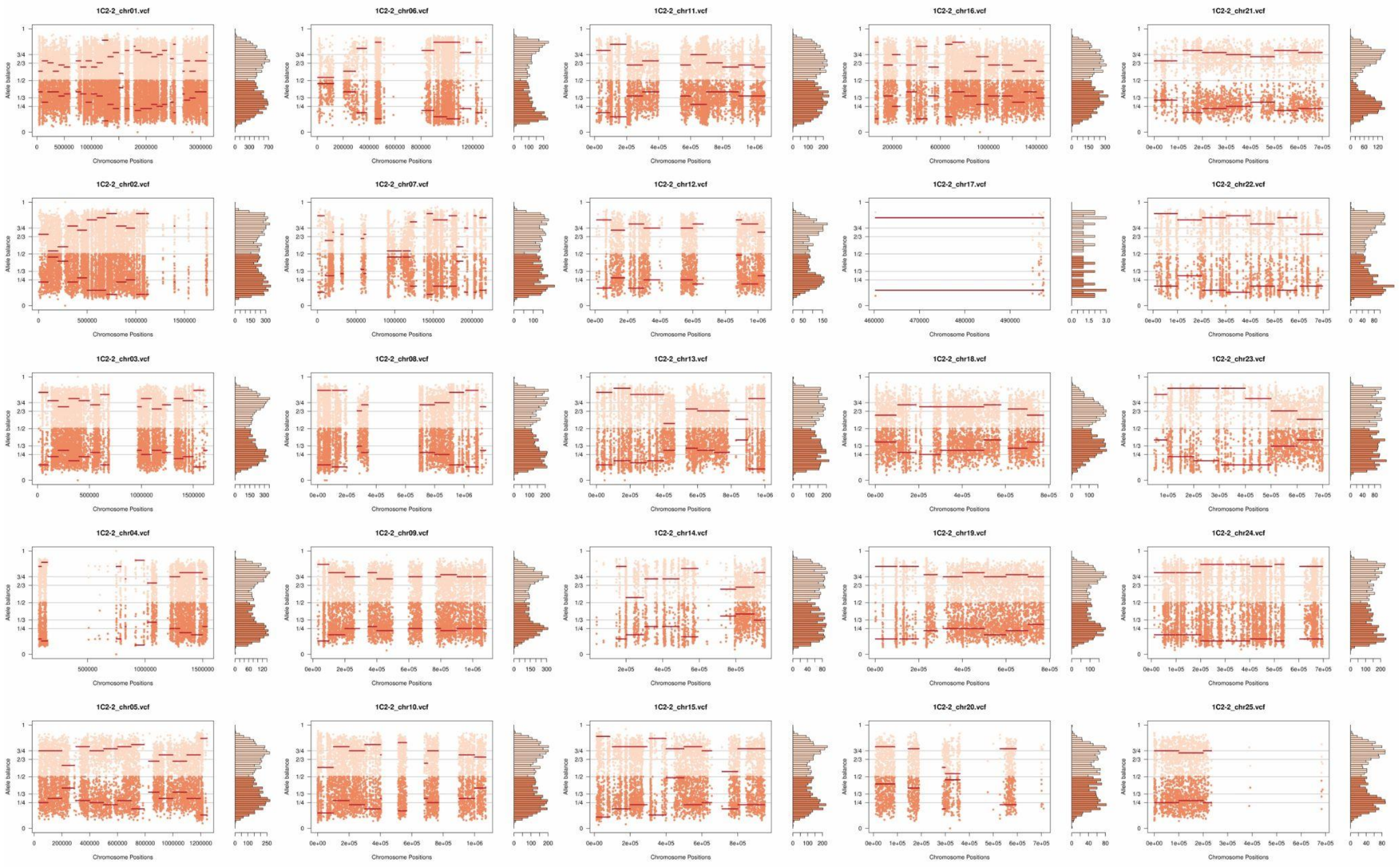
1C2-2_chr01.vcf
1C2-2_chr06.vcf
1C2-2_chr11.vcf
1C2-2_chr16.vcf
1C2-2_chr21.vcf
1C2-2_chr02.vcf
1C2-2_chr07.vcf
1C2-2_chr12.vcf
1C2-2_chr17.vcf
1C2-2_chr22.vcf
1C2-2_chr03.vcf
1C2-2_chr08.vcf
1C2-2_chr13.vcf
1C2-2_chr18.vcf
1C2-2_chr23.vcf
1C2-2_chr04.vcf
1C2-2_chr09.vcf
1C2-2_chr14.vcf
1C2-2_chr19.vcf
1C2-2_chr24.vcf
1C2-2_chr05.vcf
1C2-2_chr10.vcf
1C2-2_chr15.vcf
1C2-2_chr20.vcf
1C2-2_chr25.vcf
Allele balance
Chromosome Positions

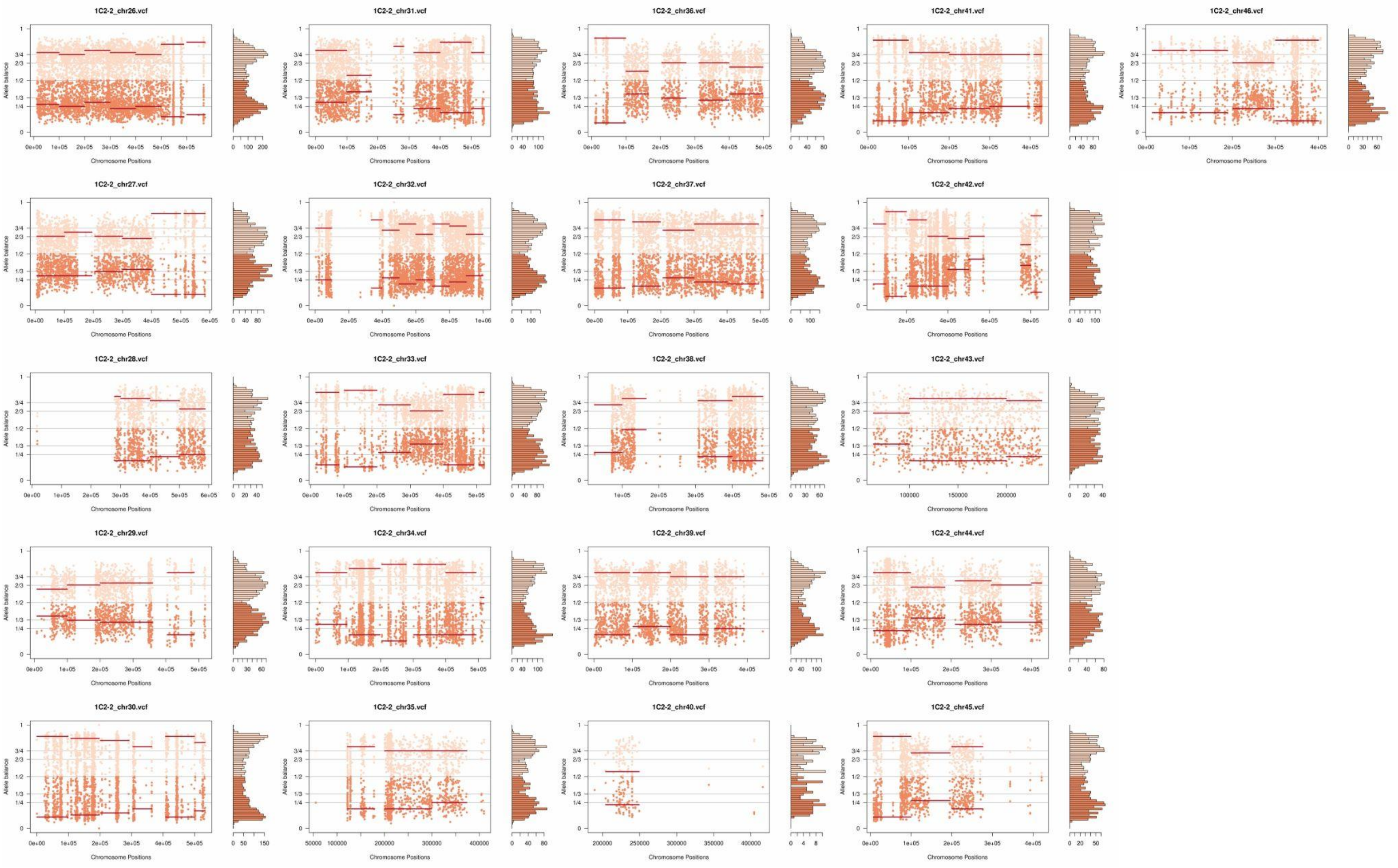
1C2-2_chr26.vcf
1C2-2_chr31.vcf
1C2-2_chr36.vcf
1C2-2_chr41.vcf
1C2-2_chr46.vcf
1C2-2_chr27.vcf
1C2-2_chr32.vcf
1C2-2_chr37.vcf
1C2-2_chr42.vcf
1C2-2_chr28.vcf
1C2-2_chr33.vcf
1C2-2_chr38.vcf
1C2-2_chr43.vcf
1C2-2_chr29.vcf
1C2-2_chr34.vcf
1C2-2_chr39.vcf
1C2-2_chr44.vcf
1C2-2_chr30.vcf
1C2-2_chr35.vcf
1C2-2_chr40.vcf
1C2-2_chr45.vcf
Allele balance
Chromosome Positions

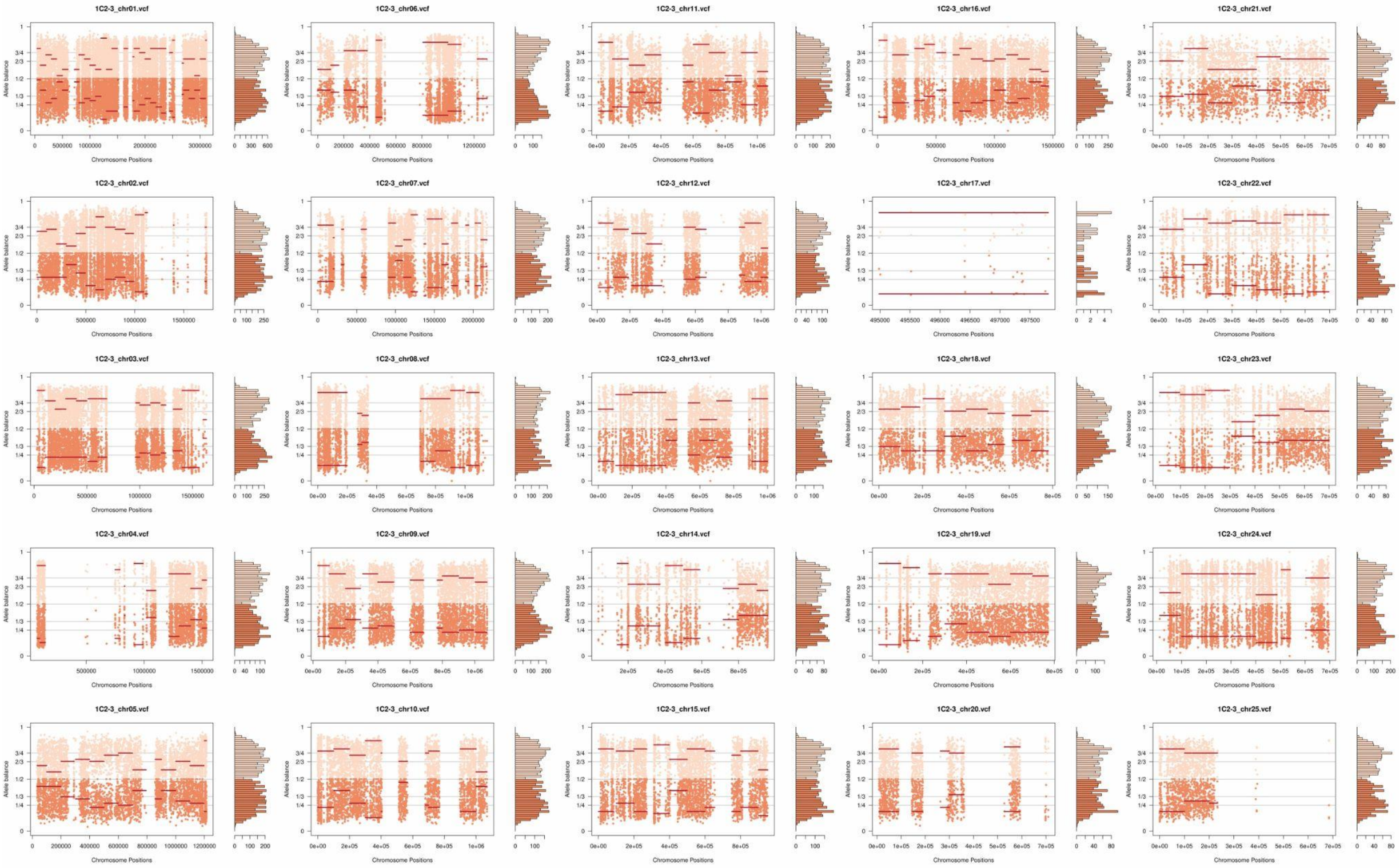
1C2-3_chr01.vcf
1C2-3_chr06.vcf
1C2-3_chr11.vcf
1C2-3_chr16.vcf
1C2-3_chr21.vcf
1C2-3_chr02.vcf
1C2-3_chr07.vcf
1C2-3_chr12.vcf
1C2-3_chr17.vcf
1C2-3_chr22.vcf
1C2-3_chr03.vcf
1C2-3_chr08.vcf
1C2-3_chr13.vcf
1C2-3_chr18.vcf
1C2-3_chr23.vcf
1C2-3_chr04.vcf
1C2-3_chr09.vcf
1C2-3_chr14.vcf
1C2-3_chr19.vcf
1C2-3_chr24.vcf
1C2-3_chr05.vcf
1C2-3_chr10.vcf
1C2-3_chr15.vcf
1C2-3_chr20.vcf
1C2-3_chr25.vcf
Allele balance
Chromosome Positions

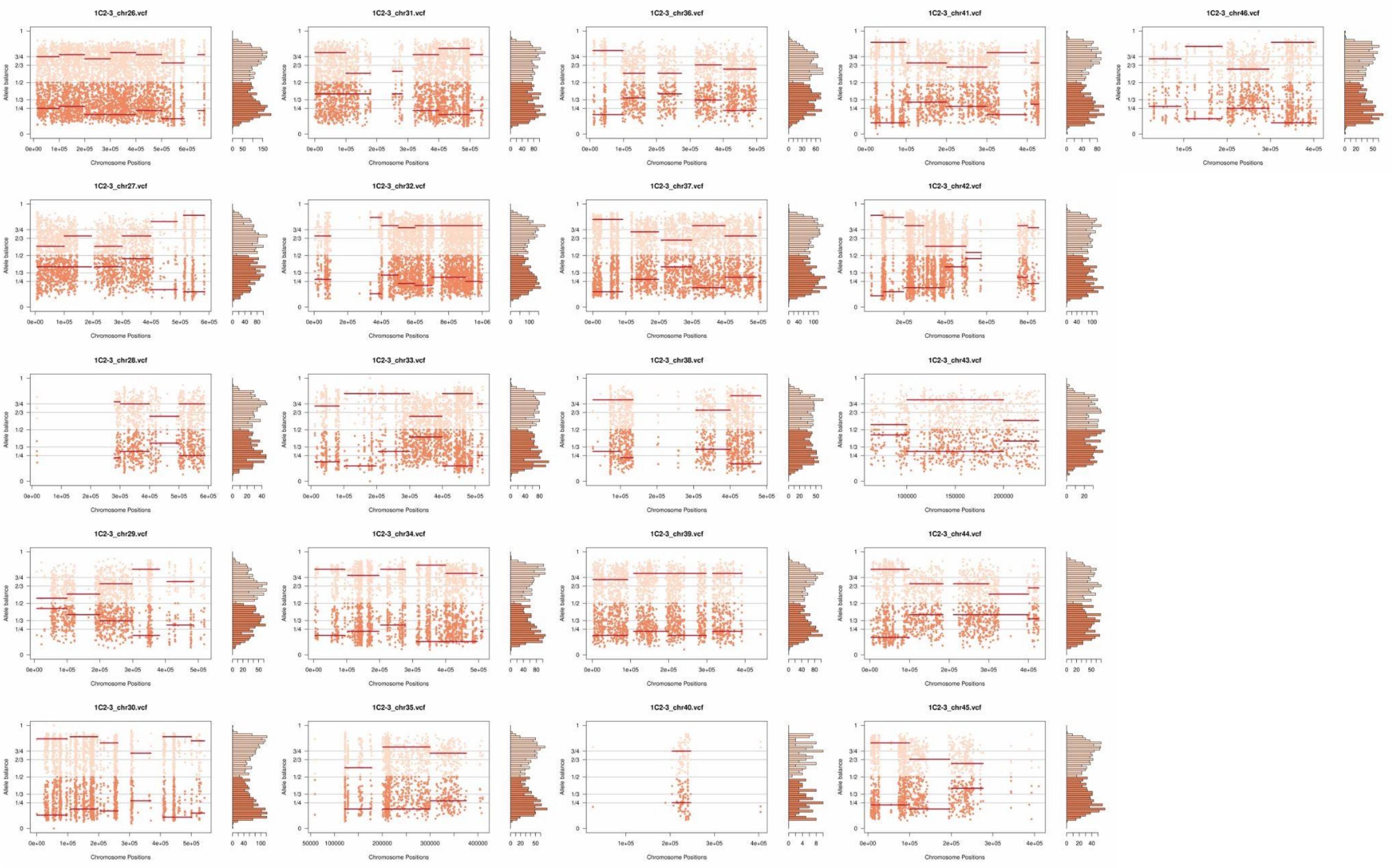
1C2-3_chr26.vcf
1C2-3_chr31.vcf
1C2-3_chr36.vcf
1C2-3_chr41.vcf
1C2-3_chr46.vcf
1C2-3_chr27.vcf
1C2-3_chr32.vcf
1C2-3_chr37.vcf
1C2-3_chr42.vcf
1C2-3_chr28.vcf
1C2-3_chr33.vcf
1C2-3_chr38.vcf
1C2-3_chr43.vcf
1C2-3_chr29.vcf
1C2-3_chr34.vcf
1C2-3_chr39.vcf
1C2-3_chr44.vcf
1C2-3_chr30.vcf
1C2-3_chr35.vcf
1C2-3_chr40.vcf
1C2-3_chr45.vcf
Allele balance
Chromosome Positions

**Supplementary Fig. 9:** Somy estimation per chromosome based on AB confirming trisomic and tetrasomic chromosomes after *in vitro* culture in the three clone replicates of hybrid 1D12. Orange points represent the proportion of the alleles in heterozygous SNP positions along the chromosome. Darker orange represents the frequency of the first allele while lighter orange represents the frequency of the second allele, red lines represent the median value in windows.

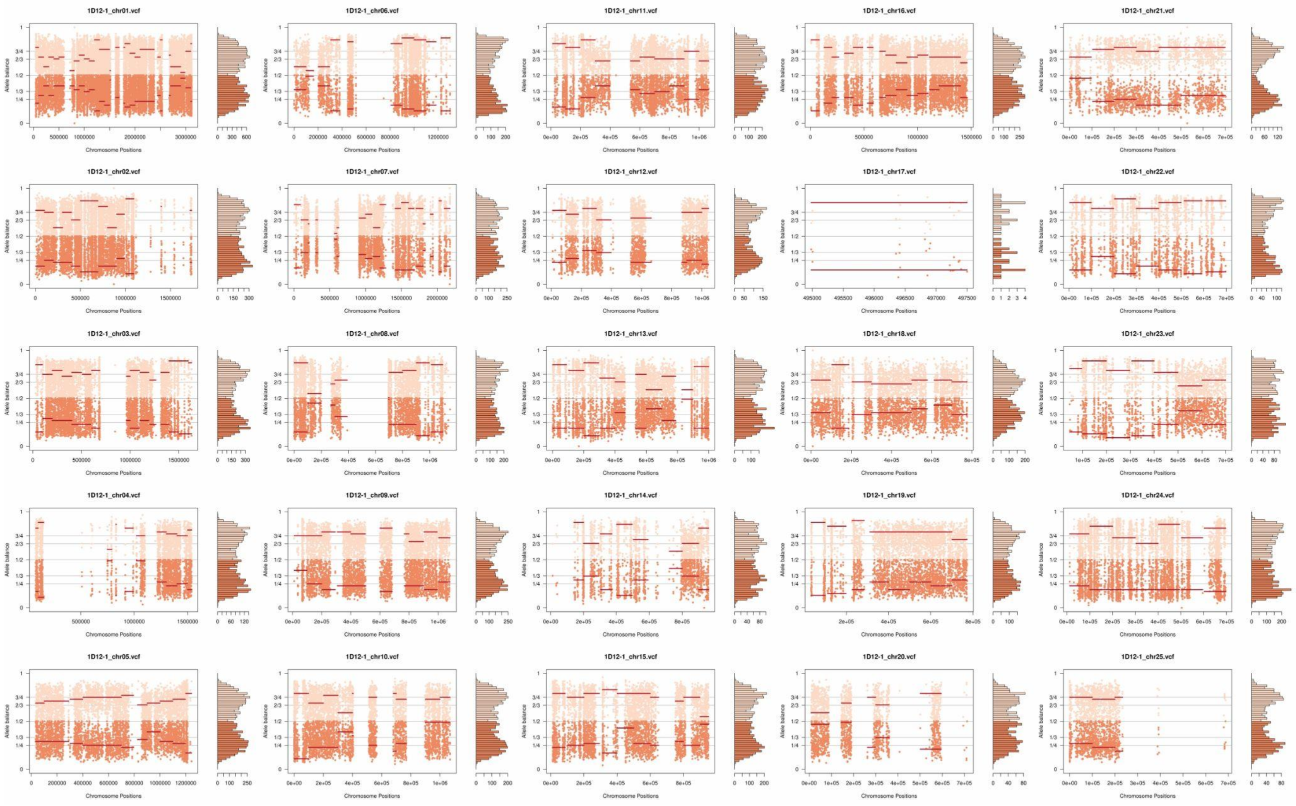

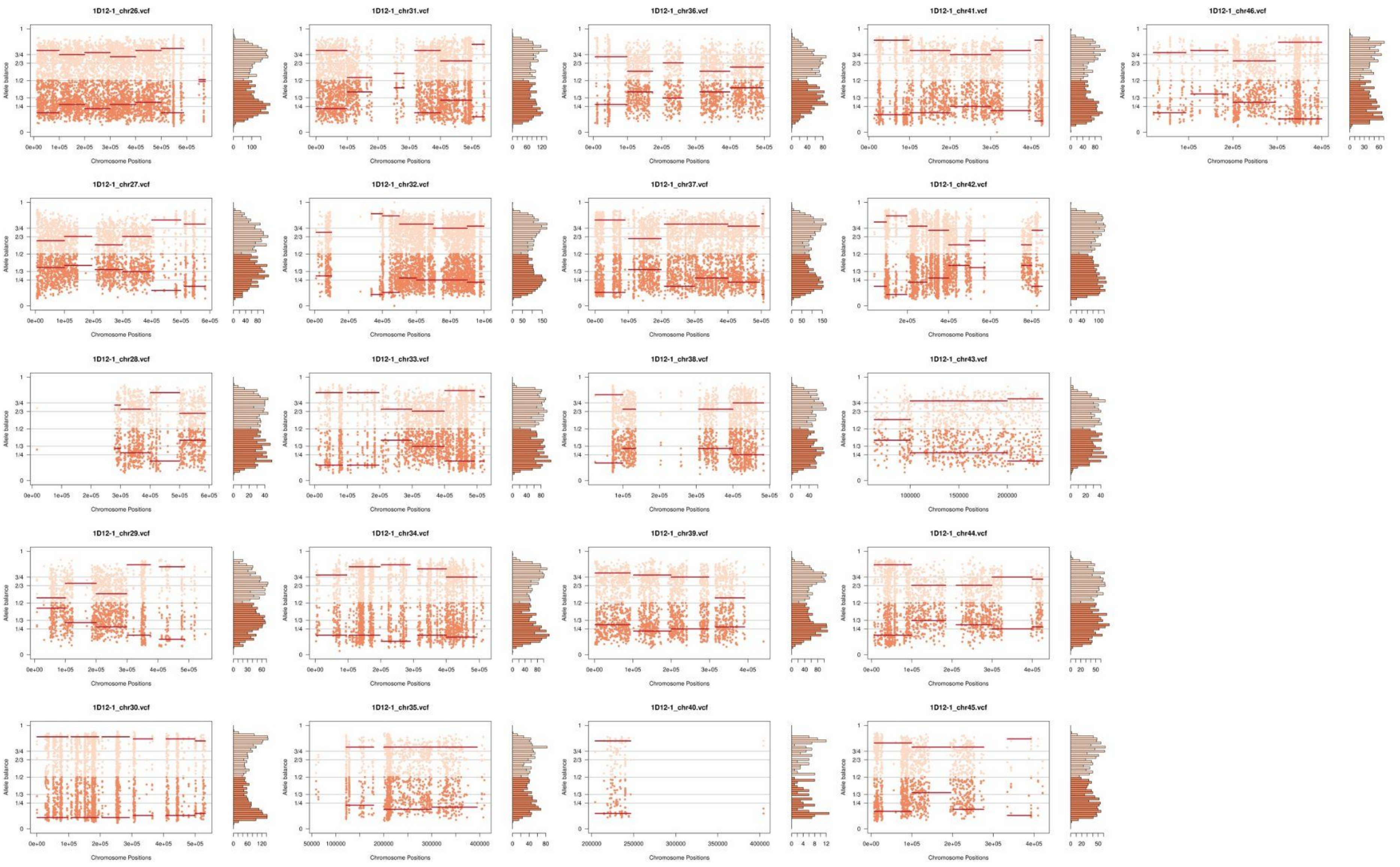
1D12-1_chr26.vcf
1D12-1_chr31.vcf
1D12-1_chr36.vcf
1D12-1_chr41.vcf
1D12-1_chr46.vcf
1D12-1_chr27.vcf
1D12-1_chr32.vcf
1D12-1_chr37.vcf
1D12-1_chr42.vcf
1D12-1_chr28.vcf
1D12-1_chr33.vcf
1D12-1_chr38.vcf
1D12-1_chr43.vcf
1D12-1_chr29.vcf
1D12-1_chr34.vcf
1D12-1_chr39.vcf
1D12-1_chr44.vcf
1D12-1_chr30.vcf
1D12-1_chr35.vcf
1D12-1_chr40.vcf
1D12-1_chr45.vcf
Allele balance
Chromosome Positions

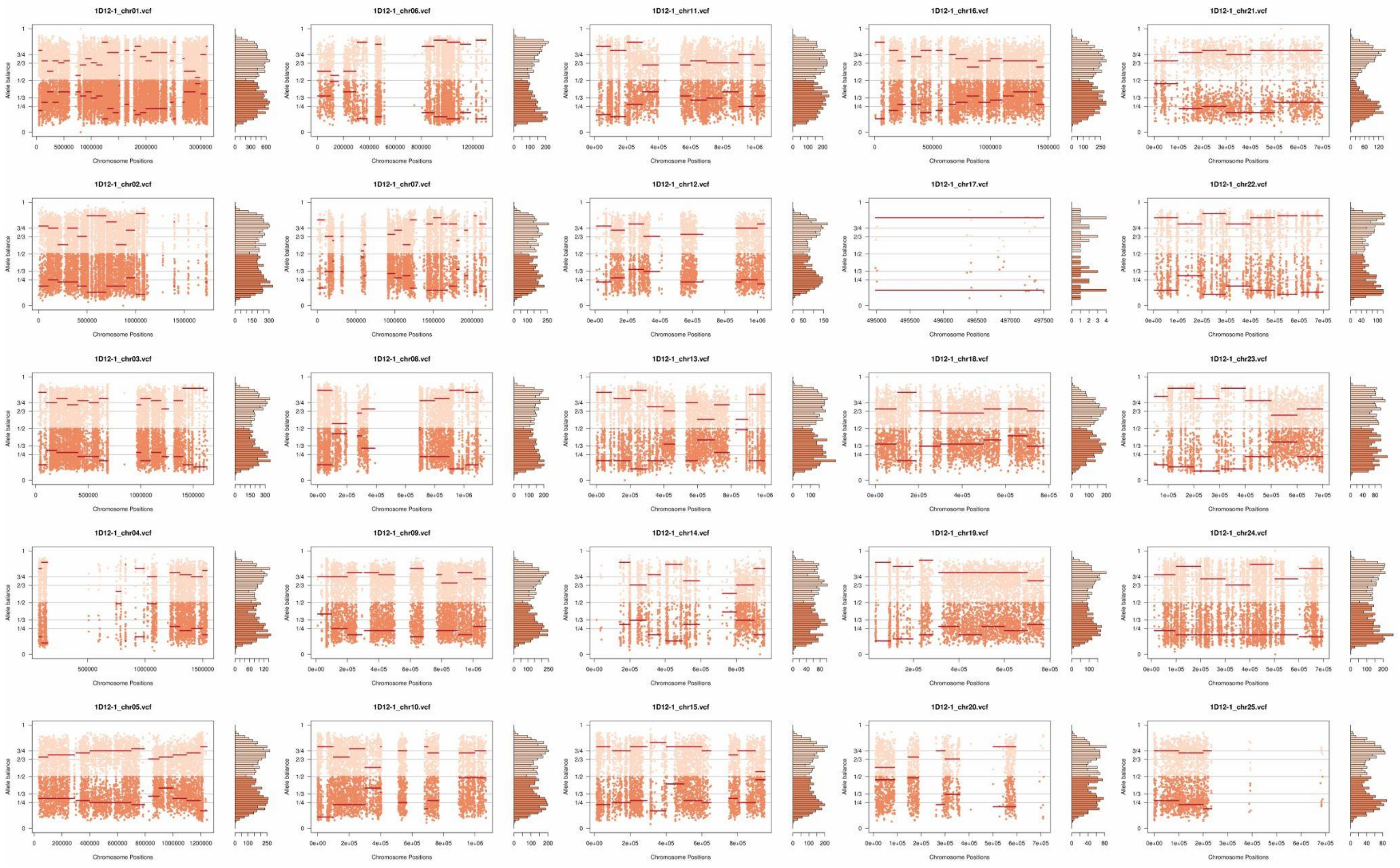
1D12-1_chr01.vcf
1D12-1_chr06.vcf
1D12-1_chr11.vcf
1D12-1_chr16.vcf
1D12-1_chr21.vcf
1D12-1_chr02.vcf
1D12-1_chr07.vcf
1D12-1_chr12.vcf
1D12-1_chr17.vcf
1D12-1_chr22.vcf
1D12-1_chr03.vcf
1D12-1_chr08.vcf
1D12-1_chr13.vcf
1D12-1_chr18.vcf
1D12-1_chr23.vcf
1D12-1_chr04.vcf
1D12-1_chr09.vcf
1D12-1_chr14.vcf
1D12-1_chr19.vcf
1D12-1_chr24.vcf
1D12-1_chr05.vcf
1D12-1_chr10.vcf
1D12-1_chr15.vcf
1D12-1_chr20.vcf
1D12-1_chr25.vcf
Allele balance
Chromosome Positions

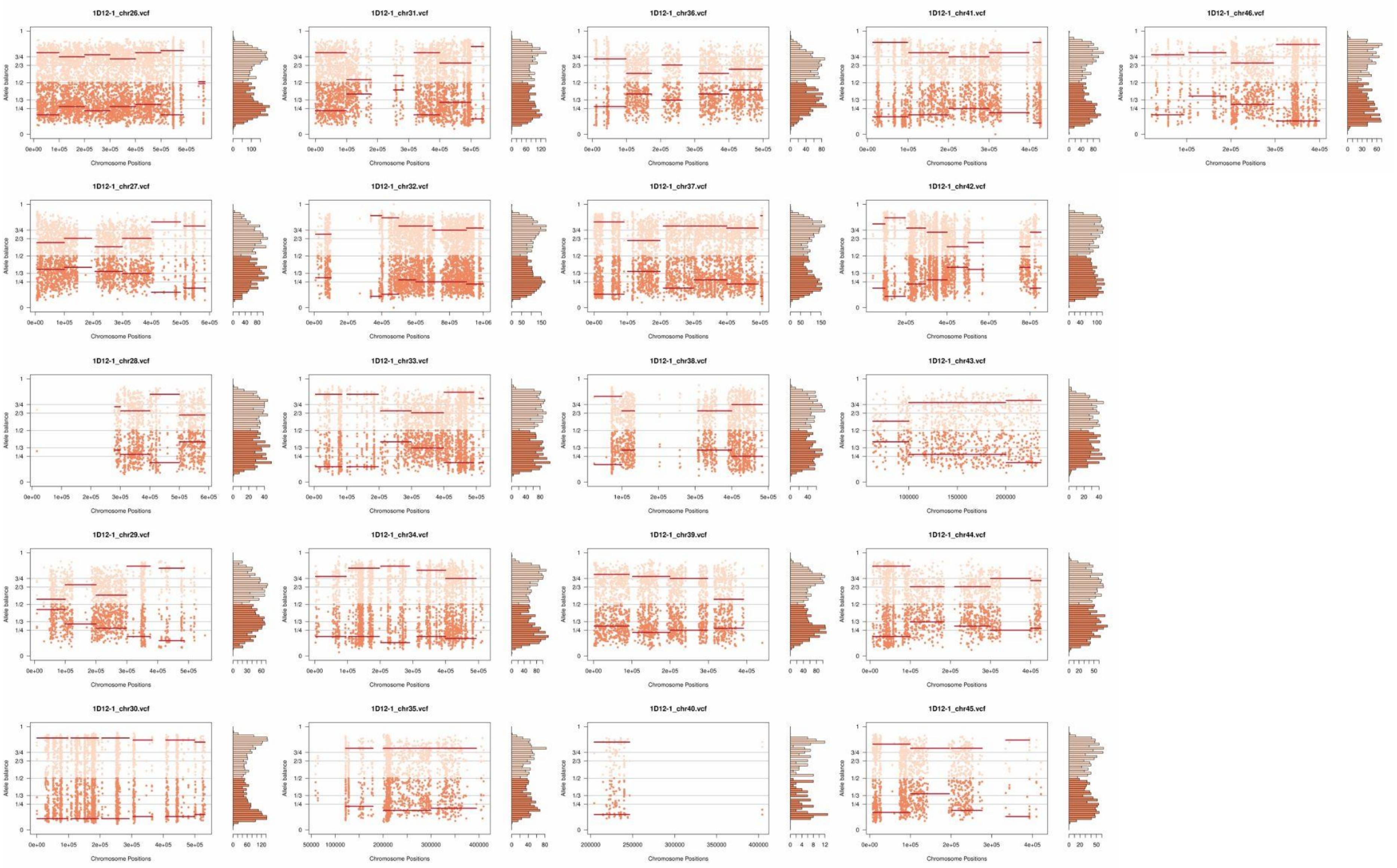
1D12-1_chr26.vcf
1D12-1_chr31.vcf
1D12-1_chr36.vcf
1D12-1_chr41.vcf
1D12-1_chr46.vcf
1D12-1_chr27.vcf
1D12-1_chr32.vcf
1D12-1_chr37.vcf
1D12-1_chr42.vcf
1D12-1_chr28.vcf
1D12-1_chr33.vcf
1D12-1_chr38.vcf
1D12-1_chr43.vcf
1D12-1_chr29.vcf
1D12-1_chr34.vcf
1D12-1_chr39.vcf
1D12-1_chr44.vcf
1D12-1_chr30.vcf
1D12-1_chr35.vcf
1D12-1_chr40.vcf
1D12-1_chr45.vcf
Allele balance
Chromosome Positions

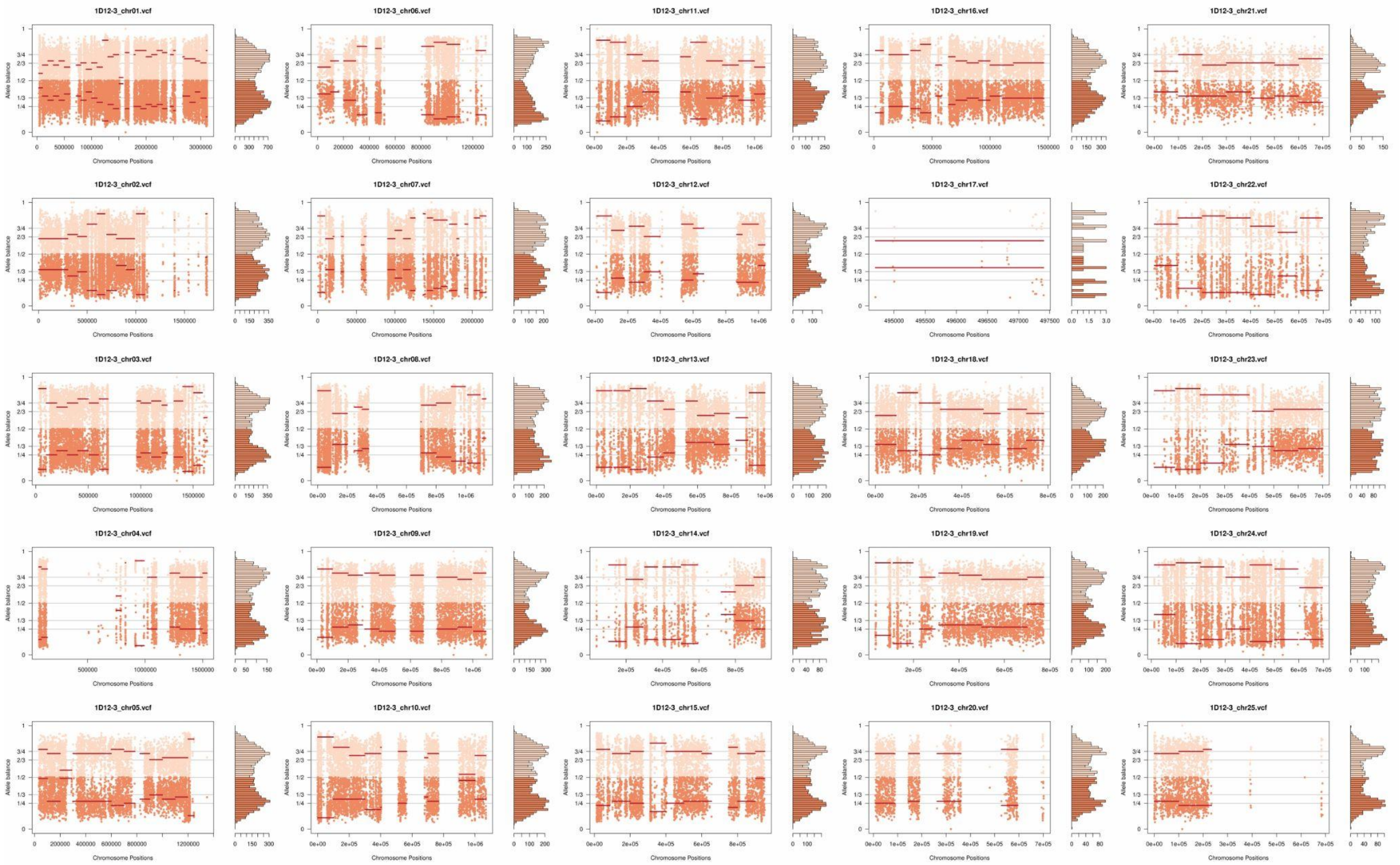
1D12-3_chr01.vcf
1D12-3_chr06.vcf
1D12-3_chr11.vcf
1D12-3_chr16.vcf
1D12-3_chr21.vcf
1D12-3_chr02.vcf
1D12-3_chr07.vcf
1D12-3_chr12.vcf
1D12-3_chr17.vcf
1D12-3_chr22.vcf
1D12-3_chr03.vcf
1D12-3_chr08.vcf
1D12-3_chr13.vcf
1D12-3_chr18.vcf
1D12-3_chr23.vcf
1D12-3_chr04.vcf
1D12-3_chr09.vcf
1D12-3_chr14.vcf
1D12-3_chr19.vcf
1D12-3_chr24.vcf
1D12-3_chr05.vcf
1D12-3_chr10.vcf
1D12-3_chr15.vcf
1D12-3_chr20.vcf
1D12-3_chr25.vcf
Allele balance
Chromosome Positions

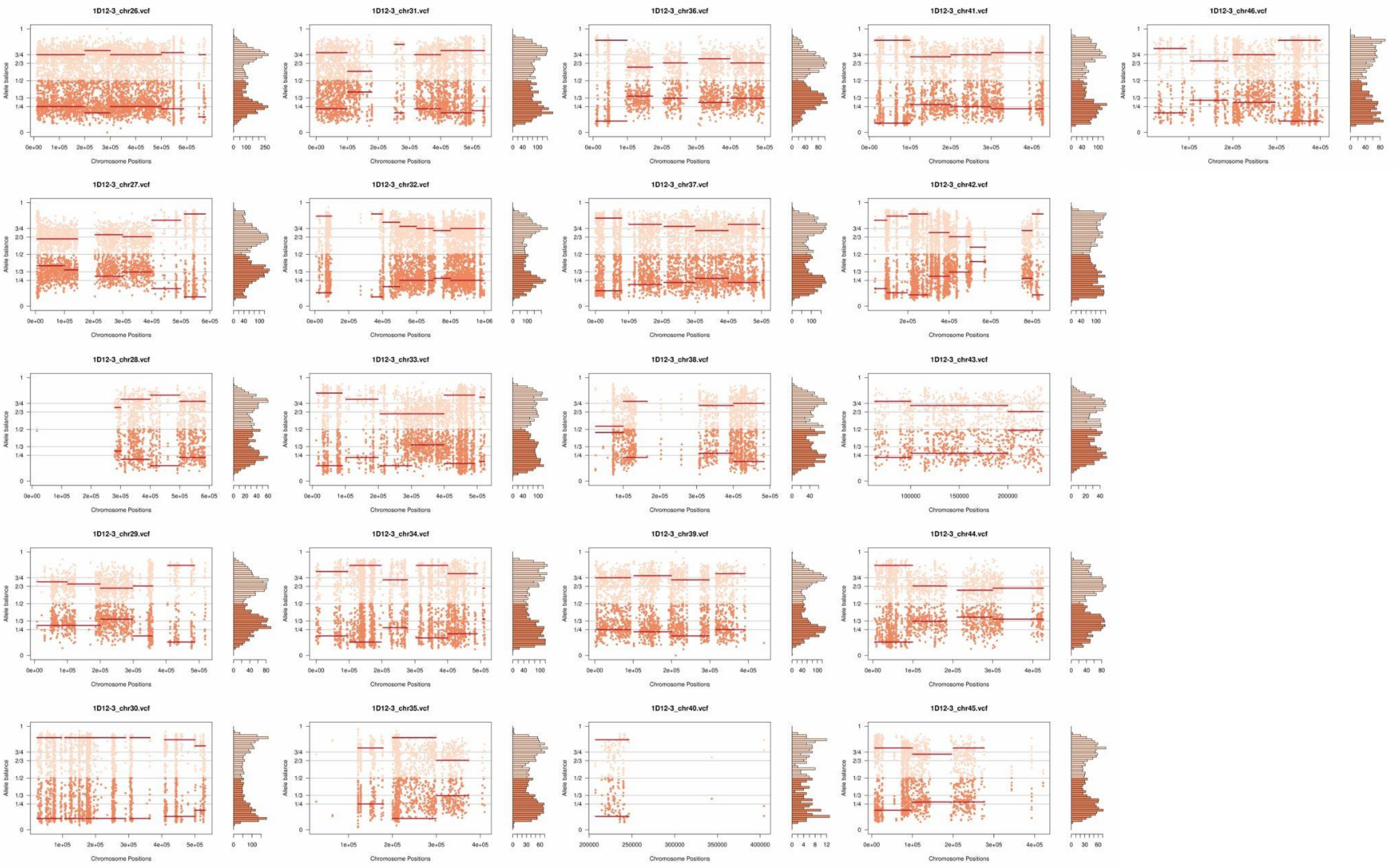
1D12-3_chr26.vcf
1D12-3_chr31.vcf
1D12-3_chr36.vcf
1D12-3_chr41.vcf
1D12-3_chr46.vcf
1D12-3_chr27.vcf
1D12-3_chr32.vcf
1D12-3_chr37.vcf
1D12-3_chr42.vcf
1D12-3_chr28.vcf
1D12-3_chr33.vcf
1D12-3_chr38.vcf
1D12-3_chr43.vcf
1D12-3_chr29.vcf
1D12-3_chr34.vcf
1D12-3_chr39.vcf
1D12-3_chr44.vcf
1D12-3_chr30.vcf
1D12-3_chr35.vcf
1D12-3_chr40.vcf
1D12-3_chr45.vcf
Allele balance
Chromosome Positions

**Supplementary Fig. 10:** Somy estimation per chromosome based on AB confirming trisomic and tetrasomic chromosomes after *in vitro* culture in the three clone replicates of hybrid 2C1. Orange points represent the proportion of the alleles in heterozygous SNP positions along the chromosome. Darker orange represents the frequency of the first allele while lighter orange represents the frequency of the second allele, red lines represent the median value in windows.

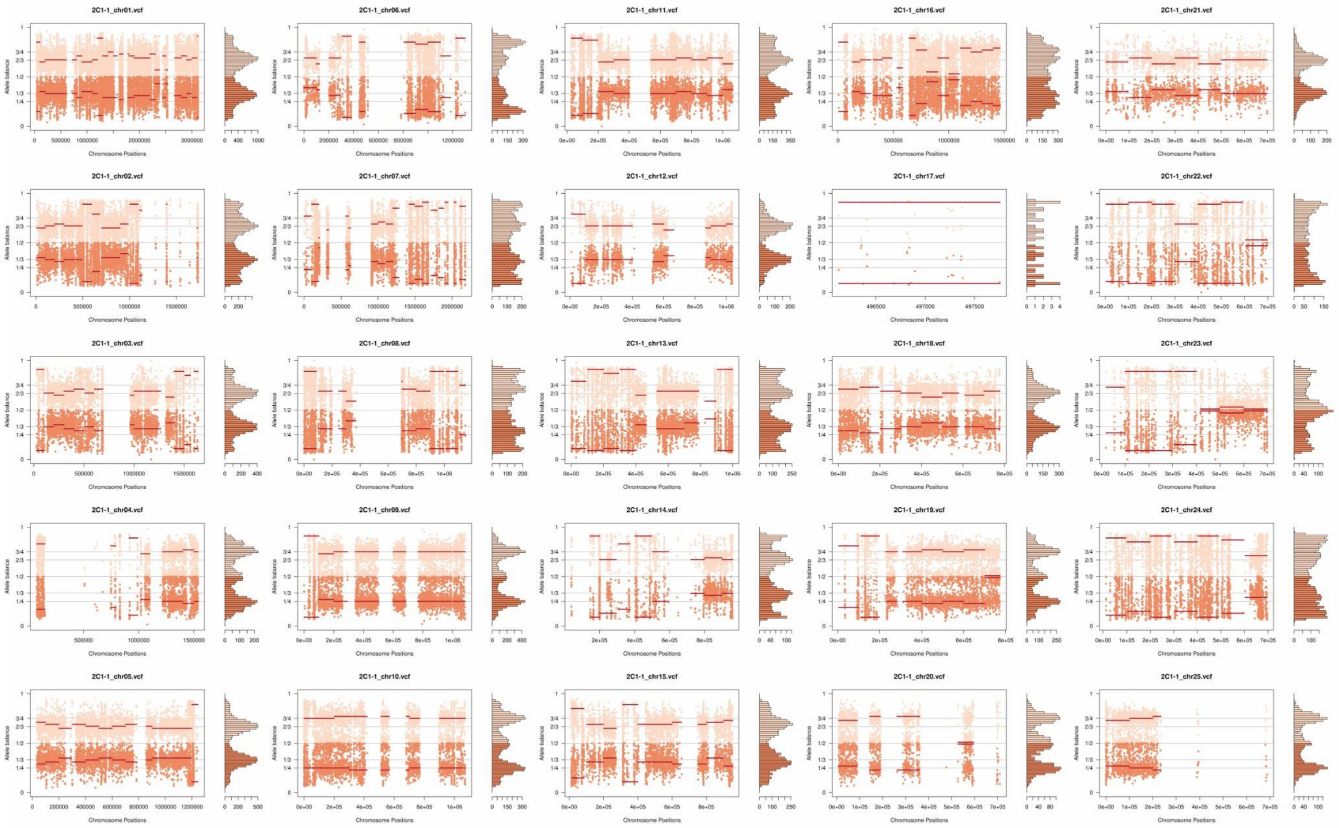

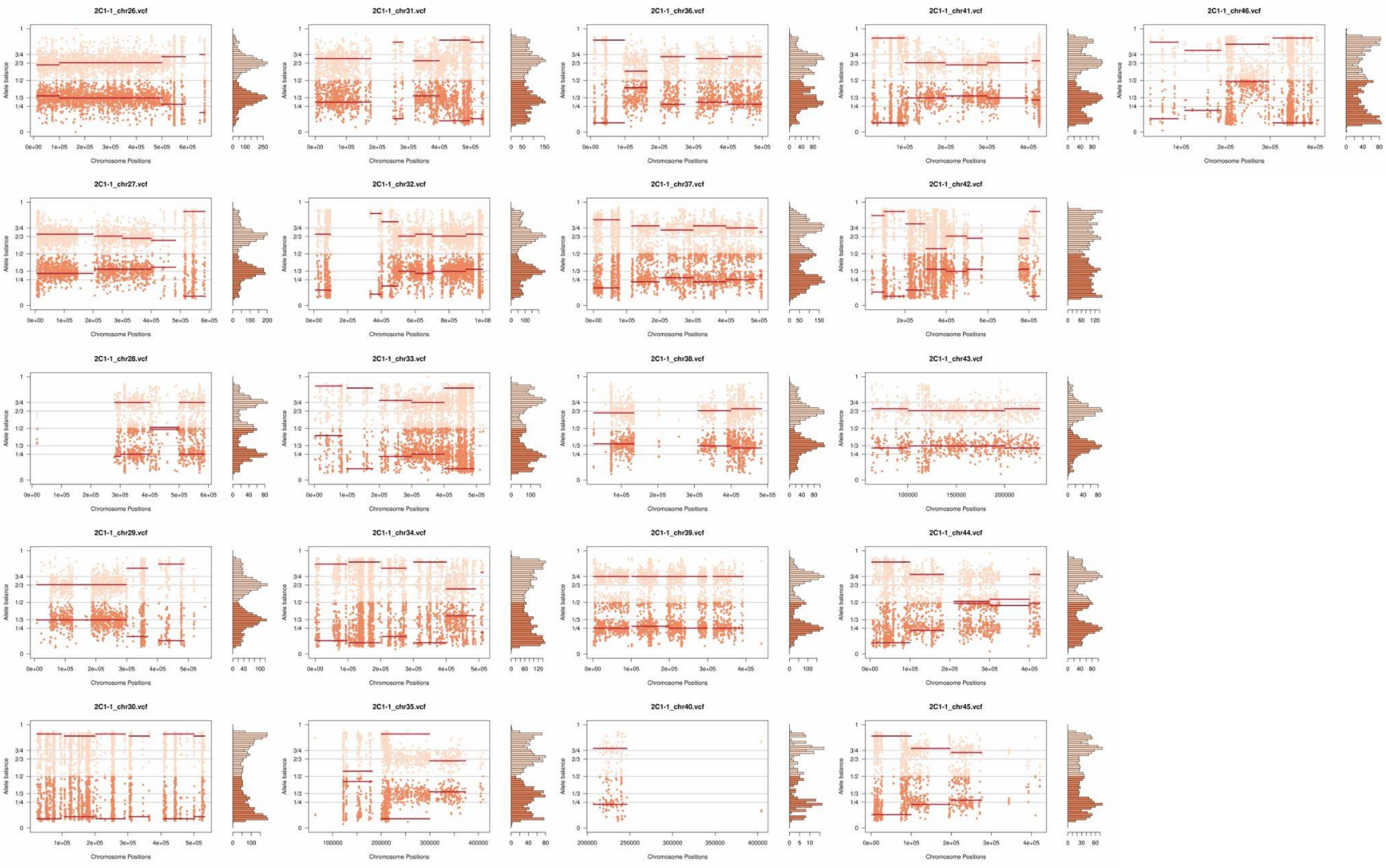
2C1-1_chr26.vcf
2C1-1_chr31.vcf
2C1-1_chr36.vcf
2C1-1_chr41.vcf
2C1-1_chr46.vcf
2C1-1_chr27.vcf
2C1-1_chr32.vcf
2C1-1_chr37.vcf
2C1-1_chr42.vcf
2C1-1_chr28.vcf
2C1-1_chr33.vcf
2C1-1_chr38.vcf
2C1-1_chr43.vcf
2C1-1_chr29.vcf
2C1-1_chr34.vcf
2C1-1_chr39.vcf
2C1-1_chr44.vcf
2C1-1_chr30.vcf
2C1-1_chr35.vcf
2C1-1_chr40.vcf
2C1-1_chr45.vcf
Allele balance
Chromosome Positions

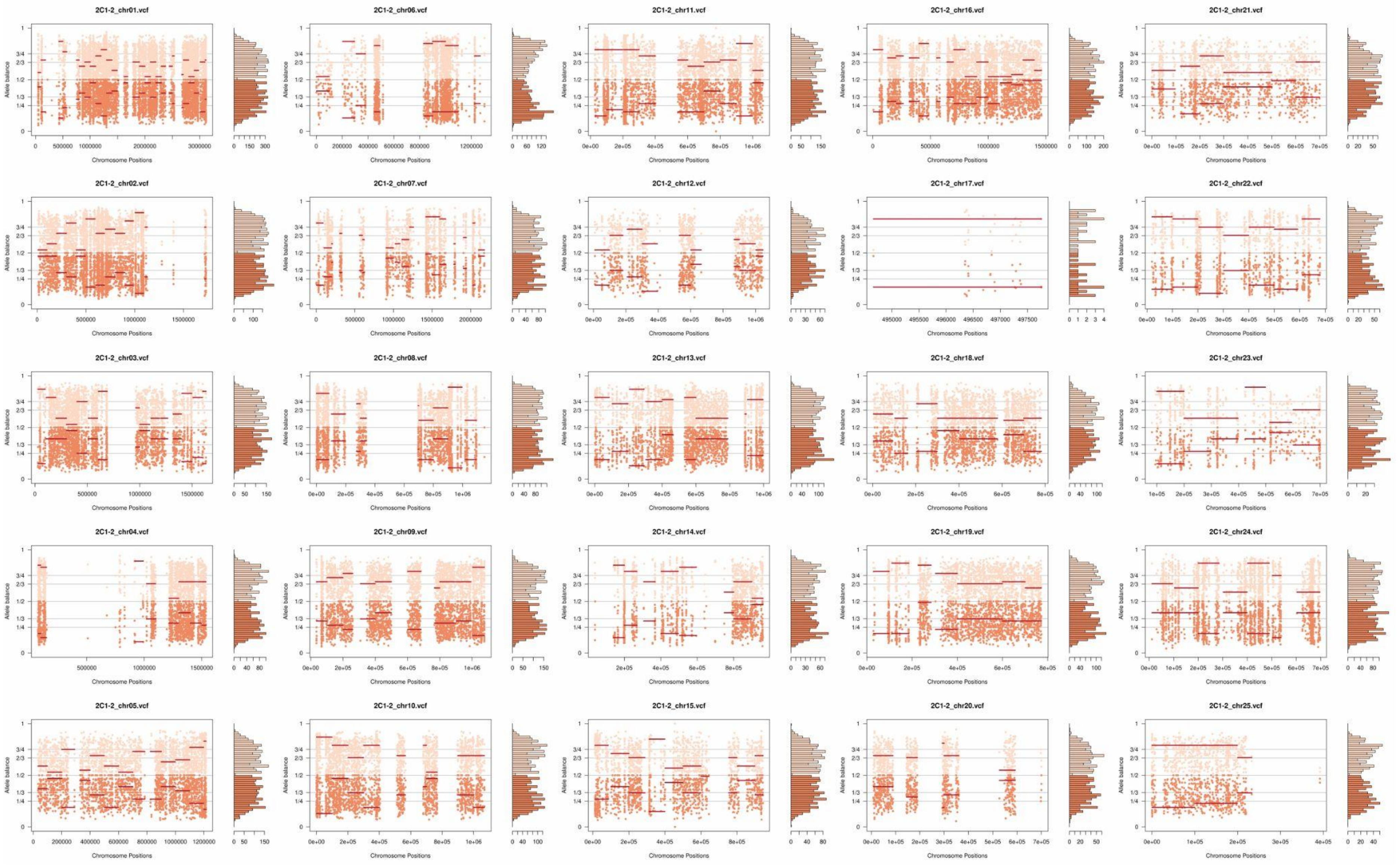
2C1-2_chr01.vcf
2C1-2_chr06.vcf
2C1-2_chr11.vcf
2C1-2_chr16.vcf
2C1-2_chr21.vcf
2C1-2_chr02.vcf
2C1-2_chr07.vcf
2C1-2_chr12.vcf
2C1-2_chr17.vcf
2C1-2_chr22.vcf
2C1-2_chr03.vcf
2C1-2_chr08.vcf
2C1-2_chr13.vcf
2C1-2_chr18.vcf
2C1-2_chr23.vcf
2C1-2_chr04.vcf
2C1-2_chr09.vcf
2C1-2_chr14.vcf
2C1-2_chr19.vcf
2C1-2_chr24.vcf
2C1-2_chr05.vcf
2C1-2_chr10.vcf
2C1-2_chr15.vcf
2C1-2_chr20.vcf
2C1-2_chr25.vcf
Allele balance
Chromosome Positions

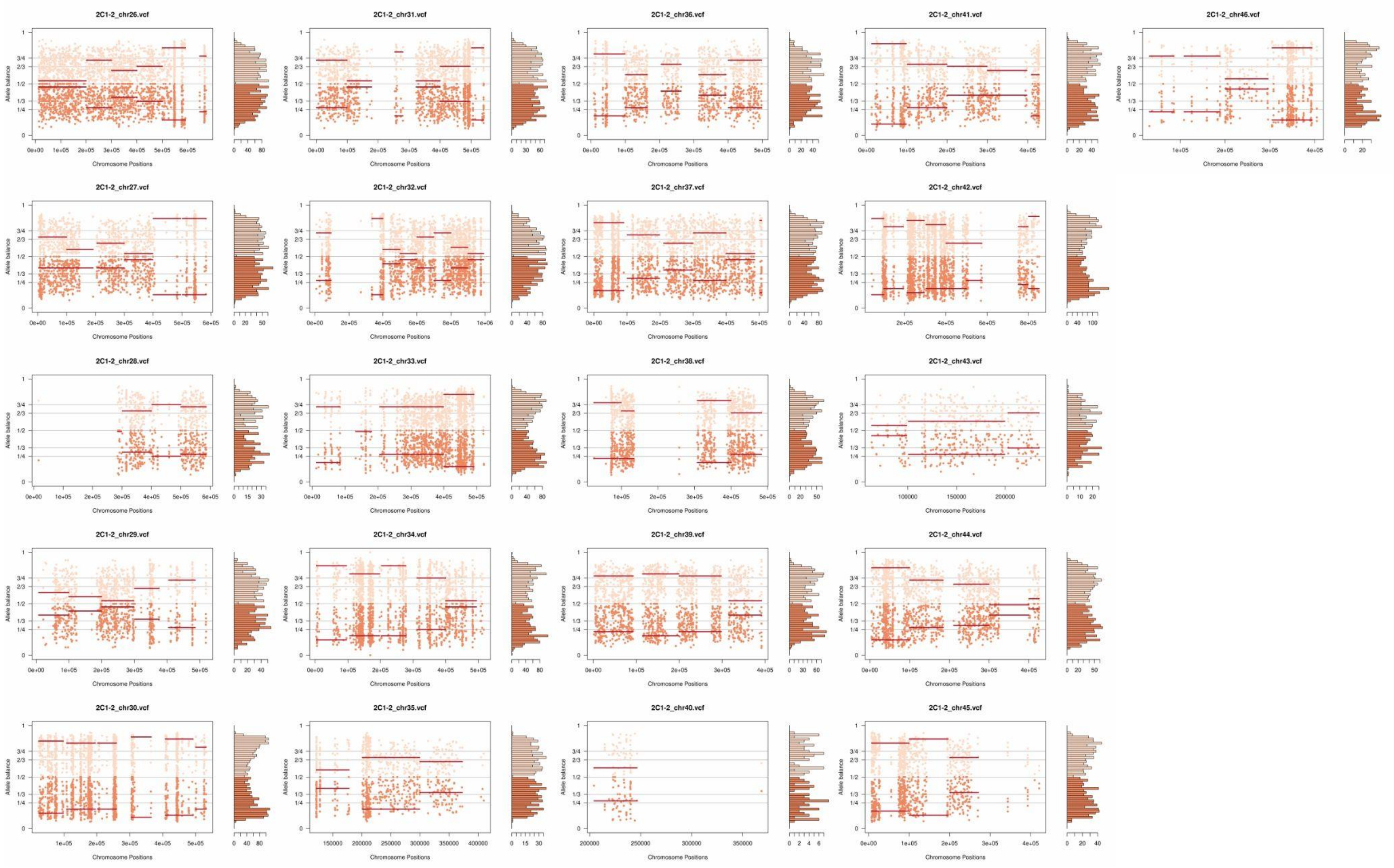
2C1-2_chr26.vcf
2C1-2_chr31.vcf
2C1-2_chr36.vcf
2C1-2_chr41.vcf
2C1-2_chr46.vcf
2C1-2_chr27.vcf
2C1-2_chr32.vcf
2C1-2_chr37.vcf
2C1-2_chr42.vcf
2C1-2_chr28.vcf
2C1-2_chr33.vcf
2C1-2_chr38.vcf
2C1-2_chr43.vcf
2C1-2_chr29.vcf
2C1-2_chr34.vcf
2C1-2_chr39.vcf
2C1-2_chr44.vcf
2C1-2_chr30.vcf
2C1-2_chr35.vcf
2C1-2_chr40.vcf
2C1-2_chr45.vcf
Allele balance
Chromosome Positions

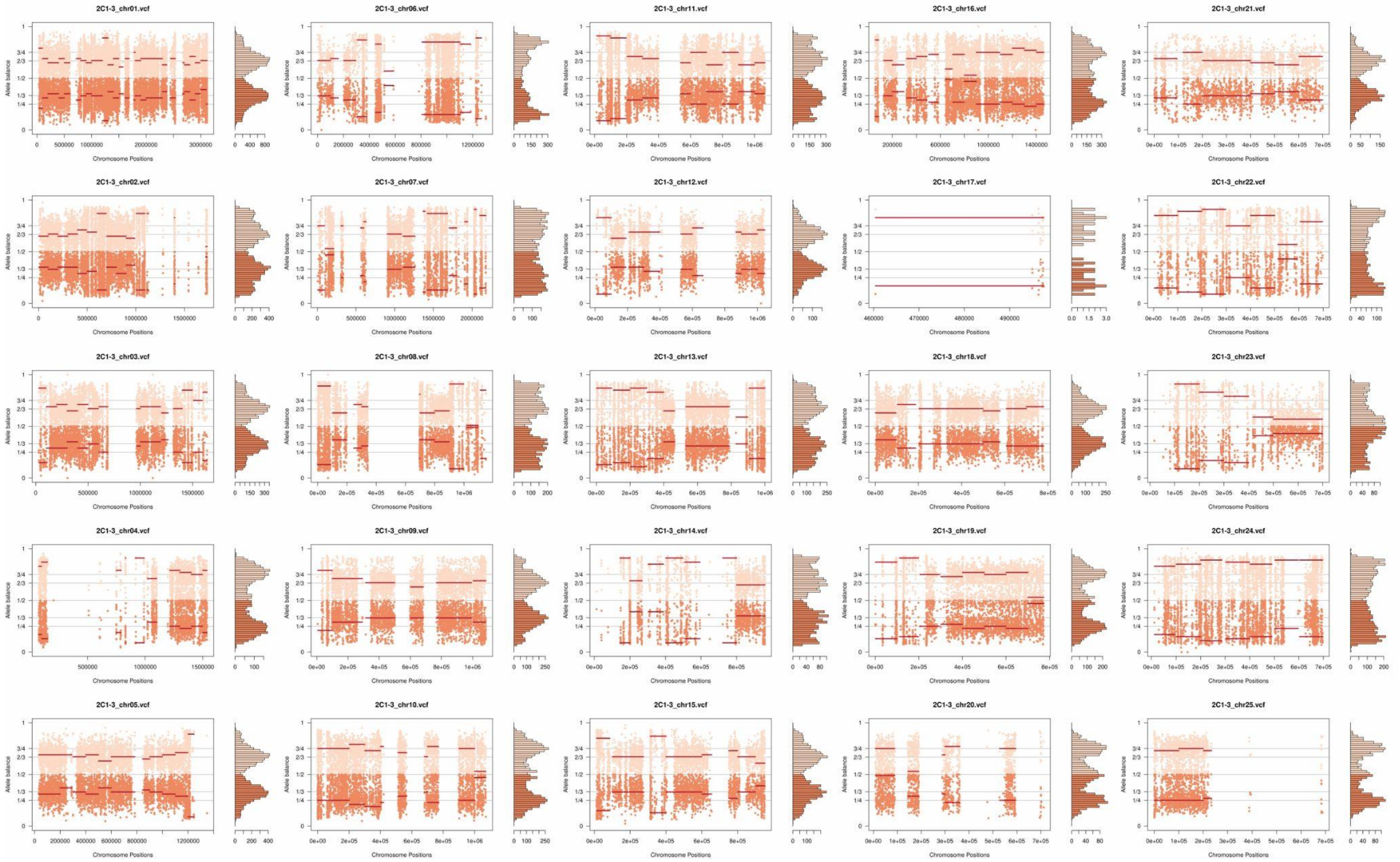
2C1-3_chr01.vcf
2C1-3_chr06.vcf
2C1-3_chr11.vcf
2C1-3_chr16.vcf
2C1-3_chr21.vcf
2C1-3_chr02.vcf
2C1-3_chr07.vcf
2C1-3_chr12.vcf
2C1-3_chr17.vcf
2C1-3_chr22.vcf
2C1-3_chr03.vcf
2C1-3_chr08.vcf
2C1-3_chr13.vcf
2C1-3_chr18.vcf
2C1-3_chr23.vcf
2C1-3_chr04.vcf
2C1-3_chr09.vcf
2C1-3_chr14.vcf
2C1-3_chr19.vcf
2C1-3_chr24.vcf
2C1-3_chr05.vcf
2C1-3_chr10.vcf
2C1-3_chr15.vcf
2C1-3_chr20.vcf
2C1-3_chr25.vcf
Allele balance
Chromosome Positions

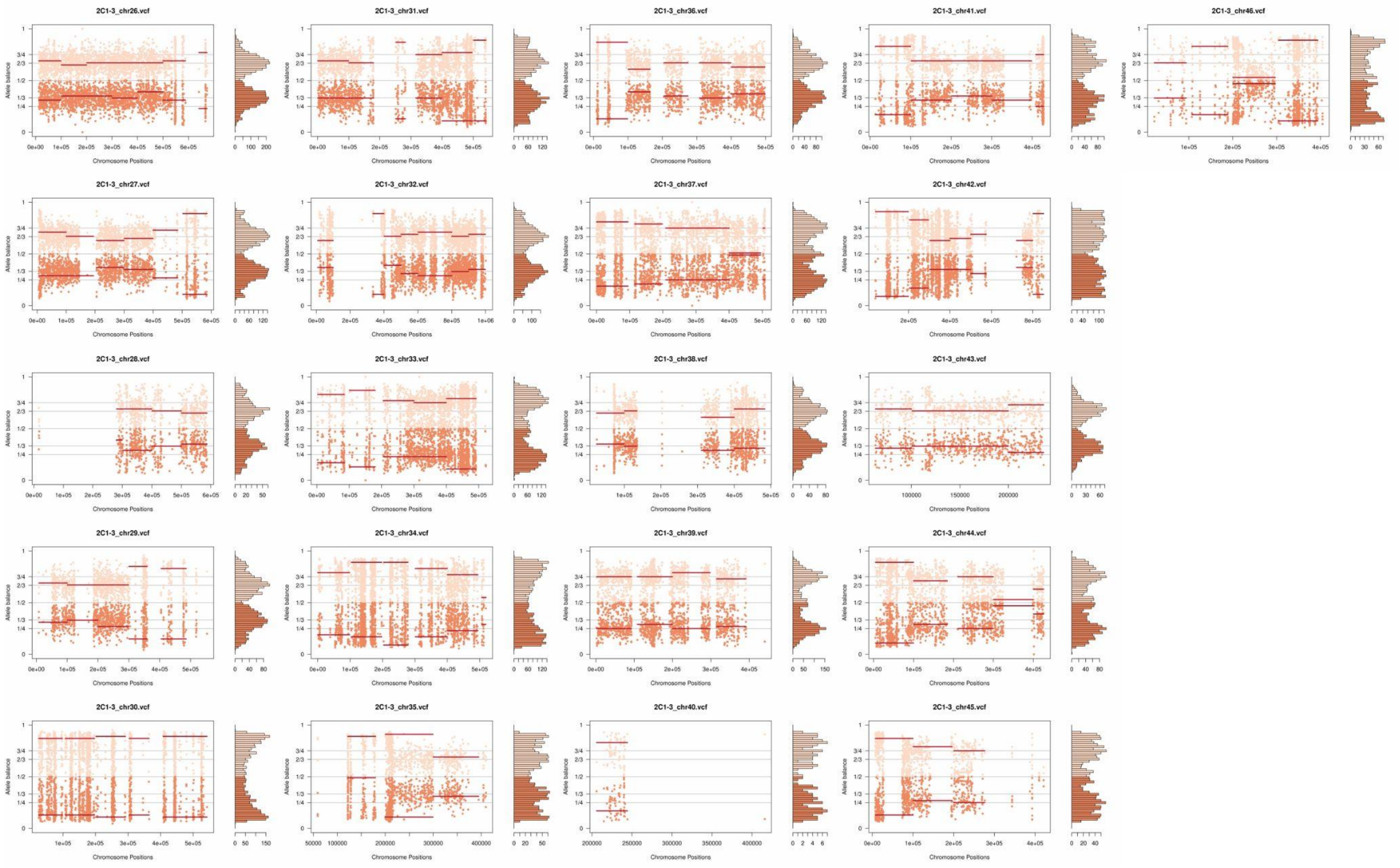
2C1-3_chr26.vcf
2C1-3_chr31.vcf
2C1-3_chr36.vcf
2C1-3_chr41.vcf
2C1-3_chr46.vcf
2C1-3_chr27.vcf
2C1-3_chr32.vcf
2C1-3_chr37.vcf
2C1-3_chr42.vcf
2C1-3_chr28.vcf
2C1-3_chr33.vcf
2C1-3_chr38.vcf
2C1-3_chr43.vcf
2C1-3_chr29.vcf
2C1-3_chr34.vcf
2C1-3_chr39.vcf
2C1-3_chr44.vcf
2C1-3_chr30.vcf
2C1-3_chr35.vcf
2C1-3_chr40.vcf
2C1-3_chr45.vcf
Allele balance
Chromosome Positions

**Supplementary Fig. 11:** Counts of genes displaying copy number variations after culture growth. **A.** Genes with copy losses. **B.** Genes with copy gains. In the x-axis is represented the counts of genes and the percentage of genes in that gene ontology category, or surface molecule family that showed CNV.

**A**

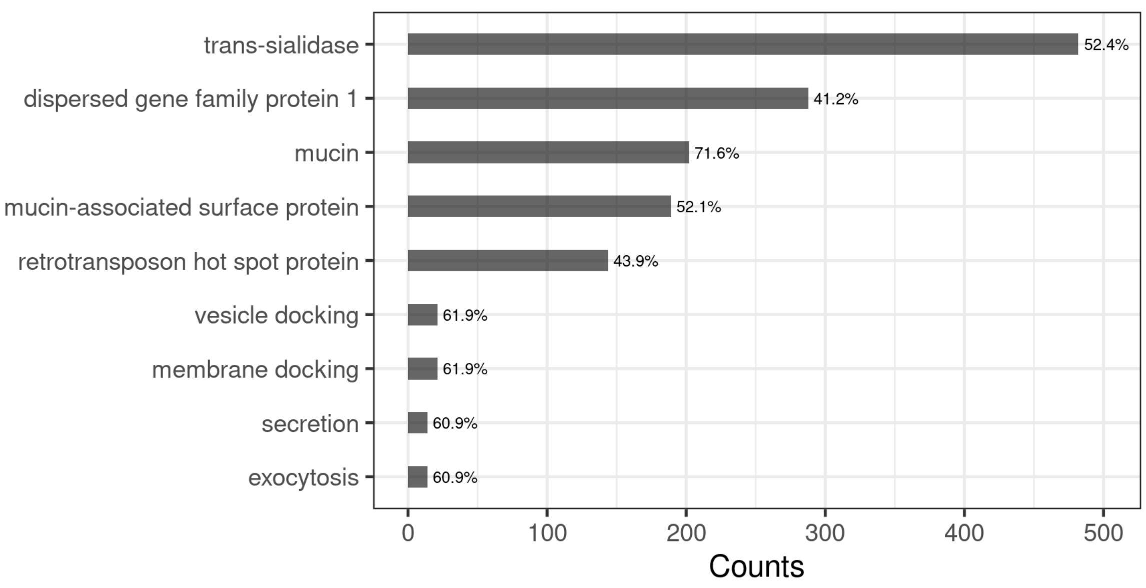

**B**

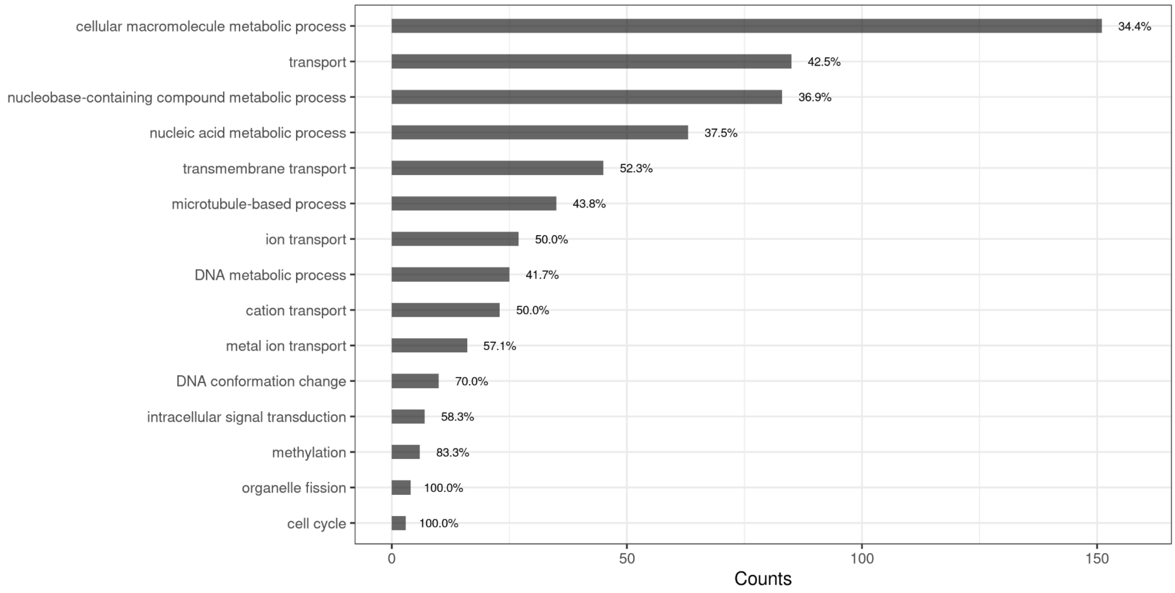

**Supplementary Table 1:** PCR-based multilocus microsatellite genotyping in parental and hybrid strains before (t=0) and after (t=800) the microevolution experiment.

| | | Microsatellite Loci and Genotypes | | | | | | | | | | |
|---|---|---|---|---|---|---|---|---|---|---|---|---|
| Parasite Line | Sample | 10101(TA) | | | 10101(TC) | | MCLF10 | | | 7093(TA)c | | |
| Parent 1 | Generation 0 | 160 | | 164 | 104 | 106 | | 183 | 185 | 182 | 186 | |
| | Generation 800 clone 1 | 160 | | 164 | 104 | 106 | | 183 | 185 | 182 | 186 | |
| | Generation 800 clone 2 | 160 | | 164 | 104 | 106 | | 183 | 185 | | *nd* | |
| Parent 2 | Generation 0 | | 162 | 164 | 104 | | 179 | | | | 186 | 188 |
| | Generation 800 clone 1 | | 162 | 164 | 104 | | 179 | | | | 186 | 188 |
| | Generation 800 clone 2 | | 162 | 164 | 104 | | 179 | | | | *nd* | |
| | Generation 800 clone 3 | | 162 | 164 | 104 | | 179 | | | | *nd* | |
| Hybrid "1D12" | Generation 0 | 160 | 162 | 164 | 104 | 106 | 179 | 183 | 185 | 182 | 186 | 188 |
| | Generation 800 clone 1 | 160 | 162 | 164 | 104 | 106 | 179 | 183 | 185 | 182 | 186 | 188 |
| | Generation 800 clone 2 | 160 | 162 | 164 | 104 | 106 | 179 | 183 | 185 | 182 | 186 | 188 |
| Hybrid "1C2" | Generation 0 | 160 | 162 | 164 | 104 | 106 | 179 | 183 | 185 | 182 | 186 | 188 |
| | Generation 800 clone 1 | 160 | 162 | 164 | 104 | 106 | 179 | 183 | 185 | 182 | 186 | 188 |
| | Generation 800 clone 2 | 160 | 162 | 164 | 104 | 106 | 179 | 183 | 185 | 182 | 186 | x |
| | Generation 800 clone 3 | 160 | 162 | 164 | 104 | 106 | 179 | 183 | 185 | 182 | 186 | 188 |
| Hybrid "2C1" | Generation 0 | 160 | 162 | 164 | 104 | 106 | 179 | 183 | 185 | 182 | 186 | x |
| | Generation 800 clone 1 | 160 | 162 | 164 | 104 | 106 | 179 | x | 185 | 182 | 186 | 188 |
| | Generation 800 clone 2 | 160 | 162 | 164 | 104 | 106 | 179 | 183 | 185 | 182 | 186 | 188 |
| | Generation 800 clone 3 | 160 | 162 | 164 | 104 | x | 179 | x | 185 | 182 | 186 | 188 |

**Supplementary Table 2**: Genome assembly statistics of parental strains.

| Feature | PI | PII |
|---|---|---|
| **Estimated genome size** | 46.1 Mb | 44.9 Mb |
| **Assembly Size** | 34.7 Mb | 32.9 Mb |
| **Number of scaffolds** | 544 | 585 |
| **Scaffolds > 100 Kb** | 114 | 96 |
| **Longest scaffold** | 886.4 Kb | 2.1 Mb |
| **Shortest scaffold** | 910 bp | 882 bp |
| **NG50** | 141.4 Kb | 183.5 Kb |
| **% of repeats** | 59.2 | 57.0 |
| **SNP rate** | 1/285 | 1/290 |

**Supplementary Table 3**: Effects of SNPs in the first generation hybrids.

| | SNPs inherited from the parental strains | | | |
|---|---|---|---|---|
| | Non-synonymous (NS) | Synonymous (S) | Nonsense | ratio NS/S |
| 1C2-0 | 16366 (57.96%) | 11557 (40.93%) | 312 (1.11%) | 1.41 |
| 1D12-0 | 15770 (57.71%) | 11242 (41.14%) | 312 (1.14%) | 1.40 |
| 2C1-0 | 16615 (58.07) | 11667 (40.78%) | 328 (1.15%) | 1.42 |
| | SNPs common to all hybrids | | | |
| | Non-synonymous (NS) | Synonymous (S) | Nonsense | ratio NS/S |
| 1C2-0 | 3796 (69.06%) | 1593 (28.98%) | 108 (1.96%) | 2.38 |
| 1D12-0 | 3459 (67.69%) | 1540 (30.14%) | 111 (2.17%) | 2.25 |
| 2C1-0 | 3666 (68.12%) | 1617 (30.04%) | 99 (1.84%) | 2.27 |
| | SNPs exclusive to each hybrid clone | | | |
| | Non-synonymous (NS) | Synonymous (S) | Nonsense | ratio NS/S |
| 1C2-0 | 1711 (69.22%) | 710 (28.72%) | 51 (2.06%) | 2.41 |
| 1D12-0 | 1553 (67.67%) | 696 (30.33%) | 46 (2.00%) | 2.23 |
| 2C1-0 | 1551 (69.09%) | 654 (29.13%) | 40 (1.78%) | 2.37 |

**Supplementary Table 4:** Number of SNPs in surface molecules genes (SM) and in other genes.

| | | SNPs in SM genes | SNPs in other genes | Ratio SM/other genes |
|---|---|---|---|---|
| | SNPs inherited from the parental strains | 4766 | 23390 | 0.20 |
| | SNPs common to all hybrids | 1398 | 1619 | 0.86 |
| 1C2-0 | SNPs exclusive to each hybrid clone | 1040 | 1423 | 0.73 |
| | | SNPs in SM genes | SNPs in other genes | Ratio SM/other genes |
| | SNPs inherited from the parental strains | 4683 | 22562 | 0.21 |
| | SNPs common to all hybrids | 1268 | 1538 | 0.82 |
| 1D12-0 | SNPs exclusive to each hybrid clone | 1002 | 1285 | 0.78 |
| | | SNPs in SM genes | SNPs in other genes | Ratio SM/other genes |
| | SNPs inherited from the parental strains | 4864 | 23660 | 0.21 |
| | SNPs common to all hybrids | 1431 | 1695 | 0.84 |
| 2C1-0 | SNPs exclusive to each hybrid clone | 973 | 1262 | 0.77 |

**Supplementary Table 5:** Number of non-synonymous SNPs in surface molecules genes (SM) and in other genes.

| | Non-synonymous SNPs inherited from parental strains | | | Non-synonymous SNPs after hybridization event | | |
|---|---|---|---|---|---|---|
| | SNPs in SM genes | SNPs in other genes | Ratio SM/other genes | SNPs in SM genes | SNPs in other genes | Ratio SM/other genes |
| 1C2-0 | 3067 | 13110 | 0.23 | 1646 | 2084 | 0.79 |
| 1D12-0 | 3029 | 12570 | 0.24 | 1516 | 1890 | 0.80 |
| 2C1-0 | 3161 | 13262 | 0.24 | 1607 | 2002 | 0.80 |

**Supplementary Table 6:** Gene Ontology analysis of expanded genes in parental strains after culture growth.

| ID | Name | Bgd count | Result count | Pct of bgd | Fold enrichment | Odds ratio | P-value | Benjamini | Bonferroni |
|---|---|---|---|---|---|---|---|---|---|
| GO:0030001 | metal ion transport | 28 | 13 | 46.4 | 5.14 | 9.11 | 2.19E-07 | 7.66E-05 | 7.66E-05 |
| GO:0055085 | transmembrane transport | 86 | 22 | 25.6 | 2.83 | 3.67 | 3.87E-06 | 6.77E-04 | 1.35E-03 |
| GO:0006812 | cation transport | 46 | 15 | 32.6 | 3.61 | 5.1 | 5.80E-06 | 6.77E-04 | 2.03E-03 |
| GO:0006811 | ion transport | 54 | 16 | 29.6 | 3.28 | 4.44 | 1.14E-05 | 9.97E-04 | 3.99E-03 |
| GO:0035556 | intracellular signal transduction | 12 | 7 | 58.3 | 6.46 | 14.44 | 2.42E-05 | 1.70E-03 | 8.48E-03 |
| GO:0009190 | cyclic nucleotide biosynthetic process | 9 | 6 | 66.7 | 7.38 | 20.57 | 3.42E-05 | 1.71E-03 | 1.20E-02 |
| GO:0009187 | cyclic nucleotide metabolic process | 9 | 6 | 66.7 | 7.38 | 20.57 | 3.42E-05 | 1.71E-03 | 1.20E-02 |
| GO:1901293 | nucleoside phosphate biosynthetic process | 37 | 12 | 32.4 | 3.59 | 05.01 | 5.46E-05 | 2.12E-03 | 1.91E-02 |
| GO:0009165 | nucleotide biosynthetic process | 37 | 12 | 32.4 | 3.59 | 05.01 | 5.46E-05 | 2.12E-03 | 1.91E-02 |
| GO:0090407 | organophosphate biosynthetic process | 44 | 13 | 29.5 | 3.27 | 4.38 | 8.01E-05 | 2.80E-03 | 2.80E-02 |
| GO:0009117 | nucleotide metabolic process | 43 | 12 | 27.9 | 3.9 | 04.03 | 2.75E-04 | 8.76E-03 | 9.64E-02 |
| GO:0019637 | organophosphate metabolic process | 56 | 14 | 25.0 | 2.77 | 3.48 | 3.07E-04 | 8.96E-03 | 1.08E-01 |
| GO:0006753 | nucleoside phosphate metabolic process | 44 | 12 | 27.3 | 03.02 | 3.9 | 3.49E-04 | 9.39E-03 | 1.22E-01 |
| GO:0006793 | phosphorus metabolic process | 175 | 29 | 16.6 | 1.84 | 2.12 | 7.31E-04 | 1.70E-02 | 2.56E-01 |
| GO:0006796 | phosphate-containing compound metabolic process | 175 | 29 | 16.6 | 1.84 | 2.12 | 7.31E-04 | 1.70E-02 | 2.56E-01 |
| GO:0007165 | signal transduction | 20 | 7 | 35.0 | 3.88 | 5.54 | 1.25E-03 | 2.74E-02 | 4.39E-01 |

| | | | | | | | | | |
|---|---|---|---|---|---|---|---|---|---|
| GO:0007154 | cell communication | 21 | 7 | 33.3 | 3.69 | 5.14 | 1.74E-03 | 3.38E-02 | 6.09E-01 |
| GO:0023052 | signaling | 21 | 7 | 33.3 | 3.69 | 5.14 | 1.74E-03 | 3.38E-02 | 6.09E-01 |
| GO:0055086 | nucleobase-containing small molecule metabolic process | 53 | 12 | 22.6 | 2.51 | 03.04 | 2.10E-03 | 3.87E-02 | 7.36E-01 |
| GO:0034654 | nucleobase-containing compound biosynthetic proc. | 72 | 14 | 19.4 | 2.15 | 2.51 | 4.20E-03 | 7.35E-02 | 1.00E+00 |
| GO:0051716 | cellular response to stimulus | 60 | 12 | 20.0 | 2.22 | 2.59 | 6.21E-03 | 1.04E-01 | 1.00E+00 |
| GO:0007018 | microtubule-based movement | 69 | 13 | 18.8 | 02.09 | 2.4 | 7.55E-03 | 1.14E-01 | 1.00E+00 |
| GO:0044248 | cellular catabolic process | 40 | 9 | 22.5 | 2.49 | 2.99 | 7.80E-03 | 1.14E-01 | 1.00E+00 |
| GO:0019438 | aromatic compound biosynthetic process | 77 | 14 | 18.2 | 02.01 | 2.3 | 7.82E-03 | 1.14E-01 | 1.00E+00 |
| GO:0006928 | movement of cell or subcellular component | 72 | 13 | 18.1 | 2.0 | 2.28 | 1.08E-02 | 1.52E-01 | 1.00E+00 |
| GO:0032259 | methylation | 6 | 3 | 50.0 | 5.54 | 10.17 | 1.18E-02 | 1.60E-01 | 1.00E+00 |
| GO:0018130 | heterocycle biosynthetic process | 82 | 14 | 17.1 | 1.89 | 2.13 | 1.36E-02 | 1.76E-01 | 1.00E+00 |
| GO:1901575 | organic substance catabolic process | 44 | 9 | 20.5 | 2.27 | 2.64 | 1.47E-02 | 1.77E-01 | 1.00E+00 |
| GO:0009056 | catabolic process | 44 | 9 | 20.5 | 2.27 | 2.64 | 1.47E-02 | 1.77E-01 | 1.00E+00 |
| GO:0019363 | pyridine nucleotide biosynthetic process | 12 | 4 | 33.3 | 3.69 | 5.1 | 1.80E-02 | 1.85E-01 | 1.00E+00 |
| GO:0019359 | nicotinamide nucleotide biosynthetic process | 12 | 4 | 33.3 | 3.69 | 5.1 | 1.80E-02 | 1.85E-01 | 1.00E+00 |
| GO:0072525 | pyridine-containing compound biosynthetic process | 12 | 4 | 33.3 | 3.69 | 5.1 | 1.80E-02 | 1.85E-01 | 1.00E+00 |
| GO:0043632 | modification-dependent macromolecule catabolic process | 18 | 5 | 27.8 | 3.8 | 3.93 | 1.86E-02 | 1.85E-01 | 1.00E+00 |

| | | | | | | | | | |
|---|---|---|---|---|---|---|---|---|---|
| GO:0006511 | ubiquitin-dependent protein catabolic process | 18 | 5 | 27.8 | 03.08 | 3.93 | 1.86E-02 | 1.85E-01 | 1.00E+00 |
| GO:0019941 | modification-dependent protein catabolic process | 18 | 5 | 27.8 | 03.08 | 3.93 | 1.86E-02 | 1.85E-01 | 1.00E+00 |
| GO:0006810 | transport | 200 | 27 | 13.5 | 1.5 | 1.63 | 1.95E-02 | 1.85E-01 | 1.00E+00 |
| GO:0051234 | establishment of localization | 200 | 27 | 13.5 | 1.5 | 1.63 | 1.95E-02 | 1.85E-01 | 1.00E+00 |
| GO:1901362 | organic cyclic compound biosynthetic process | 86 | 14 | 16.3 | 1.8 | 02.01 | 2.02E-02 | 1.86E-01 | 1.00E+00 |
| GO:0016310 | phosphorylation | 121 | 18 | 14.9 | 1.65 | 1.81 | 2.19E-02 | 1.95E-01 | 1.00E+00 |
| GO:0072524 | pyridine-containing compound metabolic process | 13 | 4 | 30.8 | 3.41 | 4.53 | 2.42E-02 | 1.95E-01 | 1.00E+00 |
| GO:0006091 | generation of precursor metabolites and energy | 13 | 4 | 30.8 | 3.41 | 4.53 | 2.42E-02 | 1.95E-01 | 1.00E+00 |
| GO:0046496 | nicotinamide nucleotide metabolic process | 13 | 4 | 30.8 | 3.41 | 4.53 | 2.42E-02 | 1.95E-01 | 1.00E+00 |
| GO:0019362 | pyridine nucleotide metabolic process | 13 | 4 | 30.8 | 3.41 | 4.53 | 2.42E-02 | 1.95E-01 | 1.00E+00 |
| GO:0051179 | localization | 204 | 27 | 13.2 | 1.47 | 1.59 | 2.47E-02 | 1.95E-01 | 1.00E+00 |
| GO:0007017 | microtubule-based process | 80 | 13 | 16.3 | 1.8 | 2.0 | 2.50E-02 | 1.95E-01 | 1.00E+00 |
| GO:0065007 | biological regulation | 81 | 13 | 16.0 | 1.78 | 1.97 | 2.75E-02 | 2.09E-01 | 1.00E+00 |
| GO:0050794 | regulation of cellular process | 73 | 12 | 16.4 | 1.82 | 02.03 | 2.83E-02 | 2.11E-01 | 1.00E+00 |
| GO:1901565 | organonitrogen compound catabolic process | 27 | 6 | 22.2 | 2.46 | 2.92 | 2.99E-02 | 2.18E-01 | 1.00E+00 |
| GO:0050789 | regulation of biological process | 74 | 12 | 16.2 | 1.8 | 1.99 | 3.12E-02 | 2.23E-01 | 1.00E+00 |
| GO:0072330 | monocarboxylic acid biosynthetic process | 15 | 4 | 26.7 | 2.95 | 3.7 | 4.00E-02 | 2.72E-01 | 1.00E+00 |
| GO:0006733 | oxidoreduction coenzyme metabolic process | 15 | 4 | 26.7 | 2.95 | 3.7 | 4.00E-02 | 2.72E-01 | 1.00E+00 |

**Supplementary Table 7**: Gene Ontology analysis of contracted genes in parental strains after culture growth.

| ID | Name | Bgd count | Result count | Pct of bgd | Fold enrichment | Odds ratio | P-value | Benjamini | Bonferroni |
|---|---|---|---|---|---|---|---|---|---|
| GO:0022610 | biological adhesion | 145 | 47 | 32.4 | 4.59 | 7.78 | 6.03E-21 | 8.09E-19 | 1.62E-18 |
| GO:0007155 | cell adhesion | 145 | 47 | 32.4 | 4.59 | 7.78 | 6.03E-21 | 8.09E-19 | 1.62E-18 |
| GO:0006508 | proteolysis | 295 | 51 | 17.3 | 2.45 | 3.29 | 2.34E-10 | 2.09E-08 | 6.26E-08 |
| GO:0019538 | protein metabolic process | 619 | 80 | 12.9 | 1.83 | 2.51 | 2.13E-09 | 1.43E-07 | 5.71E-07 |
| GO:1901564 | organonitrogen compound metabolic process | 705 | 83 | 11.8 | 1.67 | 2.22 | 9.82E-08 | 5.27E-06 | 2.63E-05 |
| GO:0043170 | macromolecule metabolic process | 754 | 86 | 11.4 | 1.62 | 2.15 | 2.31E-07 | 1.03E-05 | 6.18E-05 |
| GO:0042026 | protein refolding | 8 | 6 | 75.0 | 10.63 | 40.61 | 2.87E-06 | 1.10E-04 | 7.68E-04 |
| GO:0006807 | nitrogen compound metabolic process | 859 | 89 | 10.4 | 1.47 | 1.88 | 1.15E-05 | 3.85E-04 | 3.08E-03 |
| GO:0044238 | primary metabolic process | 919 | 91 | 9.9 | 1.4 | 1.77 | 6.11E-05 | 1.82E-03 | 1.64E-02 |
| GO:0071704 | organic substance metabolic process | 946 | 91 | 9.6 | 1.36 | 1.69 | 2.01E-04 | 5.37E-03 | 5.37E-02 |
| GO:0006414 | translational elongation | 27 | 8 | 29.6 | 4.2 | 5.72 | 3.71E-04 | 9.03E-03 | 9.94E-02 |
| GO:0006904 | vesicle docking involved in exocytosis | 20 | 6 | 30.0 | 4.25 | 5.78 | 1.93E-03 | 3.97E-02 | 5.16E-01 |
| GO:0140029 | exocytic process | 20 | 6 | 30.0 | 4.25 | 5.78 | 1.93E-03 | 3.97E-02 | 5.16E-01 |
| GO:0048278 | vesicle docking | 21 | 6 | 28.6 | 04.05 | 5.39 | 2.54E-03 | 4.01E-02 | 6.81E-01 |
| GO:0022406 | membrane docking | 21 | 6 | 28.6 | 04.05 | 5.39 | 2.54E-03 | 4.01E-02 | 6.81E-01 |
| GO:0140056 | organelle localization by membrane tethering | 21 | 6 | 28.6 | 04.05 | 5.39 | 2.54E-03 | 4.01E-02 | 6.81E-01 |
| GO:0051640 | organelle localization | 21 | 6 | 28.6 | 04.05 | 5.39 | 2.54E-03 | 4.01E-02 | 6.81E-01 |
| GO:0046903 | secretion | 23 | 6 | 26.1 | 3.7 | 4.75 | 4.20E-03 | 5.62E-02 | 1 |

| | | | | | | | | | |
|---|---|---|---|---|---|---|---|---|---|
| GO:0032940 | secretion by cell | 23 | 6 | 26.1 | 3.7 | 4.75 | 4.20E-03 | 5.62E-02 | 1 |
| GO:0006887 | exocytosis | 23 | 6 | 26.1 | 3.7 | 4.75 | 4.20E-03 | 5.62E-02 | 1 |
| GO:0051641 | cellular localization | 64 | 11 | 17.2 | 2.44 | 2.83 | 4.44E-03 | 5.66E-02 | 1 |
| GO:0008152 | metabolic process | 1336 | 112 | 8.4 | 1.19 | 1.42 | 7.43E-03 | 9.05E-02 | 1 |
| GO:0008150 | biological process | 1769 | 140 | 7.9 | 1.12 | 1.37 | 1.79E-02 | 2.08E-01 | 1 |
| GO:0006412 | translation | 167 | 19 | 11.4 | 1.61 | 1.76 | 2.37E-02 | 2.64E-01 | 1 |
| GO:0043043 | peptide biosynthetic process | 170 | 19 | 11.2 | 1.58 | 1.72 | 2.80E-02 | 3.00E-01 | 1 |
| GO:0043604 | amide biosynthetic process | 172 | 19 | 11.0 | 1.57 | 1.7 | 3.12E-02 | 3.21E-01 | 1 |
| GO:0006518 | peptide metabolic process | 173 | 19 | 11.0 | 1.56 | 1.69 | 3.29E-02 | 3.21E-01 | 1 |
| GO:0016192 | vesicle-mediated transport | 64 | 9 | 14.1 | 1.99 | 2.21 | 3.35E-02 | 3.21E-01 | 1 |
| GO:0043603 | cellular amide metabolic process | 175 | 19 | 10.9 | 1.54 | 1.66 | 3.64E-02 | 3.37E-01 | 1 |

**Supplementary Table 8:** Gene Ontology analysis of expanded genes in hybrid strains after culture growth.

| ID | Name | Bgd count | Result count | Pct of bgd | Fold enrichment | Odds ratio | P-value | Benjamini | Bonferroni |
|---|---|---|---|---|---|---|---|---|---|
| GO:0006508 | proteolysis | 295 | 172 | 58.3 | 1.89 | 3.62 | 5.61E-25 | 3.14E-22 | 3.14E-22 |
| GO:0043170 | macromolecule metabolic process | 754 | 328 | 43.5 | 1.41 | 2.12 | 9.59E-18 | 2.68E-15 | 5.36E-15 |
| GO:0006807 | nitrogen compound metabolic process | 859 | 361 | 42.0 | 1.37 | 02.02 | 8.90E-17 | 1.66E-14 | 4.98E-14 |
| GO:0019538 | protein metabolic process | 619 | 276 | 44.6 | 1.45 | 2.14 | 2.98E-16 | 4.16E-14 | 1.67E-13 |
| GO:1901564 | organonitrogen compound metabolic process | 705 | 299 | 42.4 | 1.38 | 1.96 | 5.33E-14 | 5.96E-12 | 2.98E-11 |
| GO:0071704 | organic substance metabolic process | 946 | 380 | 40.2 | 1.31 | 1.85 | 7.68E-14 | 7.16E-12 | 4.29E-11 |
| GO:0044238 | primary metabolic process | 919 | 368 | 40.0 | 1.3 | 1.82 | 4.97E-13 | 3.97E-11 | 2.78E-10 |
| GO:0008150 | biological process | 1769 | 604 | 34.1 | 1.11 | 1.46 | 1.48E-06 | 1.04E-04 | 8.30E-04 |
| GO:0055085 | transmembrane transport | 86 | 45 | 52.3 | 1.7 | 2.54 | 1.99E-05 | 1.23E-03 | 1.11E-02 |
| GO:0051179 | localization | 204 | 87 | 42.6 | 1.39 | 1.74 | 1.40E-04 | 7.81E-03 | 7.81E-02 |
| GO:0051234 | establishment of localization | 200 | 85 | 42.5 | 1.38 | 1.73 | 1.93E-04 | 9.00E-03 | 1.08E-01 |
| GO:0006810 | transport | 200 | 85 | 42.5 | 1.38 | 1.73 | 1.93E-04 | 9.00E-03 | 1.08E-01 |
| GO:0007155 | cell adhesion | 145 | 63 | 43.4 | 1.41 | 1.78 | 6.59E-04 | 2.63E-02 | 3.69E-01 |
| GO:0022610 | biological adhesion | 145 | 63 | 43.4 | 1.41 | 1.78 | 6.59E-04 | 2.63E-02 | 3.69E-01 |
| GO:0008152 | metabolic process | 1336 | 450 | 33.7 | 01.09 | 1.27 | 1.30E-03 | 4.86E-02 | 7.29E-01 |
| GO:0006811 | ion transport | 54 | 27 | 50.0 | 1.62 | 2.29 | 2.25E-03 | 7.84E-02 | 1.00E+00 |

| | | | | | | | | | |
|---|---|---|---|---|---|---|---|---|---|
| GO:0030001 | metal ion transport | 28 | 16 | 57.1 | 1.86 | 03.03 | 3.27E-03 | 1.07E-01 | 1.00E+00 |
| GO:0006812 | cation transport | 46 | 23 | 50.0 | 1.62 | 2.28 | 4.68E-03 | 1.39E-01 | 1.00E+00 |
| GO:0009187 | cyclic nucleotide metabolic process | 9 | 7 | 77.8 | 2.53 | 7.92 | 4.98E-03 | 1.39E-01 | 1.00E+00 |
| GO:0009190 | cyclic nucleotide biosynthetic process | 9 | 7 | 77.8 | 2.53 | 7.92 | 4.98E-03 | 1.39E-01 | 1.00E+00 |
| GO:0042493 | response to drug | 4 | 4 | 100.0 | 3.25 | inf | 8.93E-03 | 1.92E-01 | 1.00E+00 |
| GO:0006855 | drug transmembrane transport | 4 | 4 | 100.0 | 3.25 | inf | 8.93E-03 | 1.92E-01 | 1.00E+00 |
| GO:0042221 | response to chemical | 4 | 4 | 100.0 | 3.25 | inf | 8.93E-03 | 1.92E-01 | 1.00E+00 |
| GO:0015893 | drug transport | 4 | 4 | 100.0 | 3.25 | inf | 8.93E-03 | 1.92E-01 | 1.00E+00 |
| GO:0048285 | organelle fission | 4 | 4 | 100.0 | 3.25 | inf | 8.93E-03 | 1.92E-01 | 1.00E+00 |
| GO:0007017 | microtubule-based process | 80 | 35 | 43.8 | 1.42 | 1.78 | 8.94E-03 | 1.92E-01 | 1.00E+00 |
| GO:0006928 | movement of cell or subcellular component | 72 | 32 | 44.4 | 1.44 | 1.83 | 9.26E-03 | 1.92E-01 | 1.00E+00 |
| GO:0046483 | heterocycle metabolic process | 234 | 88 | 37.6 | 1.22 | 1.39 | 1.22E-02 | 2.28E-01 | 1.00E+00 |
| GO:0071103 | DNA conformation change | 10 | 7 | 70.0 | 2.27 | 5.28 | 1.22E-02 | 2.28E-01 | 1.00E+00 |
| GO:0032259 | methylation | 6 | 5 | 83.3 | 2.71 | 11.3 | 1.22E-02 | 2.28E-01 | 1.00E+00 |
| GO:0007018 | microtubule-based movement | 69 | 30 | 43.5 | 1.41 | 1.75 | 1.65E-02 | 2.97E-01 | 1.00E+00 |
| GO:0034641 | cellular nitrogen compound metabolic process | 378 | 134 | 35.4 | 1.15 | 1.27 | 2.14E-02 | 3.41E-01 | 1.00E+00 |
| GO:0006139 | nucleobase-containing compound metabolic process | 225 | 83 | 36.9 | 1.2 | 1.34 | 2.46E-02 | 3.41E-01 | 1.00E+00 |
| GO:0008295 | spermidine biosynthetic process | 3 | 3 | 100.0 | 3.25 | inf | 2.91E-02 | 3.41E-01 | 1.00E+00 |

| | | | | | | | | | |
|---|---|---|---|---|---|---|---|---|---|
| GO:0009308 | amine metabolic process | 3 | 3 | 100.0 | 3.25 | inf | 2.91E-02 | 3.41E-01 | 1.00E+00 |
| GO:0044106 | cellular amine metabolic process | 3 | 3 | 100.0 | 3.25 | inf | 2.91E-02 | 3.41E-01 | 1.00E+00 |
| GO:0006750 | glutathione biosynthetic process | 3 | 3 | 100.0 | 3.25 | inf | 2.91E-02 | 3.41E-01 | 1.00E+00 |
| GO:0006595 | polyamine metabolic process | 3 | 3 | 100.0 | 3.25 | inf | 2.91E-02 | 3.41E-01 | 1.00E+00 |
| GO:0019184 | nonribosomal peptide biosynthetic process | 3 | 3 | 100.0 | 3.25 | inf | 2.91E-02 | 3.41E-01 | 1.00E+00 |
| GO:0008216 | spermidine metabolic process | 3 | 3 | 100.0 | 3.25 | inf | 2.91E-02 | 3.41E-01 | 1.00E+00 |
| GO:0009309 | amine biosynthetic process | 3 | 3 | 100.0 | 3.25 | inf | 2.91E-02 | 3.41E-01 | 1.00E+00 |
| GO:0042401 | cellular biogenic amine biosynthetic process | 3 | 3 | 100.0 | 3.25 | inf | 2.91E-02 | 3.41E-01 | 1.00E+00 |
| GO:0097164 | ammonium ion metabolic process | 3 | 3 | 100.0 | 3.25 | inf | 2.91E-02 | 3.41E-01 | 1.00E+00 |
| GO:0044262 | cellular carbohydrate metabolic process | 3 | 3 | 100.0 | 3.25 | inf | 2.91E-02 | 3.41E-01 | 1.00E+00 |
| GO:0007049 | cell cycle | 3 | 3 | 100.0 | 3.25 | inf | 2.91E-02 | 3.41E-01 | 1.00E+00 |
| GO:0006596 | polyamine biosynthetic process | 3 | 3 | 100.0 | 3.25 | inf | 2.91E-02 | 3.41E-01 | 1.00E+00 |
| GO:0006576 | cellular biogenic amine metabolic process | 3 | 3 | 100.0 | 3.25 | inf | 2.91E-02 | 3.41E-01 | 1.00E+00 |
| GO:0051186 | cofactor metabolic process | 31 | 15 | 48.4 | 1.57 | 2.13 | 2.93E-02 | 3.41E-01 | 1.00E+00 |
| GO:0090304 | nucleic acid metabolic process | 168 | 63 | 37.5 | 1.22 | 1.37 | 3.33E-02 | 3.80E-01 | 1.00E+00 |
| GO:0044260 | cellular macromolecule metabolic process | 439 | 151 | 34.4 | 1.12 | 1.21 | 4.37E-02 | 4.56E-01 | 1.00E+00 |
| GO:0035556 | intracellular signal transduction | 12 | 7 | 58.3 | 1.9 | 3.16 | 4.38E-02 | 4.56E-01 | 1.00E+00 |
| GO:0006259 | DNA metabolic process | 60 | 25 | 41.7 | 1.35 | 1.62 | 4.67E-02 | 4.56E-01 | 1.00E+00 |

| | | | | | | | | | |
|---|---|---|---|---|---|---|---|---|---|
| GO:0051716 | cellular response to stimulus | 60 | 25 | 41.7 | 1.35 | 1.62 | 4.67E-02 | 4.56E-01 | 1.00E+00 |

**Supplementary Table 9**: Gene Ontology analysis of contracted genes in hybrid strains after culture growth.

| ID | Name | Bgd count | Result count | Pct of bgd | Fold enrichment | Odds ratio | P-value | Benjamini | Bonferroni |
|---|---|---|---|---|---|---|---|---|---|
| GO:0006508 | proteolysis | 295 | 144 | 48.8 | 1.81 | 2.91 | 2.48E-17 | 8.97E-15 | 8.97E-15 |
| GO:0019538 | protein metabolic process | 619 | 210 | 33.9 | 1.26 | 1.52 | 1.08E-05 | 1.95E-03 | 3.90E-03 |
| GO:1901564 | organonitrogen compound metabolic process | 705 | 234 | 33.2 | 1.23 | 1.48 | 1.74E-05 | 2.10E-03 | 6.29E-03 |
| GO:0140029 | exocytic process | 20 | 13 | 65.0 | 2.41 | 5.1 | 3.97E-04 | 2.37E-02 | 1.43E-01 |
| GO:0006904 | vesicle docking involved in exocytosis | 20 | 13 | 65.0 | 2.41 | 5.1 | 3.97E-04 | 2.37E-02 | 1.43E-01 |
| GO:0006887 | exocytosis | 23 | 14 | 60.9 | 2.26 | 4.27 | 6.27E-04 | 2.37E-02 | 2.26E-01 |
| GO:0032940 | secretion by cell | 23 | 14 | 60.9 | 2.26 | 4.27 | 6.27E-04 | 2.37E-02 | 2.26E-01 |
| GO:0046903 | secretion | 23 | 14 | 60.9 | 2.26 | 4.27 | 6.27E-04 | 2.37E-02 | 2.26E-01 |
| GO:0140056 | organelle localization by membrane tethering | 21 | 13 | 61.9 | 2.3 | 4.46 | 7.87E-04 | 2.37E-02 | 2.84E-01 |
| GO:0051640 | organelle localization | 21 | 13 | 61.9 | 2.3 | 4.46 | 7.87E-04 | 2.37E-02 | 2.84E-01 |
| GO:0048278 | vesicle docking | 21 | 13 | 61.9 | 2.3 | 4.46 | 7.87E-04 | 2.37E-02 | 2.84E-01 |
| GO:0022406 | membrane docking | 21 | 13 | 61.9 | 2.3 | 4.46 | 7.87E-04 | 2.37E-02 | 2.84E-01 |
| GO:0009072 | aromatic amino acid family metabolic process | 12 | 8 | 66.7 | 2.47 | 5.46 | 4.58E-03 | 1.27E-01 | 1.00E+00 |
| GO:0006520 | cellular amino acid metabolic process | 67 | 28 | 41.8 | 1.55 | 1.98 | 5.61E-03 | 1.45E-01 | 1.00E+00 |
| GO:0042026 | protein refolding | 8 | 6 | 75.0 | 2.78 | 8.18 | 6.30E-03 | 1.52E-01 | 1.00E+00 |
| GO:0043170 | macromolecule metabolic process | 754 | 227 | 30.1 | 1.12 | 1.23 | 1.46E-02 | 3.30E-01 | 1.00E+00 |
| GO:0006807 | nitrogen compound metabolic process | 859 | 252 | 29.3 | 01.09 | 1.18 | 3.62E-02 | 6.90E-01 | 1.00E+00 |
| GO:0044238 | primary metabolic process | 919 | 268 | 29.2 | 01.08 | 1.17 | 4.03E-02 | 6.90E-01 | 1.00E+00 |

**Supplementary Table 10**: Gene Ontology analysis of genes encoded in chromosome 19.

| ID | Name | Bgd count | Result count | Pct of bgd | Fold enrichment | Odds ratio | P-value | Benjamini | Bonferroni |
|---|---|---|---|---|---|---|---|---|---|
| GO:0019400 | alditol metabolic process | 2 | 2 | 100.0 | 41.17 | inf | 5.82E-04 | 3.93E-02 | 8.79E-02 |
| GO:0006071 | glycerol metabolic process | 2 | 2 | 100.0 | 41.17 | inf | 5.82E-04 | 3.93E-02 | 8.79E-02 |
| GO:0010467 | gene expression | 240 | 14 | 5.8 | 2.4 | 2.83 | 1.53E-03 | 3.93E-02 | 2.30E-01 |
| GO:0044262 | cellular carbohydrate metabolic process | 3 | 2 | 66.7 | 27.45 | 82.52 | 1.72E-03 | 3.93E-02 | 2.60E-01 |
| GO:0006412 | translation | 167 | 11 | 6.6 | 2.71 | 3.15 | 1.98E-03 | 3.93E-02 | 2.99E-01 |
| GO:0043043 | peptide biosynthetic process | 170 | 11 | 6.5 | 2.66 | 03.09 | 2.29E-03 | 3.93E-02 | 3.45E-01 |
| GO:0043604 | amide biosynthetic process | 170 | 11 | 6.5 | 2.66 | 03.09 | 2.29E-03 | 3.93E-02 | 3.45E-01 |
| GO:0006518 | peptide metabolic process | 173 | 11 | 6.4 | 2.62 | 03.02 | 2.63E-03 | 3.93E-02 | 3.97E-01 |
| GO:0043603 | cellular amide metabolic process | 173 | 11 | 6.4 | 2.62 | 03.02 | 2.63E-03 | 3.93E-02 | 3.97E-01 |
| GO:0042221 | response to chemical | 4 | 2 | 50.0 | 20.59 | 41.25 | 3.38E-03 | 3.93E-02 | 5.11E-01 |
| GO:0015893 | drug transport | 4 | 2 | 50.0 | 20.59 | 41.25 | 3.38E-03 | 3.93E-02 | 5.11E-01 |
| GO:0042493 | response to drug | 4 | 2 | 50.0 | 20.59 | 41.25 | 3.38E-03 | 3.93E-02 | 5.11E-01 |
| GO:0006855 | drug transmembrane transport | 4 | 2 | 50.0 | 20.59 | 41.25 | 3.38E-03 | 3.93E-02 | 5.11E-01 |
| GO:1901566 | organonitrogen compound biosynthetic process | 211 | 12 | 5.7 | 2.34 | 2.69 | 4.24E-03 | 4.57E-02 | 6.40E-01 |
| GO:0044271 | cellular nitrogen compound biosynthetic process | 241 | 13 | 5.4 | 2.22 | 2.56 | 4.57E-03 | 4.60E-02 | 6.90E-01 |
| GO:0034641 | cellular nitrogen compound metabolic process | 362 | 17 | 4.7 | 1.93 | 2.27 | 4.88E-03 | 4.61E-02 | 7.37E-01 |

| | | | | | | | | | |
|---|---|---|---|---|---|---|---|---|---|
| GO:0019751 | polyol metabolic process | 5 | 2 | 40.0 | 16.47 | 27.49 | 5.55E-03 | 4.93E-02 | 8.38E-01 |
| GO:0009059 | macromolecule biosynthetic process | 226 | 12 | 5.3 | 2.19 | 2.49 | 7.35E-03 | 5.84E-02 | 1.00E+00 |
| GO:0034645 | cellular macromolecule biosynthetic process | 226 | 12 | 5.3 | 2.19 | 2.49 | 7.35E-03 | 5.84E-02 | 1.00E+00 |
| GO:0044267 | cellular protein metabolic process | 350 | 16 | 4.6 | 1.88 | 2.18 | 8.31E-03 | 6.28E-02 | 1.00E+00 |
| GO:0006066 | alcohol metabolic process | 7 | 2 | 28.6 | 11.76 | 16.48 | 1.13E-02 | 8.08E-02 | 1.00E+00 |
| GO:0044237 | cellular metabolic process | 632 | 24 | 3.8 | 1.56 | 1.86 | 1.18E-02 | 8.08E-02 | 1.00E+00 |
| GO:1901615 | organic hydroxy compound metabolic process | 8 | 2 | 25.0 | 10.29 | 13.73 | 1.48E-02 | 8.15E-02 | 1.00E+00 |
| GO:0071840 | cellular component organization or biogenesis | 62 | 5 | 8.1 | 3.32 | 3.7 | 1.63E-02 | 8.15E-02 | 1.00E+00 |
| GO:0044249 | cellular biosynthetic process | 288 | 13 | 4.5 | 1.86 | 02.09 | 1.94E-02 | 8.15E-02 | 1.00E+00 |
| GO:0042727 | flavin-containing compound biosynthetic process | 1 | 1 | 100.0 | 41.17 | inf | 2.43E-02 | 8.15E-02 | 1.00E+00 |
| GO:0042726 | flavin-containing compound metabolic process | 1 | 1 | 100.0 | 41.17 | inf | 2.43E-02 | 8.15E-02 | 1.00E+00 |
| GO:0042364 | water-soluble vitamin biosynthetic process | 1 | 1 | 100.0 | 41.17 | inf | 2.43E-02 | 8.15E-02 | 1.00E+00 |
| GO:0042274 | ribosomal small subunit biogenesis | 1 | 1 | 100.0 | 41.17 | inf | 2.43E-02 | 8.15E-02 | 1.00E+00 |
| GO:0042127 | regulation of cell population proliferation | 1 | 1 | 100.0 | 41.17 | inf | 2.43E-02 | 8.15E-02 | 1.00E+00 |
| GO:0000462 | maturation of SSU-rRNA from tricistronic rRNA transcript | 1 | 1 | 100.0 | 41.17 | inf | 2.43E-02 | 8.15E-02 | 1.00E+00 |
| GO:0006740 | NADPH regeneration | 1 | 1 | 100.0 | 41.17 | inf | 2.43E-02 | 8.15E-02 | 1.00E+00 |
| GO:0009231 | riboflavin biosynthetic process | 1 | 1 | 100.0 | 41.17 | inf | 2.43E-02 | 8.15E-02 | 1.00E+00 |
| GO:0009110 | vitamin biosynthetic process | 1 | 1 | 100.0 | 41.17 | inf | 2.43E-02 | 8.15E-02 | 1.00E+00 |

| | | | | | | | | | |
|---|---|---|---|---|---|---|---|---|---|
| GO:0008284 | positive regulation of cell population proliferation | 1 | 1 | 100.0 | 41.17 | inf | 2.43E-02 | 8.15E-02 | 1.00E+00 |
| GO:0006098 | pentose-phosphate shunt | 1 | 1 | 100.0 | 41.17 | inf | 2.43E-02 | 8.15E-02 | 1.00E+00 |
| GO:0006952 | defense response | 1 | 1 | 100.0 | 41.17 | inf | 2.43E-02 | 8.15E-02 | 1.00E+00 |
| GO:0006771 | riboflavin metabolic process | 1 | 1 | 100.0 | 41.17 | inf | 2.43E-02 | 8.15E-02 | 1.00E+00 |
| GO:0006767 | water-soluble vitamin metabolic process | 1 | 1 | 100.0 | 41.17 | inf | 2.43E-02 | 8.15E-02 | 1.00E+00 |
| GO:0006766 | vitamin metabolic process | 1 | 1 | 100.0 | 41.17 | inf | 2.43E-02 | 8.15E-02 | 1.00E+00 |
| GO:0030490 | maturation of SSU-rRNA | 1 | 1 | 100.0 | 41.17 | inf | 2.43E-02 | 8.15E-02 | 1.00E+00 |
| GO:0006739 | NADP metabolic process | 1 | 1 | 100.0 | 41.17 | inf | 2.43E-02 | 8.15E-02 | 1.00E+00 |
| GO:0006354 | DNA-templated transcription, elongation | 1 | 1 | 100.0 | 41.17 | inf | 2.43E-02 | 8.15E-02 | 1.00E+00 |
| GO:0006368 | transcription elongation from RNA pol II promoter | 1 | 1 | 100.0 | 41.17 | inf | 2.43E-02 | 8.15E-02 | 1.00E+00 |
| GO:0006366 | transcription by RNA polymerase II | 1 | 1 | 100.0 | 41.17 | inf | 2.43E-02 | 8.15E-02 | 1.00E+00 |
| GO:0016043 | cellular component organization | 47 | 4 | 8.5 | 3.5 | 3.89 | 2.60E-02 | 8.54E-02 | 1.00E+00 |
| GO:0044260 | cellular macromolecule metabolic process | 439 | 17 | 3.9 | 1.59 | 1.8 | 3.07E-02 | 9.86E-02 | 1.00E+00 |
| GO:1901576 | organic substance biosynthetic process | 315 | 13 | 4.1 | 1.7 | 1.88 | 3.75E-02 | 1.18E-01 | 1.00E+00 |
| GO:0006370 | 7-methylguanosine mRNA capping | 2 | 1 | 50.0 | 20.59 | 40.7 | 4.80E-02 | 1.39E-01 | 1.00E+00 |
| GO:0007031 | peroxisome organization | 2 | 1 | 50.0 | 20.59 | 40.7 | 4.80E-02 | 1.39E-01 | 1.00E+00 |
| GO:0051156 | glucose 6-phosphate metabolic process | 2 | 1 | 50.0 | 20.59 | 40.7 | 4.80E-02 | 1.39E-01 | 1.00E+00 |
| GO:0016559 | peroxisome fission | 2 | 1 | 50.0 | 20.59 | 40.7 | 4.80E-02 | 1.39E-01 | 1.00E+00 |

**Supplementary Table 11**: Gene Ontology analysis of genes encoded in chromosome 37.

| ID | Name | Bgd count | Result count | Pct of bgd | Fold enrichment | Odds ratio | P-value | Benjamini | Bonferroni |
|---|---|---|---|---|---|---|---|---|---|
| GO:0032259 | methylation | 6 | 2 | 33.3 | 22.87 | 35.34 | 3.00E-03 | 1.72E-01 | 2.49E-01 |
| GO:0006298 | mismatch repair | 10 | 2 | 20.0 | 13.72 | 17.65 | 8.68E-03 | 1.72E-01 | 7.20E-01 |
| GO:0006281 | DNA repair | 32 | 3 | 9.4 | 6.43 | 7.42 | 1.07E-02 | 1.72E-01 | 8.86E-01 |
| GO:0000375 | RNA splicing, via transesterification reactions | 1 | 1 | 100.0 | 68.62 | inf | 1.46E-02 | 1.72E-01 | 1.00E+00 |
| GO:0000377 | RNA splicing, via transesterification reactions With bulged adenosine as nucleophile | 1 | 1 | 100.0 | 68.62 | inf | 1.46E-02 | 1.72E-01 | 1.00E+00 |
| GO:0000398 | mRNA splicing, via spliceosome | 1 | 1 | 100.0 | 68.62 | inf | 1.46E-02 | 1.72E-01 | 1.00E+00 |
| GO:0055114 | obsolete oxidation-reduction process | 188 | 7 | 3.7 | 2.56 | 2.91 | 1.75E-02 | 1.72E-01 | 1.00E+00 |
| GO:0006974 | cellular response to DNA damage stimulus | 40 | 3 | 7.5 | 5.15 | 5.8 | 1.96E-02 | 1.72E-01 | 1.00E+00 |
| GO:0033554 | cellular response to stress | 40 | 3 | 7.5 | 5.15 | 5.8 | 1.96E-02 | 1.72E-01 | 1.00E+00 |
| GO:0006099 | tricarboxylic acid cycle | 2 | 1 | 50.0 | 34.31 | 69.14 | 2.89E-02 | 1.72E-01 | 1.00E+00 |
| GO:0008380 | RNA splicing | 2 | 1 | 50.0 | 34.31 | 69.14 | 2.89E-02 | 1.72E-01 | 1.00E+00 |
| GO:0009060 | aerobic respiration | 2 | 1 | 50.0 | 34.31 | 69.14 | 2.89E-02 | 1.72E-01 | 1.00E+00 |
| GO:0015980 | energy derivation by oxidation of organic compounds | 2 | 1 | 50.0 | 34.31 | 69.14 | 2.89E-02 | 1.72E-01 | 1.00E+00 |
| GO:0045333 | cellular respiration | 2 | 1 | 50.0 | 34.31 | 69.14 | 2.89E-02 | 1.72E-01 | 1.00E+00 |
| GO:0006397 | mRNA processing | 3 | 1 | 33.3 | 22.87 | 34.56 | 4.31E-02 | 2.24E-01 | 1.00E+00 |
| GO:0016071 | mRNA metabolic process | 3 | 1 | 33.3 | 22.87 | 34.56 | 4.31E-02 | 2.24E-01 | 1.00E+00 |

**Supplementary Table 12**: Gene Ontology analysis of contracted genes in Parental 2 clones after culture growth.

| ID | Name | Bgd count | Result count | Pct of bgd | Fold enrichment | Odds ratio | P-value | Bonferroni | Benjamini |
|---|---|---|---|---|---|---|---|---|---|
| GO:0046364 | monosaccharide biosynthetic process | 23 | 23 | 100.0 | 3.45 | inf | 3.57E-13 | 9.21E-11 | 2.30E-11 |
| GO:0019319 | hexose biosynthetic process | 23 | 23 | 100.0 | 3.45 | inf | 3.57E-13 | 9.21E-11 | 2.30E-11 |
| GO:0006094 | gluconeogenesis | 23 | 23 | 100.0 | 3.45 | inf | 3.57E-13 | 9.21E-11 | 2.30E-11 |
| GO:0000077 | DNA damage checkpoint | 8 | 8 | 100.0 | 3.45 | inf | 4.91E-05 | 1.27E-02 | 7.92E-04 |
| GO:0000075 | cell cycle checkpoint | 8 | 8 | 100.0 | 3.45 | inf | 4.91E-05 | 1.27E-02 | 7.92E-04 |
| GO:0031570 | DNA integrity checkpoint | 8 | 8 | 100.0 | 3.45 | inf | 4.91E-05 | 1.27E-02 | 7.92E-04 |
| GO:0009058 | biosynthetic process | 407 | 145 | 35.6 | 1.23 | 1.42 | 1.14E-03 | 2.95E-01 | 1.74E-02 |
| GO:0008152 | metabolic process | 1336 | 487 | 36.5 | 1.26 | 1.88 | 1.51E-15 | 3.90E-13 | 3.90E-13 |
| GO:0006508 | proteolysis | 295 | 108 | 36.6 | 1.26 | 1.47 | 1.84E-03 | 4.75E-01 | 2.50E-02 |
| GO:0006520 | cellular amino acid metabolic process | 67 | 30 | 44.8 | 1.54 | 02.02 | 4.03E-03 | 1.00E+00 | 4.72E-02 |
| GO:0005975 | carbohydrate metabolic process | 50 | 23 | 46.0 | 1.59 | 2.11 | 7.58E-03 | 1.00E+00 | 8.50E-02 |
| GO:0022610 | biological adhesion | 145 | 67 | 46.2 | 1.59 | 2.19 | 5.24E-06 | 1.35E-03 | 1.13E-04 |
| GO:0007155 | cell adhesion | 145 | 67 | 46.2 | 1.59 | 2.19 | 5.24E-06 | 1.35E-03 | 1.13E-04 |
| GO:0055114 | obsolete oxidation-reduction process | 188 | 87 | 46.3 | 1.59 | 2.23 | 1.70E-07 | 4.39E-05 | 4.88E-06 |
| GO:0051726 | regulation of cell cycle | 15 | 8 | 53.3 | 1.84 | 2.81 | 4.11E-02 | 1.00E+00 | 4.24E-01 |
| GO:0044283 | small molecule biosynthetic process | 40 | 26 | 65.0 | 2.24 | 4.65 | 2.19E-06 | 5.66E-04 | 5.66E-05 |
| GO:0048519 | negative regulation of biological process | 13 | 9 | 69.2 | 2.39 | 5.55 | 3.07E-03 | 7.92E-01 | 3.77E-02 |
| GO:0048523 | negative regulation of cellular process | 13 | 9 | 69.2 | 2.39 | 5.55 | 3.07E-03 | 7.92E-01 | 3.77E-02 |

| | | | | | | | | | |
|---|---|---|---|---|---|---|---|---|---|
| GO:0042026 | protein refolding | 8 | 6 | 75.0 | 2.58 | 7.38 | 9.37E-03 | 1.00E+00 | 1.01E-01 |
| GO:0019318 | hexose metabolic process | 29 | 23 | 79.3 | 2.73 | 9.6 | 2.51E-08 | 6.47E-06 | 8.09E-07 |
| GO:0005996 | monosaccharide metabolic process | 29 | 23 | 79.3 | 2.73 | 9.6 | 2.51E-08 | 6.47E-06 | 8.09E-07 |
| GO:0045786 | negative regulation of cell cycle | 10 | 8 | 80.0 | 2.76 | 9.86 | 1.22E-03 | 3.16E-01 | 1.76E-02 |
| GO:0006006 | glucose metabolic process | 26 | 23 | 88.5 | 03.05 | 19.22 | 3.57E-10 | 9.20E-08 | 1.53E-08 |
| GO:0009072 | aromatic amino acid family metabolic process | 12 | 11 | 91.7 | 3.16 | 27.23 | 1.04E-05 | 2.68E-03 | 2.06E-04 |
| GO:0016051 | carbohydrate biosynthetic process | 24 | 23 | 95.8 | 3.3 | 57.72 | 6.23E-12 | 1.61E-09 | 3.21E-10 |

**Supplementary Table 13**: Number of non-synonymous SNPs in surface molecules genes (SM) and in other genes after culture growth.

| | Non-synonymous SNPs presents in the first generation | | | Non-synonymous SNPs after culture growth | | |
|---|---|---|---|---|---|---|
| | SNPs in SM genes | SNPs in other genes | Ratio SM/other genes | SNPs in SM genes | SNPs in other genes | Ratio SM/other genes |
| P1-1 | 3498 | 10826 | 0.32 | 866 | 1056 | 0.82 |
| P1-2 | 3424 | 10771 | 0.32 | 851 | 1058 | 0.80 |
| P1-3 | 3473 | 10824 | 0.32 | 980 | 1134 | 0.86 |
| P2-1 | 2466 | 9502 | 0.26 | 1935 | 2148 | 0.90 |
| P2-2 | 2526 | 9537 | 0.26 | 2039 | 2233 | 0.91 |
| P2-3 | 2410 | 9353 | 0.26 | 1902 | 2243 | 0.85 |
| 1C2-1 | 789 | 982 | 0.80 | 593 | 733 | 0.81 |
| 1C2-2 | 912 | 1224 | 0.75 | 595 | 785 | 0.76 |
| 1C2-3 | 846 | 1084 | 0.78 | 647 | 800 | 0.81 |
| 1D12-1 | 842 | 1180 | 0.71 | 543 | 698 | 0.78 |
| 1D12-2 | 288 | 439 | 0.66 | 215 | 254 | 0.85 |
| 1D12-3 | 977 | 1300 | 0.75 | 537 | 723 | 0.74 |
| 2C1-1 | 1154 | 1456 | 0.79 | 620 | 724 | 0.86 |
| 2C1-2 | 694 | 849 | 0.82 | 505 | 671 | 0.75 |
| 2C1-3 | 1057 | 1344 | 0.79 | 541 | 754 | 0.72 |
